# Centromere divergence and allopolyploidy reshape carnivorous sundew genomes

**DOI:** 10.1101/2025.07.27.666937

**Authors:** Laura Ávila Robledillo, Aaryan Bhatia, Jonathan Kirshner, Steven J. Fleck, Dirk Becker, Gerhard Bringmann, Jordan R. Brock, Daniela Drautz-Moses, Matthias Freund, Rainer Hedrich, Luis Herrera-Estrella, Enrique Ibarra-Laclette, Ines Kreuzer, Tianying Lan, Sachiko Masuda, Martín Mata-Rosas, Todd P. Michael, Héctor Montero, Sitaram Rajaraman, Michaela Richter, David Sankoff, Stephan C. Schuster, Ken Shirasu, Sonja Trebing, Yves Van de Peer, Gerd Vogg, Tan Qiao Wen, Yue Zhang, Chunfang Zheng, Kenji Fukushima, Jarkko Salojärvi, André Marques, Victor A. Albert

**Author notes:** Corresponding authors. (V.A.A.); (A.M.); (J.S.); (K.F.). First author (Laura Ávila Robledillo). These authors contributed equally (Aaryan Bhatia, Jonathan Kirshner, Steven J. Fleck).

## Abstract

Centromeres are essential for chromosome function, yet how contrasting centromere architectures shape polyploid genome evolution and plant diversification remains unresolved. In monocentric chromosomes, centromeric activity is restricted to one chromosomal region; in holocentric chromosomes, it is distributed along the chromosome length. Hybridization followed by genome doubling can place chromosome sets with different centromeres, repeat complements, and rearrangement histories in the same nucleus. We examined whether such genomic mergers are associated with later chromosome restructuring in carnivorous Caryophyllales, focusing on chromosome-scale assemblies of putatively holocentric *Drosera regia* and monocentric *D. capensis*. The two sundews retain broad syntenic continuity and related polyploid ancestry, but their present genomes differ sharply. *D. regia* preserves several ancestral chromosomal fusions and dispersed candidate centromere-associated satellite repeats, whereas *D. capensis* has stronger internal duplication, extensive rearrangement, and localized centromeres. Phylogenomic and structural analyses suggest a reticulate evolutionary history for *D. regia*, with part of its genome showing preferential structural affinity to the Venus flytrap lineage. Whereas *D. regia* exhibits threefold syntenic depth consistent with hexaploidy, *D. capensis* exhibits sixfold depth consistent with dodecaploidy, and structural comparisons do not support simple doubling of the extant *D. regia*-like architecture. Repeat profiling across a broad sample of *Drosera* also shows rapid turnover and species-level differences in centromere-associated DNA. These results place centromere organization alongside polyploid history and chromosome restructuring as a major axis of genome evolution in sundews.

## Introduction

In nutrient-poor habitats, flowering plants may alter how they acquire limiting elements. Carnivory is one such solution: it has evolved repeatedly, allowing plants to obtain nitrogen and phosphorus from captured animals^1,2^. The transition to carnivory involves changes in leaf development^3,4^ and the reuse of pathways involved in defense^5,6^, nutrient transport^7,8^, and secretion^8,9^. Several of these pathways are also associated with immunity and senescence^10–13^.

Carnivory has arisen at least ten times in five angiosperm orders^2^, producing snap traps, suction bladders, and pitfall pitchers among other forms. Pitcher traps alone evolved in three lineages separated by more than 100 million years^1,2^. Two especially species-rich carnivorous radiations occur in Lamiales and Caryophyllales^1,2^. While both of these clades include species sharing a convergent flypaper-trapping mode – butterworts (*Pinguicula*) and sundews (*Drosera*), respectively – they also contain carnivores with immensely divergent forms such as the suction trapping bladderworts (*Utricularia*) and the snap-trapping Venus flytrap (*Dionaea*).

Within Caryophyllales, *Drosera* includes roughly 250 species^14^, most of which capture prey with adhesive, mucilage-secreting tentacles^15^. The basic flypaper trap is conserved, but morphology and ecology vary widely across the genus and its close relatives. These close relatives include the snap-trapping Venus flytrap and water-wheel plant, *Dionaea* and *Aldrovanda*, the pitcher plant *Nepenthes*, the morphologically divergent flypaper trappers *Triphyophyllum*^16–18^ and *Drosophyllum*^19,20^, and three genera of secondarily non-carnivorous vines (*Ancistrocladus*, *Habropetalum*, *Dioncophyllum*)^16^.

Sundews are also remarkable for variation in chromosomal and genomic architecture^21–24^. Cytogenetic studies have reported both monocentric chromosomes, with a single localized centromere, and holocentric chromosomes, where centromeric activity is distributed along the entire chromosome length^21,23,24^. However, small chromosome size and a lack of genome-scale data have left sundew centromere structure unresolved.

Across eukaryotes, centromere organization influences chromosome-scale evolutionary dynamics beyond its immediate role in genome architecture^25–27^. Repeat composition, transposable element (TE) distribution, histone modifications, and recombination patterns all shape large-scale chromosomal behavior^25–28^. In holocentric species, chromosome fissions and fusions can occur without loss of centromere function, potentially accelerating structural evolution^25–27,29–31^. Despite this potential significance, the evolutionary roles of holocentricity in flowering plants remain poorly understood.

Recent comparative genomic studies of carnivorous plants have already shown that extensive gene family remodeling, lineage-specific innovation, and sometimes repeated genome duplication have accompanied the evolution of carnivory in lineages such as *Cephalotus*^10^, *Utricularia*^32^, and *Nepenthes*^13^. Carnivorous Caryophyllales also show repeated paleopolyploidy, extensive gene-tree conflict, and rapid chromosome-number evolution^33,34^.

These observations make polyploid history central to interpreting chromosome evolution in the group. Polyploidy has also been associated more broadly with ecological expansion, stress tolerance, evolutionary persistence, and morphological or physiological novelty^35–38^. After WGD, plant genomes can develop differentiated subgenomes, biased gene retention, rearrangements, and new repeat distributions. Some consequences may buffer structural change through redundancy (as proposed for the carnivore *Utricularia*^32^), whereas multivalent formation, homoeologous recombination, aneuploid gametes, and reduced fertility can disrupt meiosis before diploidized inheritance is established^39,40^. The relevant issue here is therefore what happens to duplicated genomes after multiplication, not simply whether polyploidy occurred. In angiosperms, there is no general expectation that polyploidy should favor either monocentric or holocentric chromosomes. Even so, combining divergent chromosome sets creates an opportunity for centromere features to change during stabilization, and such changes have been documented in allopolyploid wheat and cotton^41,42^.

The carnivorous Caryophyllales bring several of these variables together, since centromere organization, chromosome structure, phylogenetic discordance, and polyploid history of unknown nature characterize the group. If allopolyploid origins underlie some of its lineages, the group’s phylogeny cannot be represented by a single bifurcating species tree. Gene-tree conflict and repeated duplication or hybridization must be read together with synteny, Ks distributions, and chromosome structure to provide a more complete summary of genome history^33,34^. *D. regia* is especially interesting in this regard because it is morphologically distinctive, cytogenetically unusual, and falls outside the core sampled *Drosera* lineage in some phylogenetic analyses. Its position also bears on the older question of whether the species should be separated taxonomically from *Drosera*. The point of the present comparison is therefore not polyploidy alone, but the conjunction of contrasting centromere organization, divergent post-polyploid restructuring, and evidence for a mosaic genome history in *D. regia*.

Here, we address the complex evolutionary trajectories within carnivorous Caryophyllales focusing on *D. regia* and *D. capensis*, phylogenetically and morphologically distinct sundews that differ substantially in reported centromere structure and ploidy level. To place these differences in broader context, we also survey DNA repeat content across *Drosera* to assess its contribution to centromere diversity within the genus. We also use phylogenetic and genomic evidence across the broader carnivorous Caryophyllales lineage, including newly sequenced genomes of *Triphyophyllum*, *Ancistrocladus*, and *Dionaea*^43–46^, to ask whether deeply divergent clades contributed to *Drosera* paleohistory via ancient admixture, and how such reticulation may relate to centromere and chromosome change. Our analyses reveal sharply contrasting centromere and post-polyploid genome architectures in the two sundews, together with evidence that *D. regia* has a mosaic evolutionary history that includes subgenome-specific structural affinity to a *Dionaea*-like lineage. The comparison therefore places contrasting centromere architectures in the context of the polyploid and structural histories reconstructed for the same genomes.

## Chromosome-scale sundew genomes

To enable comparisons of repeat organization, synteny, and chromosome architecture between our focal sundew species, we generated chromosome-scale assemblies for *D. capensis* and *D. regia* using PacBio SMRT sequencing, Illumina sequencing, and Hi-C chromosome-conformation capture (**Supplementary Materials Table S1**). The final *D. capensis* assembly spans 284,038,097 bp, with a contig N50 of approximately 1.0 Mb, a scaffold N50 of 12.65 Mb, and 94.8% complete embryophyta_odb10 BUSCOs. The *D. regia* assembly spans 282,006,236 bp, with a contig N50 of 287 kb, a scaffold N50 of 15.76 Mb, and 96.7% complete embryophyta_odb10 BUSCOs. Against Earth BioGenome Project reference benchmarks (https://www.earthbiogenome.org/report-on-assembly-standards), *D. capensis* reaches approximately megabase-scale contig continuity; both assemblies have chromosome-scale scaffolds and recover most conserved embryophyte genes.

Genome-size estimates add an important qualification. The *D. capensis* assembly span is close to both the KmerGenie estimate of 286.1 Mb and an earlier published flow-cytometric 1C estimate of 293.5 Mb, but it is substantially below the 1C estimate of 394.5 Mb calculated from the measurement reported by Veleba et al.^47^ The expected genome size of *D. capensis*, and therefore its completeness by total span, consequently remains uncertain. By contrast, the *D. regia* assembly span is below both its KmerGenie estimate of 348.7 Mb and its flow-cytometric 1C estimate of 417.5 Mb, suggesting that repetitive or otherwise difficult-to-assemble sequence is underrepresented, consistent with its lower PacBio coverage and lower contig-level continuity (**Supplementary Materials Table S1**; **Supplementary Materials Fig. S10**).

The number of large pseudochromosomal scaffolds in the *D. capensis* and *D. regia* assemblies matched their known haploid chromosome numbers, n = 20 and n = 17, respectively^33^ (**Fig. 1**). Our chromosome-level assemblies were annotated using RNA-seq data from *D. capensis* and other species (see Methods). We annotated 32,230 and 32,090 high-confidence gene models along the 17 and 20 pseudomolecules of *D. regia* and *D. capensis*, respectively.

**Fig. 1:**
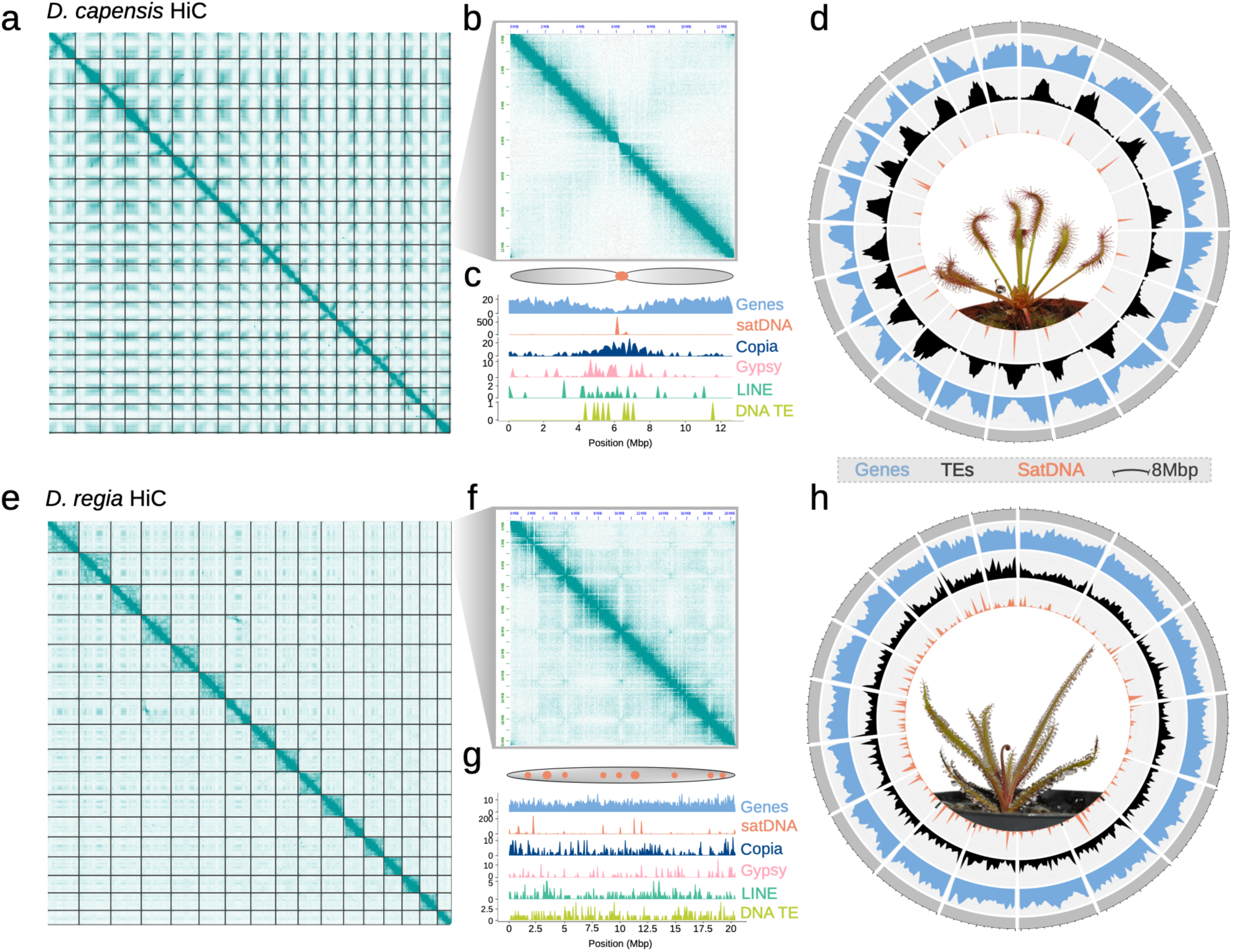
Contrasting spatial genome organization in monocentric *D. capensis* and putatively holocentric *D. regia* chromosomes. (**a, e**) Genome-wide Hi-C contact maps and (**b, f**) single-chromosome views show the distribution of major repetitive sequence types (**c, g**), clustered in *D. capensis* and uniform in *D. regia*, consistent with localized and chromosome-wide centromere organization, respectively. Sequence-type density was calculated in 100 kb windows. (**d, h**) Circos plots of gene, TE, and satDNA distributions in *D. capensis* (**d**) and *D. regia* (**h**) further illustrate genomic patterns consistent with localized versus chromosome-wide centromere organization.

## Chromosome organization diverges between two *Drosera* genomes

Several species of *Drosera* have been reported to have holocentric chromosomes with centromeric foci distributed along the length of the chromosome^21–24^. We combined Hi-C contact maps with genome-wide repeat annotations to infer the centromeric organization of the *D. capensis* and *D. regia* chromosomes (**Fig. 1a-h**). The *D. capensis* Hi-C map displayed a classic monocentric architecture, showing distinct A (euchromatin) and B (heterochromatin) compartments, including some degree of a telomere-to-centromere interaction axis (**Fig. 1a-b**). Correspondingly, repeat profiling of *D. capensis* revealed a single, size-restricted region of satellite DNA (satDNA) probably marking the centromere, flanked by a high density of transposable elements in the pericentromeric region, tapering toward the telomeres, while genes were concentrated on the distal chromosome arms (**Fig. 1c-d**). In contrast, the *D. regia* chromosomes lacked large-scale compartmentalization and showed no telomere-to-centromere interaction axis in their Hi-C maps (**Fig. 1e-f**), consistent with a non-monocentric structure. The *D. regia* repeat landscape mirrored this difference: satDNA occurred as multiple short arrays dispersed along the chromosomes, and both transposable elements and genes were uniformly distributed, without the typical pericentromeric enrichment (**Fig. 1g-h**).

These results are in close agreement with recent reports for other holocentric plants. As recently argued by Hofstatter, et al.^25^ and Mata-Sucre, et al.^27^, the concept of chromosome arms does not apply to holocentric species. Hi-C contact patterns and chromosome-wide repeat distributions are consistent with holocentricity, although neither directly identifies functional kinetochore sites. At present, our inference of holocentricity in *D. regia* is not matched by other previously-assembled genomes of carnivorous Caryophyllales under study; for example, the Hi-C contact map of *Dionaea muscipula* reflected a chromosomal architecture comparable to that of *D*. *capensis*, in contrast to previous non-genomic research that suggested holocentricity for *Dionaea muscipula*^22^ (**Supplementary Materials Fig. S1**).

## Cytological and satellite evidence for contrasting centromere organization

We examined the genomes of *D. regia* and *D. capensis* to identify CENH3, a centromere-specific variant of histone H3 that plays an essential role in kinetochore assembly and is notably divergent across species^25^. In both species, two copies of CENH3 were identified, and antibodies were generated against their variable N-terminal tails (**Supplementary Materials Fig. S2a**). Immunostaining using these CENH3 antibodies produced distinct foci in the interphase nuclei of *D. capensis*, confirming monocentric chromosome organization (**Fig. 2a**). In contrast, no specific signal was observed in interphase nuclei of *D. regia* (**Fig. 2b**).

**Fig. 2:**
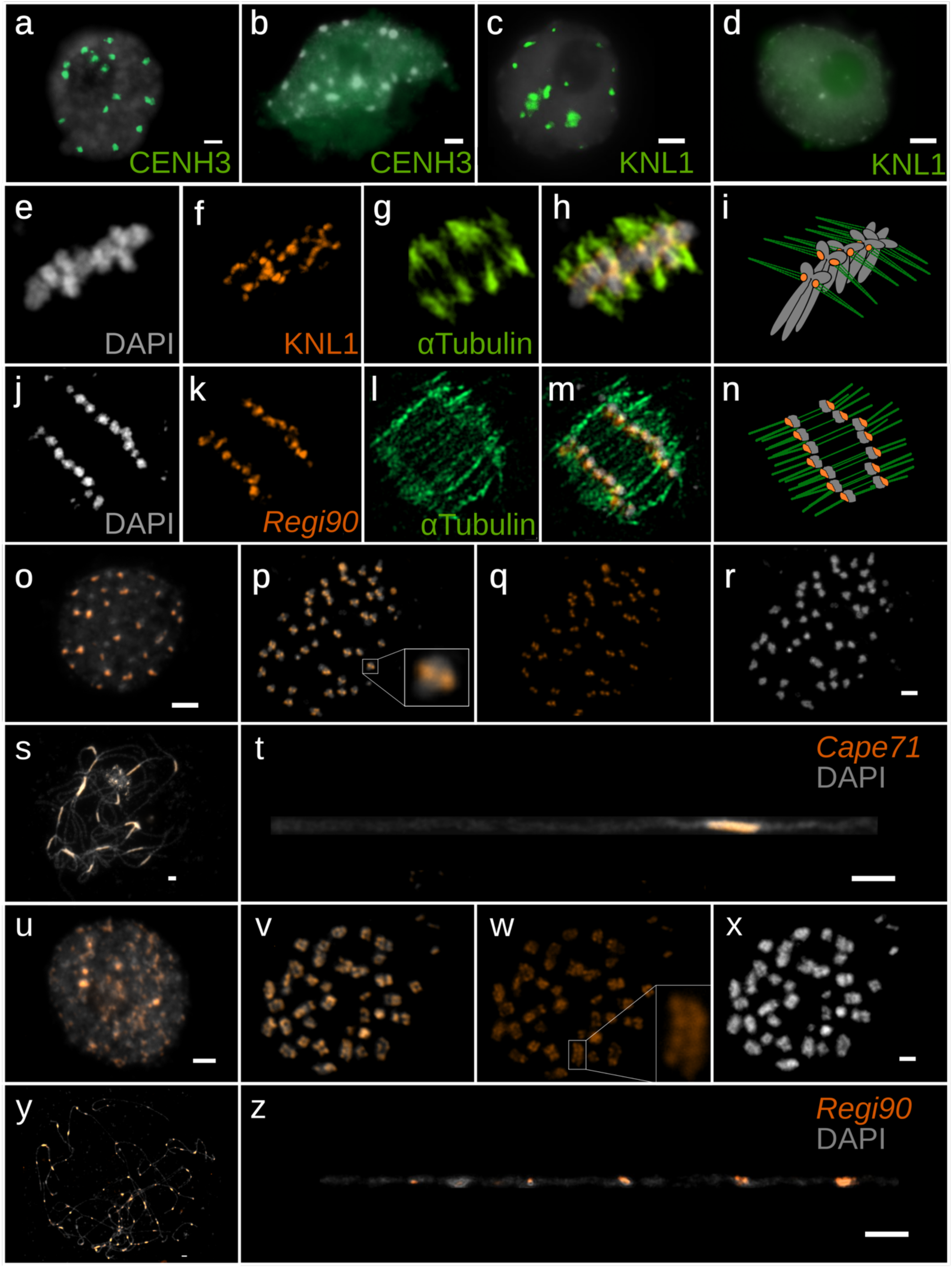
Centromere morphologies in *D. capensis* and *D. regia* revealed by immunostaining and FISH. Immunostaining of interphase nuclei with CENH3 antibodies shows localized signals in *D. capensis* (**a**) and non-specific signals in *D. regia* (**b**). Similar patterns are observed with KNL1 antibodies in *D. capensis* (**c**) and *D. regia* (**d**). (**e-h**) Immunostaining of *D. capensis* metaphase chromosomes with KNL1 and α-tubulin reveals localized kinetochore/microtubule attachment regions. (**i**) Model summarizing *D. capensis* observations. (**j-m**) Combined α-tubulin immunostaining and FISH for *Regi90*, the dominant 90-bp satellite repeat in *D. regia*, show α-tubulin signal along the metaphase chromosome length together with chromosome-wide *Regi90* signals; these images indicate spatial proximity but do not resolve direct molecular colocalization. (**n**) Model summarizing *D. regia* observations. (**o-t**) FISH with the *Cape71* probe in *D. capensis* shows clustered centromeric signals. (**u-z**) In *D. regia*, *Regi90* satellite signals are distributed along the chromosomes. Panels illustrate contrasting patterns in interphase nuclei (**o, u**), metaphase chromosomes (**p-r, v-x**), and pachytene chromosomes (**s-t, y-z**).

Given that several holocentric lineages do not require CENH3 for centromere function^48^, we investigated the presence of the kinetochore component KNL1, a protein conserved across many plant lineages^49^. In *D. capensis*, KNL1 antibodies revealed distinct nuclear foci at interphase (**Fig. 2c**), consistent with localized centromeric activity. However, no specific KNL1 signal could be detected in *D. regia* interphase nuclei (**Fig. 2d**). RNA-seq evidence for genes encoding several kinetochore-related proteins (*Supplementary Note 1*) nevertheless supports the presence and transcription of *KNL1* and *NDC80* in both taxa, as well as one clearly intact and expressed *CENH3* paralog in each species, with fragmented *KNL1* transcript recovery in *D. regia* arguing against a simple interpretation based on wholesale gene loss. Metaphase chromosomes of *D. capensis* stained with KNL1 displayed signals confined to discrete chromosomal regions, colocalizing with α-tubulin, further supporting monocentricity (**Fig. 2e-i**). In *D. regia*, although KNL1 signals were absent, α-tubulin was observed to associate along the entire length of the mitotic chromosomes (**Fig. 2j-m**; summarized in **Fig. 2n**). These results provide clear evidence for monocentricity in *D. capensis*, and support a chromosome-wide, non-monocentric organization in *D. regia*. Because the *D. regia* chromosomes are small and these preparations have limited spatial resolution, the cytology cannot distinguish a continuous holocentric state from a chromosome-wide set of discrete centromeric domains. We therefore describe *D. regia* as putatively holocentric, while recognizing that a punctate, chromosome-wide (polycentric) centromere organization cannot be excluded. Failure to detect KNL1 or CENH3 in *D. regia* could also reflect antibody or fixation effects rather than absence of kinetochore localization. Our inference rests on the combination of Hi-C contact structure, dispersed satellite FISH, α-tubulin attachment along metaphase chromosomes, and genome-wide repeat distribution. Together, these features are consistent with a chromosome-wide centromere organization in *D. regia*.

To complement the cytological and protein-localization results, we next examined the distribution of satellite DNA (satDNA) repeats, which, although not universally required for centromere function, are prominent features of many plant centromeres, especially in angiosperms^50–52^. Each species exhibited a dominant satDNA family: *Cape71* in *D. capensis*, consisting of a 71 base pair monomer tandem repeat, and *Regi90* in *D. regia*, a 90 base pair tandem repeat. We treat *Regi90* as a candidate centromere-associated repeat; its FISH signal reports the chromosomal distribution of the repeat rather than direct kinetochore localization. The *Cape71* and *Regi90* sequences showed no significant similarity to each other (**Supplementary Materials Fig. S2b**), hinting that centromere evolution more broadly in *Drosera* could be more dynamic than centromere architecture alone would suggest.

Fluorescence in situ hybridization (FISH) using a probe for *Cape71* revealed concentrated centromeric signals in *D. capensis* during interphase and metaphase. At pachytene, *Cape71* formed one concentrated cluster per *D. capensis* chromosome (**Fig. 2o-t**), consistent with the same localized pattern. *Regi90* behaved differently in *D. regia*: interphase nuclei contained many dispersed foci (**Fig. 2u**), metaphase chromosomes carried line-like signals along their length (**Fig. 2v-x**), and pachytene chromosomes showed multiple foci (**Fig. 2y-z**), as reported for other holocentric plants^53,54^. α-tubulin and *Regi90* signals occupied the same broad chromosome regions in combined preparations, although the resolution does not permit a claim of molecular colocalization. Genome sequence places *Regi90* in short arrays, usually about 10 to 20 kilobases, across all *D. regia* chromosomes (**Supplementary Materials Fig. S2c**). The distribution resembles repeat-based holocentromeres described in *Rhynchospora*^54^, although direct CENH3 or KNL1 localization has not been demonstrated in *D. regia*.

## Contrasting satellite DNA evolution in two sundews

To examine how the dominant satellite repeat families evolved in *D. regia* and *D. capensis*, we constructed phylogenetic trees based on sequences from the *Regi90* and *Cape71* families. Separate alignments for the two repeat families contained abundant SNP and indel variation. We use this sequence divergence to examine relationships among repeat copies, recognizing that differences in divergence do not directly establish when duplication or movement occurred. When monomers were labeled by chromosome and genomic position, the trees contained both chromosome-restricted clades and clades mixing several chromosomes (**Fig. 3**). One interpretation of this pattern is that local amplification was followed, at least in some repeat lineages, by dispersal to additional genomic sites.

**Fig. 3:**
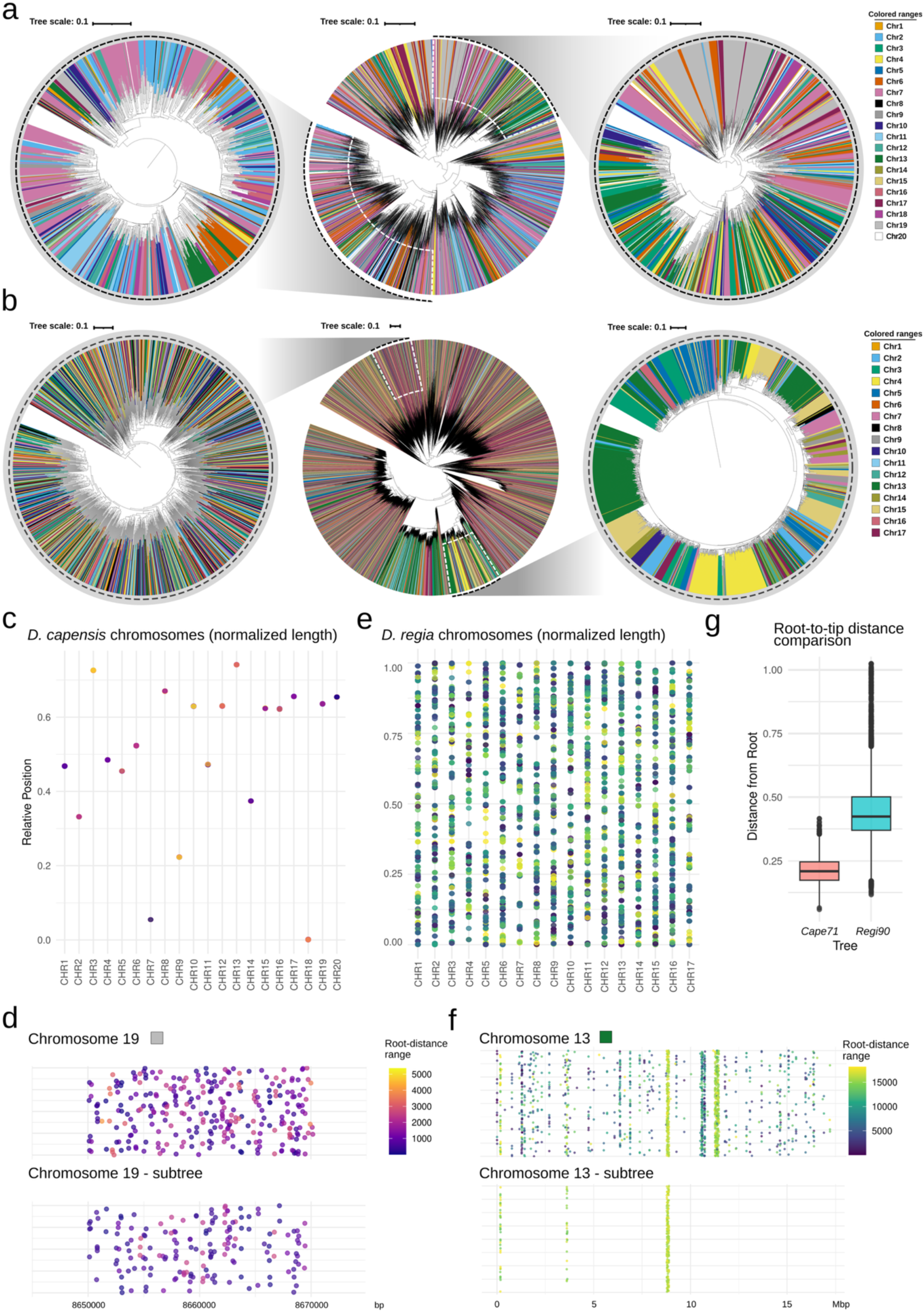
Satellite DNA monomer phylogenies and chromosomal distributions in *D. capensis* and *D. regia*. (a,b) *Cape71* monomer tree for *D. capensis* **(a)** and *Regi90* monomer tree for *D. regia* **(b)**. The radial tree in each panel contains all monomers; flanking trees enlarge the indicated sectors, with dotted lines linking each enlargement to its position in the complete tree. Tip color gives chromosome of origin. We use “chromosome-specific” for clades dominated by monomers from one chromosome, irrespective of the blue highlighting. **(c)** *Cape71* monomers plotted at normalized positions along *D. capensis* chromosomes; dot color is ranked midpoint-root-to-tip distance within the *Cape71* tree. **(d)** Chromosome 19 detail from **(c)**, with all mapped monomers above and members of the selected subtree below. **(e)** *Regi90* monomers plotted in the same way for *D. regia*, with distances ranked within the *Regi90* tree. **(f)** Chromosome 13 detail from **(e)**. **(g)** Unranked root-to-tip distances from the two monomer trees. Vertical jitter in **(d)** and **(f)** is used only to separate overlapping points.

The *Cape71* phylogeny (**Fig. 3a**) includes multiple chromosome-specific clades with long branches, indicating that monomer subfamilies have arisen independently and undergone sustained amplification in a chromosome-restricted manner. In contrast, the *Regi90* tree (**Fig. 3b**) shows a starburst-like topology with short internal branches and widespread intermixing of terminal tips from different chromosomes. This pattern is consistent with episodic amplification over extended timescales, with ongoing local expansions superimposed on existing *Regi90* diversity. Few same-chromosome clades are present in *D. regia*, suggesting regionally distributed, nascent amplifications.

These differences suggest that *D. capensis* experiences stronger and more sustained local concerted evolution. In this species, repeat variants that arise on a chromosome tend to remain and amplify there, maintaining their identity over time. *D. regia*, by contrast, exhibits greater sequence divergence and inter-chromosomal mixing of repeats, with fewer chromosome-restricted subfamilies.

To more directly assess chromosome-specific structure, we labeled tree tips by chromosome of origin and applied midpoint-rooting to visualize relative branch lengths. In *D. capensis* (**Fig. 3c**), each point represents a *Cape71* monomer, plotted at the midpoint of its genomic interval, normalized by chromosome length on the vertical axis, and colored by root-to-tip distance (phylogenetic divergence in the monomer tree). The color of each dot within each vertical band indicates that centromeric regions on individual chromosomes house repeats with different levels of sequence divergence. These divergent subfamilies are consistent with amplification of distinct repeat lineages on individual chromosomes.

Zooming in on chromosome 19 (**Fig. 3d**), a narrow gradient in root-distance values is visible across the centromeric region. This gradient persists across the whole centromeric region even when the analysis is restricted to monomers from a particular *Cape71* subtree. A nearly monochromatic cluster within this ∼20 kb region highlights a group of closely related monomers (i.e., homogenized sequences).

In *D. regia*, the *Regi90* distribution plot (**Fig. 3e**) shows a continuous scatter of monomers along chromosomes, with color variation spanning the full root-to-tip distance range, indicating extensive mixing of monomers with different divergence levels across the genome. On chromosome 13 (**Fig. 3f**), monomers span the full color gradient and are broadly distributed across the chromosome. Within the selected **Fig. 3b** subtree, however, a subset of chromosome 13 monomers occupies a much narrower range of root-to-tip distances. That local concentration is compatible with recent amplification of a related *Regi90* subset, including the possibility of polymeric organization, but it does not distinguish among mechanisms.

*Regi90* and *Cape71* differ in both divergence and chromosome specificity. *Regi90* has longer root-to-tip paths (**Fig. 3g**), but its monomers are more thoroughly mixed among chromosomes. On the other hand, *Cape71* is less divergent overall and forms more chromosome-restricted, internally similar clades. Pairwise sequence comparisons provide the same qualitative contrast (**Supplementary Materials Fig. S3**). *Cape71* arrays are relatively more homogeneous within and between *D. capensis* chromosomes (**Supplementary Materials Fig. S3a**), whereas *Regi90* arrays in *D. regia* average only about 75 percent identity in both intra– and inter-chromosomal comparisons (**Supplementary Materials Fig. S3c**).

Chromosome-by-chromosome summaries agree with this result: median root-to-tip distance is higher for *Regi90* across *D. regia* pseudochromosomes than for *Cape71* across *D. capensis* pseudochromosomes (**Supplementary Materials Fig. S4**). Together with the independent ModDot sequence-identity comparisons, these phylogenetic results support greater sequence divergence and weaker homogenization of *Regi90*, with the differences evident across chromosomes rather than confined to a single chromosome.

These patterns do not point to a single mechanism. The chromosome-restricted *Cape71* clades could arise mainly through local amplification within arrays, including unequal sister chromatid exchange or break-induced replication, with relatively little exchange among chromosomes. *Regi90* shows the opposite combination, with high monomer divergence and extensive interchromosomal mixing. This pattern is compatible with repeated movement among chromosomes, for example through ectopic recombination or reintegration of extrachromosomal repeat circles, together with local expansions, although these mechanisms remain hypothetical.

One confounding factor in *D. capensis* is the large amount of satDNA located on small, unplaced scaffolds. Of the 17 megabases of *Cape71* identified by TideCluster, only 1.12 megabases were assigned to chromosomes, possibly reflecting assembly challenges due to high sequence similarity. In *D. regia*, 3.8 megabases of *Regi90* were annotated, with 3.5 megabases placed into pseudochromosomes. Consequently, chromosome-level comparisons sample a substantially smaller proportion of *Cape71* than *Regi90*, potentially affecting estimates of sequence diversity and homogenization.

Collectively, these data reveal distinct evolutionary trajectories for the dominant satellite DNA families in sundews. In monocentric *D. capensis*, *Cape71* arrays show stronger local identity and homogenization. In putatively holocentric *D. regia*, *Regi90* exhibits greater sequence divergence and broader genomic dispersion. These contrasting patterns are consistent with a relationship between chromosome-wide centromere organization and satellite DNA evolutionary dynamics, while not demonstrating that *Regi90* itself is a functional centromere repeat.

## High repeat turnover in sundews

Among the most abundant transposable elements, the genomes of *D. regia* and *D. capensis* were generally rich in *Copia* LTR retrotransposons^55^, with a special dominance of *SIRE* in *D. capensis* and *TORK* in *D. regia* **(Supplementary Materials Fig. S5a)**. Although the majority of TEs in *D. capensis* were found in pericentromeric regions, a fraction of *TORK* elements was present in proximity to genes at a greater frequency than other TEs **(Supplementary Materials Fig. S5b)**, similar to the patterns observed in *D. regia* **(Supplementary Materials Fig. S5b, right panels)**. However, comparison of the distribution of different TEs around *Cape71* and *Regi90* satellite loci in relation to the putative centromere positions revealed a preference for LINEs in *D. capensis* **(Supplementary Materials Fig. S5b, upper left)** and for the chromovirus *REINA* in *D. regia* **(Supplementary Materials Fig. S5b, upper right)**. While chromoviruses preferentially target centromeric regions, LINE elements are often found pericentromerically^56,57^. These TE associations characterize the genomic environments of Cape71 and Regi90 rather than independently establishing their centromeric functions. Together with the cytological evidence, these repeat and TE distributions are consistent with *Cape71* and *Regi90* being centromere-associated candidates. For *Regi90*, however, direct association with a functional kinetochore has not yet been shown.

Next, we aimed to characterize the dynamics and abundance of repetitive DNA across the *Drosera* genus. Although the specific composition of such DNAs is highly variable among angiosperm species, these sequences have proven to be major players in the evolution of eukaryotes^58^. For this purpose we included Illumina reads from an additional 10 *Drosera* species to analyze their repetitive fractions (**Supplementary Materials Table S2**). Across a phylogenetically broad but taxonomically limited sample of Drosera, comparative repeatome profiling reveals marked species-level differences in repetitive DNA composition, including satellite repeat families (**Supplementary Materials Fig. S5c,d**). The sampled taxa span several major lineages, including *D. regia*, African and temperate representatives of *Drosera* s.s., Australian tuberous sundews, pygmy sundews, and the woolly/petiolaris complex. However, because several species-rich or phylogenetically distinctive lineages remain unsampled, these patterns should be interpreted as trends across the sampled diversity rather than as a complete genus-wide survey. These similarities apparently reflect their phylogenetic relationships, as the sister species *D. capensis* and *D. aliciae* share a great diversity of repetitive elements. Apart from some families of transposable elements, such as ATHILA, SIRE or LINE, the satellite repeat *Cape71* in *D. capensis* was found to be shared with *D. aliciae* (**Supplementary Materials Fig. S5c**). In addition, a closer look into the similarities between *Cape71* and the remaining satellite repeats found among the 12 species revealed that *Cape71* is related to both of the satDNA clusters found in *D. spatulata* (**Supplementary Materials Fig. S5d**), thus reflecting the close evolutionary relationships of the three species. Another case of repeatome interrelationship that matches phylogeny was observed for *D. anglica*, *D. filiformis* and *D. intermedia*, where most of the different families of transposable elements were shared between these species but not with the other *Drosera* species studied. However, in this case, no shared satDNA repeats were found. Finally, great similarities among the elements of *D. erythrorhiza* and *D. peltata* were also found. Despite their different bursts of the RETAND and LINE families, these species shared most of their elements with their sister species *D. menziesii* (**Supplementary Materials Fig. S5c**). In addition, when the satDNA fractions of these three species were compared, we observed that the three taxa shared the same family of satDNA (**Supplementary Materials Fig. S5d**).

Within the limited set of taxa analyzed here, the putatively holocentric lineages included (*D. regia*, *D. scorpioides*, *D. broomensis*) show a comparatively high proportion of lineage-specific repeat clusters, including satDNAs (**Supplementary Materials Fig. S5c**). In this regard, *Regi90* does not share significant similarity with any of the other satellites analyzed, except for short stretches of homopolymer/mononucleotide repeats (AAA)n (**Supplementary Materials Fig. S2b, S5d**). Nevertheless, within the monocentric lineages represented here we observe phylogeny-consistent satDNA similarity, including *Cape71* sharing between *D. capensis* and *D. aliciae* and its relationship to the two satDNA clusters identified in *D. spatulata* (**Supplementary Materials Fig. S5d**), whereas the putatively holocentric taxa included are dominated by lineage-specific satDNA clusters (**Supplementary Materials Fig. S5c-d**). Broader taxon sampling will be required to test whether this trend generalizes across *Drosera*. Altogether, these results reveal marked lineage-specific variation in repetitive DNA that may reflect differences in centromere organization, phylogenetic history, or both.

In summary, patterns of centromere organization and repeat distribution in *D. regia*, *D. capensis* and the other species examined indicate that these features are not static and may be reshaped during major genomic events. Repeat turnover, TE activation, and centromere repositioning can accompany or follow hybridization, especially when divergent parental genomes merge. Such events often coincide with whole-genome duplication, producing a “genomic shock”^59,60^ that destabilizes prior chromosomal configurations. The order of events nevertheless remains unresolved: holocentricity could have made chromosome fissions or fusions easier to retain, or it could have arisen after major karyotypic change and helped stabilize the resulting chromosomes^61^. The rarity of holocentricity among angiosperms argues against assuming either route is general. Broader comparisons within *Drosera* will be needed to determine which sequence of events applies in the genus. The present data are consistent with the possibility that chromosome-wide kinetochore activity could preserve segregation after major structural changes. Such buffering may not eliminate the costs of missegregation, altered recombination, or reduced fertility, but it could affect which hybrid or polyploid karyotypes survive long enough to stabilize and persist.

## Phylogeny and genome structural evolution of the Caryophyllales carnivore clade

Because the centromere and structural differences between *D. regia* and *D. capensis* must be interpreted in an evolutionary context, we next examined phylogenetic relationships, gene-tree conflict, and syntenic evidence for polyploidy across carnivorous Caryophyllales. This framework allows the contrasting chromosome architectures of the two sundews to be evaluated in light of broader patterns of hybridization, genome multiplication, and post-polyploid restructuring in the clade.

### Backbone phylogeny reveals unexpected relationships for D. regia and Nepenthes

The genome of *Nepenthes* revealed a complex allopolyploid history with five subgenomes of *x*=8^13^, which may complicate inference of relationships within *Drosera*. To address this, we generated backbone phylogenies for Caryophyllales carnivores using conserved Benchmarking Universal Single-Copy Ortholog (BUSCO)^62^ gene from *Drosera*, its close relatives, and selected outgroup angiosperms. Two datasets were analyzed: nucleotide and inferred amino acid sequences for 22 and 50 species, respectively (**Supplementary Materials Fig. S6**). The different taxon samplings were used to test the stability of the inferred relationships under different outgroup compositions. Phylogenetic reconstructions were performed using two methods: maximum likelihood (ML) analyses^63^ of concatenated BUSCO alignments, and ASTRAL species tree reconstruction^64^ summarizing individual BUSCO gene trees. Across all four reconstructions, *D. regia* grouped with *Aldrovanda* plus *Dionaea* rather than with the other sampled *Drosera*. The placement of *Nepenthes*, however, varied among datasets and phylogenetic methods. Some reconstructions resolved *Nepenthes* as sister to *Drosera* plus the snap-trapping lineages, whereas the protein-based ASTRAL analysis placed *Nepenthes* as sister to the clade comprising *Drosophyllum*, *Triphyophyllum*, and *Ancistrocladus* (**Supplementary Materials Fig. S6**). This latter relationship is also consistent with a prominent resolution represented in the gene-tree cloudogram (Fig. 4) and with an earlier BUSCO-based analysis^13^. This difference indicates that a single bifurcating topology may not fully describe relationships among these polyploid genomes. ^65^.

**Fig. 4:**
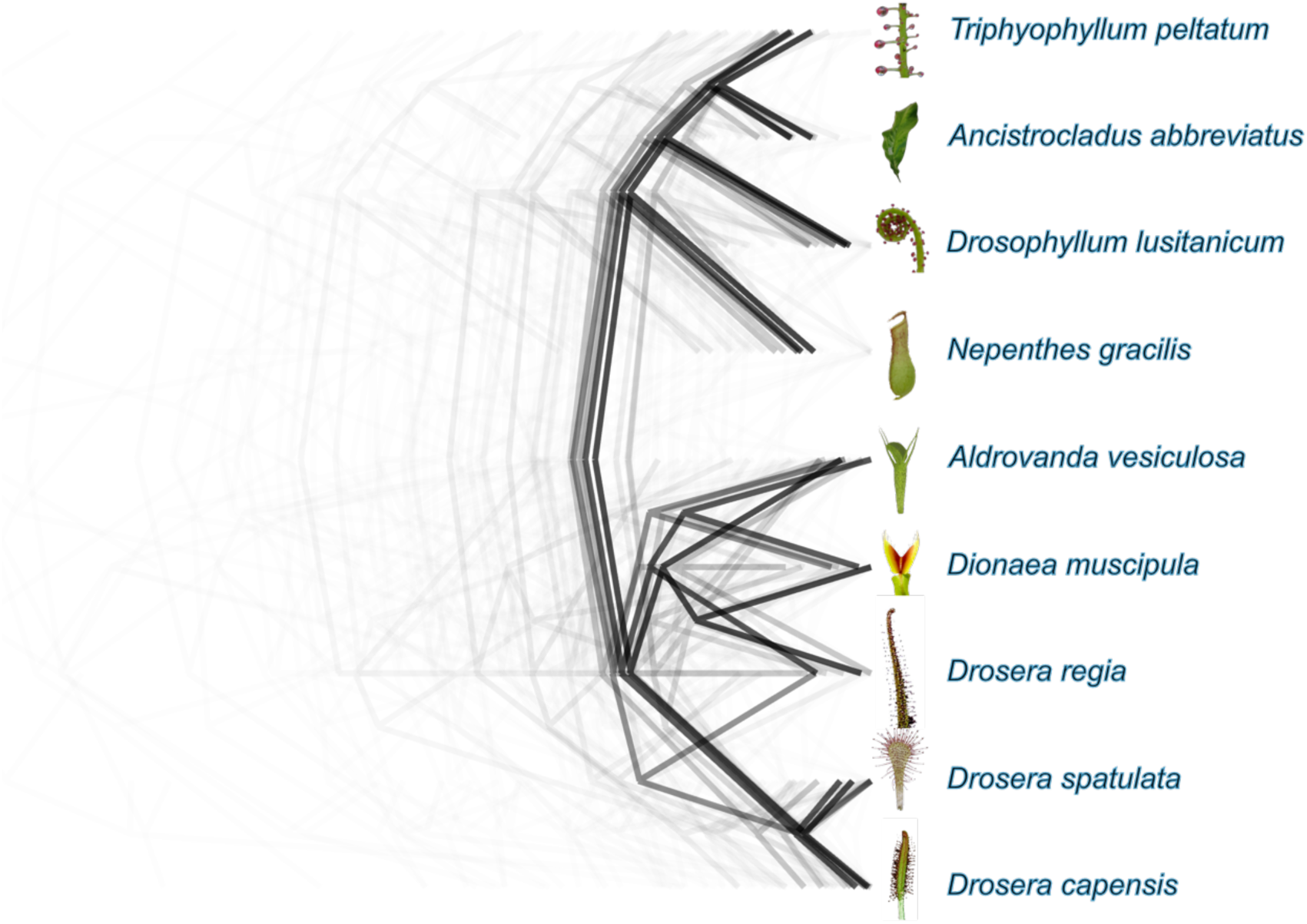
DensiTree cloudogram of 563 single-copy ortholog phylogenies. Each line is one undated maximum-likelihood tree for the same focal taxon set. Trees were rooted with *Ancistrocladus abbreviatus*, plotted in the same tip order, and left at their inferred branch lengths; the DensiTree consensus is superimposed and darker paths mark topologies followed by more trees. Branch depth therefore varies among loci and is not a shared time scale. Alternative placements of *D. regia*, *Dionaea*, and *Aldrovanda* show the extent of gene-tree discordance. The cloudogram is descriptive and does not separate hybridization from incomplete lineage sorting.

### Single copy genes display substantial gene tree conflict

To examine gene-tree conflict within the group (as anticipated from earlier results^34^), we analyzed single-copy orthologs from eight carnivorous Caryophyllales species and the non-carnivorous relative *Ancistrocladus abbreviatus*. A DensiTree cloudogram^66^ of 563 maximum-likelihood gene trees containing the complete focal taxon set was used to visualize variation in gene-tree topology and branch length (**Fig. 4**). The individual gene trees were not time-calibrated, but were rooted on the *Ancistrocladus* branch for consistent visualization and displayed with a common tip order. The analysis revealed substantial conflict in the placement of *D. regia*, with some loci grouping it with *Aldrovanda* and *Dionaea* and others grouping it with the remaining sampled *Drosera* species. Relationships involving *Aldrovanda* and *Dionaea* were also heterogeneous among gene trees. The cloudogram is used here only to show that loci carry different histories. Both hybridization and incomplete lineage sorting could produce this pattern, and the plot does not distinguish between them. Differences in branch depth are likewise not interpreted as divergence times.

### Genome structure suggests complex polyploidy

To delve further into these relationships, gene-based synteny analyses were performed using GENESPACE^67^. Genomes of *D. capensis* and *D. regia* were compared to those of closely related outgroups, including *Nepenthes*, *Dionaea muscipula*^43^, *Ancistrocladus abbreviatus*, and *Triphyophyllum peltatum*. The *N. gracilis* genome, which contains five *x*=8 subgenomes^13^, exhibited strongly conserved synteny across the Caryophyllales species evaluated. Accordingly, *x*=8 is supported as the ancestral karyotype for the lineage^33^. *Ancistrocladus* and *Triphyophyllum* shared a chromosomal fission followed by a tetraploidy event, accounting for their higher basic chromosome number (*x*=9, **Fig. 5a**). *Dionaea* was resolved as an allotetraploid, with two distinct subgenomes (**Fig. 5a,b, Supplementary Materials Fig. S7)**, indicating a hybrid origin. Among sundews, most *D. regia* chromosomes matched *N. gracilis* chromosomes in a 3:1 pattern (**Fig. 5a**), consistent with *D. regia*’s proposed hexaploid structure. This could be further resolved into 2:1 phasing using Ks values (**Fig. 5c**; **Supplementary Materials Fig. S8**)

**Fig. 5:**
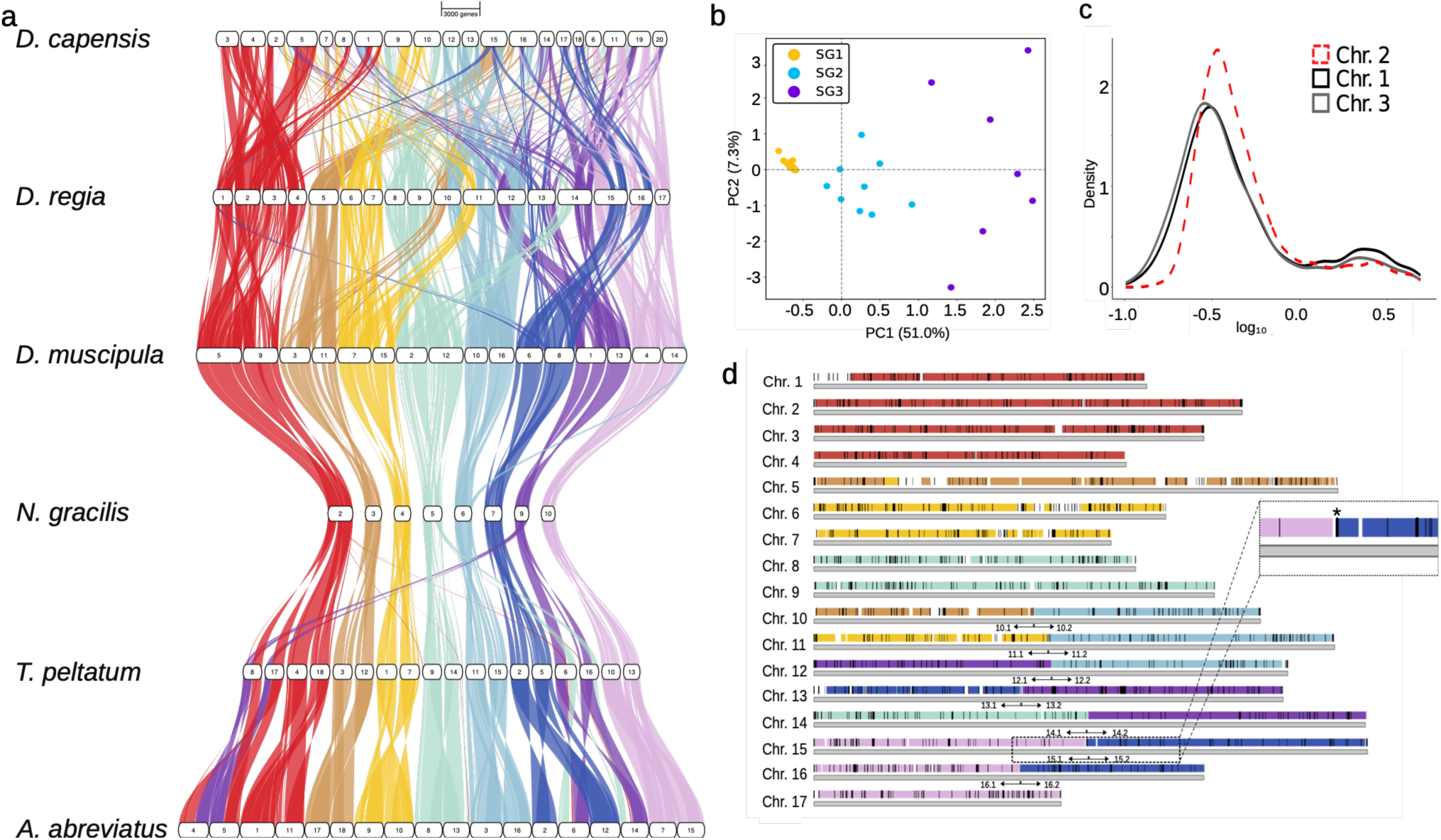
Structural rearrangements in *Drosera* genomes with contrasting inferred centromere organization. (**a**) Synteny of the dominant *N. gracilis* subgenome (8 chromosomes) with *Ancistrocladus abbreviatus*, *Triphyophyllum peltatum*, *Dionaea muscipula*, hexaploid *D. regia*, and dodecaploid *D. capensis*. *Dionaea* is doubled but minimally rearranged; *D. regia* shows triplicate structure, whereas *D. capensis* shows sixfold/dodecaploid organization with extensive rearrangement; the two genomes share broad ancestral substrate but do not resolve as a simple shared three-subgenome architecture. (**b**) SubPhaser PCA of *Dionaea* and D. regia assemblies indicates *Dionaea* is an allopolyploid of subgenomes 2 (cyan) and 3 (purple), with *D. regia* (yellow) diverging in k-mer composition (see **Supplementary Materials Fig. S7**). (**c**) Ks density plot phases *D. regia*’s triplicate subgenomes into a 2:1 pattern: homoeologous chromosomes 1 and 3 show similar Ks distributions, whereas homoeolog 2 shows higher Ks, indicating greater sequence divergence (x = log₁₀ Ks; y = density; see **Supplementary Materials Fig. S8**). **(d)** Black bars mark *Regi90* arrays relative to the inferred *D. regia* fusion/fission boundaries; chromosome 15 provides one example of an array near a boundary. Background colors retain the *N. gracilis*-derived synteny assignments from **(a)**. Proximity alone does not establish *Regi90* as the cause of a rearrangement.

Notably, several inferred fusion/fission breakpoints in *D. regia* occur at or near *Regi90* arrays (**Fig. 5d**). This spatial association is compatible with, but does not demonstrate, the possibility that repetitive DNA provided homologous substrates for ectopic recombination during structural reorganization^25,29^. In contrast, *D. capensis* showed substantial chromosomal rearrangement and reduced syntenic collinearity. Nonetheless, a 6:1 match with the eight-chromosome dominant subgenome of *N. gracilis* supports a dodecaploid organization in *D. capensis* that is consistent with additional multiplication of an older hexaploid-related substrate, but does not by itself show that *D. capensis* is simply a doubled retained version of the extant *D. regia*-like three-subgenome architecture (**Fig. 5a**).

Using a concatenated *D. regia* and *N. gracilis* hybrid assembly in CoGe SynMap^68^, Ks-colored syntenic anchors separated younger, intermediate, and older classes of duplicated or interspecific gene pairs (**Fig. 6a,b**). Within *D. regia*, these Ks classes are consistent with a 2:1 phasing among three homoeologous chromosomes, with two more closely related copies and one more divergent copy. Fractionation-bias profiles further showed that the three *D. regia* homoeologs retain comparable gene-retention landscapes relative to the dominant *N. gracilis* subgenome, whereas recessive *N. gracilis* chromosomes generally show lower retention (**Fig. 6c-e**). We use these comparisons mainly to phase the three *D. regia* homoeologs and to compare their retention profiles with *N. gracilis*; they are not intended to order every polyploidization event. Saul et al.^13^ inferred that the additional *N. gracilis* subgenomes came from closely related ancestors, which may help explain why *Nepenthes* is less discordant in the cloudogram than *Dionaea* or *D. regia* (**Fig. 4**).

**Fig. 6:**
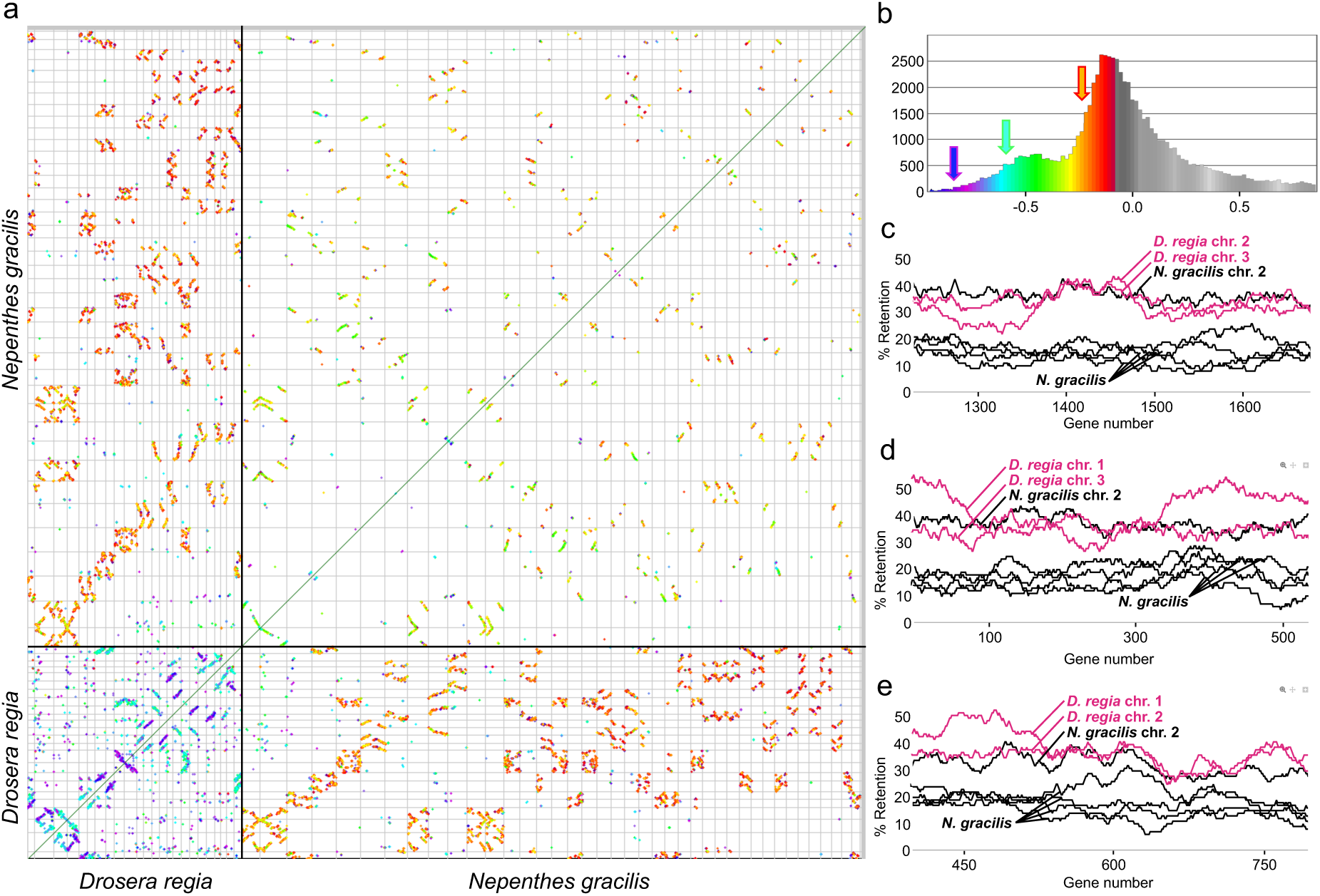
Ks-resolved synteny and fractionation relationships between *Drosera regia* and *Nepenthes gracilis*. **(a)** SynMap dot plot of the concatenated *D. regia* and *N. gracilis* assemblies compared against themselves. The upper-right and lower-left quadrants show within-genome paralogous relationships in *N. gracilis* and *D. regia*, respectively, whereas the off-diagonal quadrants show syntenic relationships between the two species. Syntenic anchors are colored by log₁₀(Ks), using the scale summarized in panel **b**. **(b)** Distribution of log₁₀(Ks) for the syntenic gene pairs in **(a)**. Purple/blue, green/yellow, and orange/red bins indicate the youngest, intermediate, and oldest highlighted anchor classes, respectively; grey bins contain still higher-divergence pairs. Bars drawn on opposite sides of the axis are separated only for display and do not represent negative counts. **(c-e)** Representative fractionation-bias profiles. Pink traces correspond to the three *D. regia* homoeologs; black and grey traces are the matched *N. gracilis* chromosomes. The three *D. regia* copies have broadly similar retention profiles against the dominant *N. gracilis* subgenome, while recessive *N. gracilis* chromosomes generally retain fewer genes. These panels are used to support the 2:1 Ks phasing of *D. regia* and to place it beside the multi-subgenome *N. gracilis* genome; they do not determine the full chronology of polyploid events.

To assess whether *D. capensis* and *D. regia* share a single higher-order hexaploid restructuring history, we combined additional synteny and topology analyses, Ks comparisons of species splits and polyploidy events (**Supplementary Materials Fig. S9**), and quantitative comparisons of genome structure (described in detail in **Supplementary Note 2**). Across these analyses, *D. regia* and *D. capensis* consistently showed broad syntenic continuity and related ancestral genomic substrate, and *D. capensis* retained strong internal duplicated structure. The combined evidence does not support a simple doubling of the extant *D. regia*-like triplet in *D. capensis*, and we did not recover a unique higher-order rearrangement history shared by the two genomes. One analysis nevertheless revealed a modest excess of shared connections between ancestral chromosome blocks, consistent with the persistence of some ancestral structural features (**Supplementary Note 2 Fig. 2d**). Overall, the results support shared ancestral chromosome substrate followed by separate higher-order restructuring. The strongest *D. regia*-*Dionaea* relationship is confined to one inferred *D. regia* subgenome; the remainder of the *D. regia* genome does not show an equally strong or stable relationship to *Dionaea*. Together with the conflicting gene-tree placements, this favors a reticulate interpretation but does not justify treating *D. regia* and *D. capensis* as descendants of a single conserved hexaploid architecture. Subgenome-wise phylogenetic reconstruction, for example using multilabeled gene tree methods (cf. Tomlin et al., 2024^69^), and additional comparative sampling across *Drosera* will be needed to resolve the timing and origin of these ancient multiplication events. They may also help determine whether the major fusions in *D. regia* preceded or followed holocentric chromosome evolution, and whether *D. capensis* retains an ancestral monocentric state or represents a secondary reversion.

### A reticulate allopolyploid origin model for D. regia

Figure 7 combines the backbone phylogeny, synteny, fractionation, Ks, and chromosome-number evidence into a working genome-history model for the sampled carnivorous Caryophyllales (**Supplementary Note 3**). The left-hand network starts from an ancestral *x*=8 substrate, most directly represented by the dominant *Nepenthes* subgenome, and adds the lineage-specific multiplications, fusions or fissions, and reticulate connections indicated by these analyses. In this view, *Triphyophyllum* and *Ancistrocladus* share one derived polyploid history; *Nepenthes* carries a more complex decaploid history; *Drosophyllum* is modeled as retaining an *x* = 8-derived framework modified by chromosome fusion; and *Aldrovanda* has a separate inferred hexaploidy. The two boxes at right concern *D. regia* alone. Both models assume a tripartite genome and a single component with stronger *Dionaea*-like structural affinity. Model I treats that component as a reticulate contribution from a *Dionaea*-like ancestor. Model II instead attributes the same signal to differential retention of an older shared component. Model I more directly links the structural affinity to the recurrent phylogenetic placement of *D. regia* near the snap-trap lineage (**Supplementary Materials Fig. S6**); model II remains viable because the fractionation signal itself does not establish direction or timing, and because *D. regia* remains morphologically sundew-like. In either case, *D. regia* and *D. capensis* share ancestral substrate without retaining the same higher-order three-subgenome assembly.

**Fig. 7:**
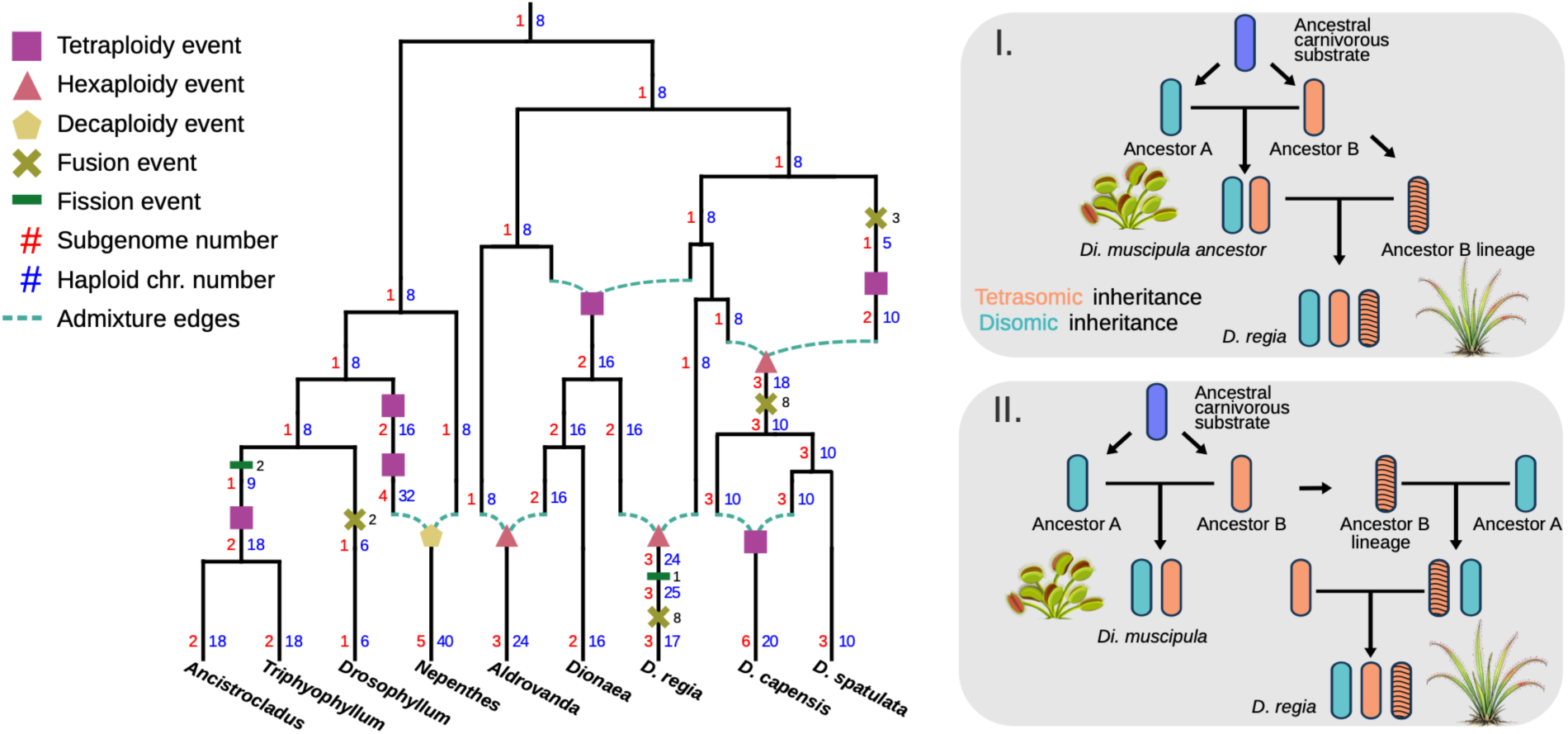
Reticulate genome-history framework for sampled carnivorous Caryophyllales and alternative origin scenarios for the tripartite genome of *Drosera regia*. The left network combines the sequence backbone with synteny, chromosome number, Ks, and subgenome/fractionation evidence. Red numbers give inferred subgenome number and blue numbers give haploid chromosome number; symbols mark the inferred polyploidy, fusion/fission, and admixture events. The reconstruction begins with an *x*=8 carnivorous substrate, represented most directly by the dominant *Nepenthes* subgenome. *Triphyophyllum* and *Ancistrocladus* then follow a shared derived polyploid trajectory, *Drosophyllum* is modeled as retaining an *x* = 8-derived framework modified chiefly by chromosome fusion, *Nepenthes* carries the more complex five-component/decaploid history, and *Aldrovanda* is assigned a separate hexaploidy. *Nepenthes* carries the more complex five-component/decaploid history, and *Aldrovanda* is assigned a separate hexaploidy. The two grey boxes show alternative explanations for the tripartite *D. regia* genome. In model I, the structurally *Dionaea*-like component – channel R2 in **Supplementary Note 2** – is a reticulate contribution from a *Dionaea*-like ancestor, with the other two components coming from the broader *Drosera*-side background. In model II, the same R2 affinity results from differential retention of an older component already present deeper in the ancestry, rather than from a direct *Dionaea*-like donor. Model I is preferred based on the combined structural and phylogenomic evidence; model II remains the conservative alternative and is compatible with the strongly *Drosera*-like morphology of *D. regia*. Both models require broad ancestral substrate shared by *D. regia* and *D. capensis*, but neither treats *D. capensis* as a simple doubled copy of the extant *D. regia* three-component architecture.

We also examined post-polyploid gene retention and loss as a separate genomic test of this interpretation (**Supplementary Note 3**). FractBias copy sets were tested for shared fractionation scars; an HMM phased local copy correspondences along ancestral *Vitis* coordinates; chromosome-local copies were threaded into species-wide components; and a transitive optimization grouped those threads into inferred parental-lineage classes before any sequence-tree reconciliation. Using synteny-derived copy depths, the analysis recovered cross-species component relationships consistent with **Fig. 7**: *Dionaea* has two inferred classes, *D. regia* three, *Nepenthes* five, and *D. capensis* six, with only one *D. regia* thread sharing a widely distributed class with *Dionaea*. *D. capensis* contains all three *D. regia*-associated classes but adds three different ones rather than a second copy of the *D. regia* triplet. Together, these findings reinforce the mosaic genome-history framework in **Fig. 7**, revealing shared ancestral genomic components retained in different combinations across lineages.

These results are also relevant to the taxonomic position of *D. regia*, which may warrant recognition as a separate, monotypic genus outside of *Drosera*. The name *Freatulina regia* (Stephens) Chrtek & Slavíková is already available for this purpose^70^. Morphologically, the species is distinct from most sundews, bearing woody rhizomes, operculate pollen released in tetrads, undivided floral styles, and a non-circinate flowering scape: traits that are either considered ancestral or absent from other members of the genus^71^. The species also exhibits pronounced leaf-blade wrapping around prey^72^. Genomically, *D. regia* is also distinctive: it lies outside the main sampled *Drosera* lineage, but one component of its genome shows clear structural affinity to the snap-trap lineage, consistent with a mosaic allopolyploid history not captured by either placement alone.

## Conclusions

The two chromosome-scale assemblies place monocentric *D. capensis* beside a putatively holocentric *D. regia* genome. Their dominant satellites are organized differently: *Cape71* is localized in *D. capensis*, whereas *Regi90* is dispersed along *D. regia* chromosomes. *Regi90* should nevertheless be treated as a candidate centromere-associated repeat, since we have not demonstrated direct binding to functional kinetochore proteins. Comparative profiling across *Drosera*, including these two species, highlights rapid satDNA turnover and lineage-specific TE enrichments, both consistent with recurrent centromere remodeling. Indeed, the extended regional centromeres recently described in the Venus flytrap^49^ suggest that Droseraceae as a whole span a continuum of centromere organization.

Failure to detect KNL1 in *D. regia* is not, by itself, evidence that the protein is absent from centromeres; antibody cross-reactivity, fixation, or other technical factors could also explain the result. RNA-seq recovers *KNL1* and *NDC80*^49^ transcripts in both species, and each genome retains one clearly intact *CENH3* paralog plus a second paralog that is structurally compromised or ambiguous. We therefore base the holocentric interpretation on the combined cytological and genomic evidence rather than on a single negative antibody result.

Synteny also shows that *D. regia* and *D. capensis* retained much of the same ancestral genomic substrate, despite very different subsequent chromosome evolutionary histories. While *D. regia* preserves relatively large-scale organization, *D. capensis* is much more extensively rearranged. Repeat evolution, TE activity, and diploidization may all contribute to that difference. Holocentricity offers an additional, testable possibility: chromosome-wide kinetochore activity could make some rearranged chromosomes easier to transmit and retain in *D. regia*. We therefore view holocentric buffering as a hypothesis for the persistence of rearranged chromosomes in that species, not as a demonstrated consequence of polyploidy. Broader sampling is needed to test whether chromosome-wide centromere organization arose repeatedly within *Drosera*, and whether it preceded or followed the major structural changes seen in these genomes.

Our phylogenomic and subgenome results point to a mosaic paleopolyploid history for *D. regia*, with one inferred component showing a stronger relationship to the Venus flytrap lineage. *D. regia* and *D. capensis* share substantial ancestral substrate, but *D. capensis* is not simply a doubled copy of the extant *D. regia*-like architecture. Reticulate genome formation could have triggered TE activity and chromosome reorganization; conversely, the putative holocentric state of *D. regia* may have helped some rearranged chromosomes persist after that disruption^59,60^. These remain linked hypotheses that require experimental validation, rather than a demonstrated causal sequence.

The implications of this study may extend beyond Droseraceae. Holocentric chromosomes could help some lineages, including the uniformly holocentric sedges (Cyperaceae), tolerate chromosome change associated with polyploidization, but that possibility needs direct comparative tests. Additional *Drosera* genomes spanning the monocentric to chromosome-wide centromere range will provide a direct test of whether centromere organization predicts which rearrangements persist after polyploidization. If so, centromere architecture may help set the structural limits within which polyploid genomes stabilize and diversify.

## Acknowledgments

We acknowledge the following sources for funding: United States National Science Foundation grants (2030871 to V.A.A. and PRFB-2109178 to J.R.B.), Human Frontier Science Program Young Investigators grant (RGY0082/2021 to K.F. and R.T.), JSPS KAKENHI (23K20050 to K.F.), Max Planck Society (core funding to A.M.), European Research Council (ERC StG HoloRECOMB, grant no. 101114879 to A.M.), the German Science Foundation (DFG MA 9363/6-1 to A.M.), the Research Council of Finland (decision 318288 to J.S). L.A.R. is financially supported by DFG (German Research Foundation) grant MA 9363/6-1. K.S. is supported by RIKEN TRIP initiative. The DFG also funded this work under Germany’s Excellence Strategy – EXC 493 2048/1-390686111 (to A.M.).

## Disclosure

ChatGPT (GPT-5.6 Sol) was used to assist with language editing and to review computational workflows for internal consistency. The tool also supported the development and documentation of selected analytical approaches described in Supplementary Note 2. For Supplementary Note 3, ChatGPT assisted in developing the FractLineage methodology, including suggestions concerning its mathematical formulation and computational implementation, and the design of its graphical outputs. The tool also aided preparation of the evidence-flow diagram in Supplementary Note 3 Fig. 1. ChatGPT was also used to examine the consistency of mathematical and computational procedures and to improve their descriptions. The authors reviewed the AI-assisted methodological developments and retain full responsibility for the mathematical formulation, implementation, results, and scientific interpretation of all analyses. The associated scripts and reproducibility materials are described in the Data and Code Availability statements.

## Author contributions

V.A.A. conceived the research. V.A.A. and A.M. led the study, with contributions from J.S. and K.F. M.M.-R. provided *Drosera* plant material, and E.I.-L. extracted DNAs. S.C.S. and D.D.-M. sequenced *Drosera* genomes, J.S. and A.M. assembled them and S.R. annotated them. K.F. led the sequencing, assembly and annotation of the *Dionaea*, *Triphyophyllum* and *Ancistrocladus* genomes with sequencing support from K.S. and S.M. S.J.F. and A.M. phased the *Drosera* and *Dionaea* genomes, with contributions from J.R.B. L.A.R. performed all cytological analyses and led research on repetitive DNAs, the latter with contributions from J.K. (phylogenetic analyses) and T.P.M. L.A.R., S.J.F., A.B. performed the transcriptomic analyses for centromeric gene expression (Supplementary Note 1. A.M., A.B., and V.A.A. performed syntenic comparisons, with contributions from D.S., Y.Z. and C.Z. S.J.F. performed *ksrates* analyses. H.M. performed gene family and phylogenetic analyses, with contributions from S.J.F. and V.A.A. V.A.A., J.K., A.M. and L.A.R. developed phylogenetic network and allopolyploidy reconstructions. Supplementary Note 2 was designed by A.B and V.A.A, and Supplementary Note 3 by V.A.A. Additional bioinformatic analyses were done by T.L., M.R., T.Q.W. and J.S. Additional materials and research support were provided by D.B., G.B., M.F., R.H., L.H.-E., I.K., S.T., Y.V.d.P. and G.V. V.A.A., L.A.R., A.B. and A.M. wrote the manuscript and/or supplement, and all authors approved the final version.

## Competing interests

The authors declare no competing interests.

## Data and code availability

The *Triphyophyllum peltatum*, *Ancistrocladus abbreviatus*, and *Dionaea muscipula* genomes are available at NCBI via accessions BAAHMP010000001-BAAHMP010000429, BAAHMO010000001-BAAHMO010000131 and BAAGIJ010000001-BAAGIJ010003706, respectively. The processed *Drosera* reference genomes and sequencing reads are available at the European Nucleotide Archive at PRJEB128437. Processed RNA-seq data files, the ancestral 24 pre-fusion chromosomes of *D. regia*, all BUSCO gene trees, and BUSCO single copy gene trees for DensiTree are available at Zenodo (pending: XXX, XXX, and XXX), as are all custom scripts and other materials required for replication of analyses and graphics. The REXdb database Viridiplantae v.3.0 is publicly available at Github (https://github.com/repeatexplorer/rexdb).

## Notes

### Competing Interest Statement

The authors have declared no competing interest.

### Summary of Updates

Revised version for resubmission 11 November 2026

## References

1 Albert, V. A., Williams, S. E. & Chase, M. W. Carnivorous plants: phylogeny and structural evolution. Science 257, 1491–1495 (1992).

2 Fleischmann, A. et al. Evolution of carnivory in angiosperms. Carnivorous plants: physiology, ecology, and evolution 1, 22–42 (2018).

3 Fukushima, K. et al. Oriented cell division shapes carnivorous pitcher leaves of Sarracenia purpurea. Nature Communications 6, 6450 (2015).

4 Whitewoods, C. D. et al. Evolution of carnivorous traps from planar leaves through simple shifts in gene expression. Science 367, 91–96 (2020).

5 Bemm, F. et al. Venus flytrap carnivorous lifestyle builds on herbivore defense strategies. Genome Research 26, 812–825 (2016).

6 Pavlovič, A. & Mithöfer, A. Jasmonate signalling in carnivorous plants: copycat of plant defence mechanisms. Journal of experimental botany 70, 3379–3389 (2019).

7 Adlassnig, W. et al. Endocytotic uptake of nutrients in carnivorous plants. The Plant Journal 71, 303–313 (2012).

8 Freund, M. et al. The digestive systems of carnivorous plants. Plant physiology 190, 44–59 (2022).

9 Płachno, B. et al. Fluorescence labelling of phosphatase activity in digestive glands of carnivorous plants. Plant Biology 8, 813–820 (2006).

10 Fukushima, K. et al. Genome of the pitcher plant Cephalotus reveals genetic changes associated with carnivory. Nature Ecology & Evolution 1, 0059 (2017).

11 Lan, T. et al. Long-read sequencing uncovers the adaptive topography of a carnivorous plant genome. Proceedings of the National Academy of Sciences 114, E4435–E4441 (2017).

12 Pavlovič, A. How the diversity in digestion in carnivorous plants may have evolved. New Phytologist (2025).

13 Saul, F. et al. Subgenome dominance shapes novel gene evolution in the decaploid pitcher plant Nepenthes gracilis. Nature plants 9, 2000–2015 (2023).

14 Cross, A. T., Krueger, T. A., Gonella, P. M., Robinson, A. S. & Fleischmann, A. S. Conservation of carnivorous plants in the age of extinction. Global Ecology and Conservation 24, e01272 (2020).

15 Fleischmann, A., Cross, A. T., Gibson, R., Gonella, P. M. & Dixon, K. W. Systematics and evolution of Droseraceae. Oxford Scholarship Online (2018).

16 Metcalfe, C. R. The Anatomical Structure of the Dioncophyllaceae in Relation to the Taxonomic Affinities of the Family. Kew Bulletin 6, 351–368 (1951).

17 Green, S., Green, T. L. & Heslop-Harrison, Y. Seasonal heterophylly and leaf gland features in Triphyophyllum (Dioncophyllaceae), a new carnivorous plant genus. Botanical Journal of the Linnean Society 78, 99–116 (1979).

18 Winkelmann, T., Bringmann, G., Herwig, A. & Hedrich, R. Carnivory on demand: phosphorus deficiency induces glandular leaves in the African liana Triphyophyllum peltatum. New Phytologist 239, 1140–1152 (2023).

19 Martín-Rodríguez, I., Vargas, P., Ojeda, F. & Fernández-Mazuecos, M. An enigmatic carnivorous plant: ancient divergence of Drosophyllaceae but recent differentiation of Drosophyllum lusitanicum across the Strait of Gibraltar. Systematics and Biodiversity 18, 525–537 (2020).

20 Płachno, B. J., Kapusta, M., Stolarczyk, P., Świątek, P. & Lichtscheidl, I. Differences in the Occurrence of Cell Wall Components between Distinct Cell Types in Glands of Drosophyllum lusitanicum. International Journal of Molecular Sciences 24, 15045 (2023).

21 Shirakawa, J., Nagano, K. & Hoshi, Y. A chromosome study of two centromere differentiating Drosera species, D. arcturi and D. regia. Caryologia 64, 453–463 (2011).

22 Kolodin, P. et al. Holocentric chromosomes may be an apomorphy of Droseraceae. Plant Syst Evol 304, 1289–1296 (2018).

23 Sheikh, S. A., Kondo, K. & Hoshi, Y. Study of diffused centromeric nature of *Drosera* chromosomes. Cytologia 60, 43–47 (1995).

24 Shirakawa, J., Hoshi, Y. & Kondo, K. Chromosome differentiation and genome organization in carnivorous plant family Droseraceae. Chromosome Botany 6, 111–119 (2011).

25 Hofstatter, P. G. et al. Repeat-based holocentromeres influence genome architecture and karyotype evolution. Cell 185, 3153–3168 (2022).

26 Kuo, Y. T. et al. Holocentromeres can consist of merely a few megabase-sized satellite arrays. Nature Communications 14 (2023). https://doi.org/ARTN350210.1038/s41467-023-38922-7

27 Mata-Sucre, Y. et al. Repeat-based holocentromeres of the woodrush Luzula sylvatica reveal insights into the evolutionary transition to holocentricity. Nat Commun 15, 9565 (2024).

28 Castellani, M. et al. Meiotic recombination dynamics in plants with repeat-based holocentromeres shed light on the primary drivers of crossover patterning. Nat Plants 10, 423–438 (2024).

29 Zhang, M. et al. A holocentric pangenome links karyotype evolution to meiotic recombination. bioRxiv, 2026.2001.2017.700048 (2026).

30 Włodzimierz, P. et al. Pangenome analysis reveals the evolutionary dynamics of repeat-based holocentromeres. bioRxiv, 2026.2001.2017.700053 (2026).

31 Zhang, M. et al. Sex without crossovers mimics clonal reproduction in the holocentric plant <em>Rhynchospora tenuis</em>. bioRxiv, 2026.2001.2017.700054 (2026).

32 Ibarra-Laclette, E. et al. Architecture and evolution of a minute plant genome. Nature 498, 94–98 (2013).

33 Mohn, R. A. et al. Dramatic difference in rate of chromosome number evolution among sundew (Drosera L., Droseraceae) lineages. Evolution 77, 2314–2325 (2023).

34 Walker, J. F. et al. Widespread paleopolyploidy, gene tree conflict, and recalcitrant relationships among the carnivorous Caryophyllales. American Journal of Botany 104, 858–867 (2017).

35 Van de Peer, Y., Mizrachi, E. & Marchal, K. The evolutionary significance of polyploidy. Nature Reviews Genetics 18, 411–424 (2017).

36 Van de Peer, Y., Ashman, T.-L., Soltis, P. S. & Soltis, D. E. Polyploidy: an evolutionary and ecological force in stressful times. The Plant Cell 33, 11–26 (2021).

37 Barker, M. S., Jiao, Y. & Glennon, K. L. Doubling down on polyploid discoveries: Global advances in genomics and ecological impacts of polyploidy. American journal of botany 111, e16395 (2024).

38 Ræbild, A. et al. Polyploidy–A tool in adapting trees to future climate changes? A review of polyploidy in trees. Forest Ecology and Management 560, 121767 (2024).

39 Gaeta, R. T. & Chris Pires, J. Homoeologous recombination in allopolyploids: the polyploid ratchet. New Phytologist 186, 18–28 (2010).

40 Li, Z. et al. Patterns and processes of diploidization in land plants. Annual review of plant biology 72, 387–410 (2021).

41 Han, J. et al. Centromere-size reduction and chromatin state dynamics following intergenomic hybridization in cotton. PLoS genetics 21, e1011689 (2025).

42 Su, H. et al. Centromere Satellite Repeats Have Undergone Rapid Changes in Polyploid Wheat Subgenomes. Plant Cell 31, 2035–2051 (2019).

43 Trebing, S. et al. Impaired trap closure in the counting-deficient Venus flytrap mutant DYSCALCULIA is caused by cell wall biomechanics. bioRxiv, 2025.2006. 2026.661685 (2025).

44. University of Würzburg. Triphyophyllum peltatum genome assembly Tripe_1.2, <https://www.ncbi.nlm.nih.gov/datasets/genome/GCA_059738685.1/> (2025).

45. University of Würzburg. Dionaea muscipula genome assembly Diomu_v2.1, <https://www.ncbi.nlm.nih.gov/datasets/genome/GCA_054771495.1/> (2025).

46. University of Würzburg. Ancistrocladus abbreviatus genome assembly Ancab_1.1, <https://www.ncbi.nlm.nih.gov/datasets/genome/GCA_054771515.1/> (2025).

47 Veleba, A. et al. Evolution of genome size and genomic GC content in carnivorous holokinetics (Droseraceae). Annals of Botany 119, 409–416 (2017).

48 Marques, A. & Drinnenberg, I. A. Same but different: Centromere regulations in holocentric insects and plants. Current Opinion in Cell Biology 93, 102484 (2025).

49 Oliveira, L. et al. KNL1 and NDC80 represent new universal markers for the detection of functional centromeres in plants. Chromosome Research 32, 3 (2024).

50 Thakur, J., Packiaraj, J. & Henikoff, S. Sequence, chromatin and evolution of satellite DNA. International Journal of Molecular Sciences 22, 4309 (2021).

51 Comai, L., Maheshwari, S. & Marimuthu, M. P. Plant centromeres. Current Opinion in Plant Biology 36, 158–167 (2017).

52 Henderson, I. et al. Cyclical evolution of centromere architecture across 193 eukaryote species. Research Square preprint (2025).

53 Costa, L. et al. Repeat competition and ecological shifts drive the evolution of the mobilome in Rhynchospora Vahl (Cyperaceae), the holocentric beaksedges. Annals of Botany 135, 909–924 (2024).

54 Marques, A. et al. Holocentromeres in Rhynchospora are associated with genome-wide centromere-specific repeat arrays interspersed among euchromatin. Proceedings of the National Academy of Sciences 112, 13633–13638 (2015).

55 Wicker, T. et al. A unified classification system for eukaryotic transposable elements. Nature reviews genetics 8, 973–982 (2007).

56 Neumann, P. et al. Plant centromeric retrotransposons: a structural and cytogenetic perspective. Mobile DNA 2, 4 (2011).

57 Bousios, A., Kakutani, T. & Henderson, I. R. Centrophilic retrotransposons of plant genomes. Annual Review of Plant Biology 76 (2025).

58 Wells, J. N. & Feschotte, C. A field guide to eukaryotic transposable elements. Annual review of genetics 54, 539–561 (2020).

59 de Tomás, C. & Vicient, C. M. The genomic shock hypothesis: genetic and epigenetic alterations of transposable elements after interspecific hybridization in plants. Epigenomes 8, 2 (2023).

60 Shimizu, K. K. Robustness and the generalist niche of polyploid species: genome shock or gradual evolution? Current Opinion in Plant Biology 69, 102292 (2022).

61 Zedek, F. & Bureš, P. Holocentric chromosomes: from tolerance to fragmentation to colonization of the land. Annals of Botany 121, 9–16 (2018).

62 Manni, M., Berkeley, M. R., Seppey, M. & Zdobnov, E. M. BUSCO: assessing genomic data quality and beyond. Current Protocols 1, e323 (2021).

63 Wong, T. K. et al. IQ-TREE 3: Phylogenomic Inference Software using Complex Evolutionary Models. Molecular Biology and Evolution 43 (2026).

64 Mirarab, S. et al. ASTRAL: genome-scale coalescent-based species tree estimation. Bioinformatics 30, i541–i548 (2014).

65 Stull, G. W., Pham, K. K., Soltis, P. S. & Soltis, D. E. Deep reticulation: the long legacy of hybridization in vascular plant evolution. The Plant Journal 114, 743–766 (2023).

66 Bouckaert, R. R. DensiTree: making sense of sets of phylogenetic trees. Bioinformatics 26, 1372–1373 (2010).

67 Lovell, J. T. et al. GENESPACE tracks regions of interest and gene copy number variation across multiple genomes. elife 11, e78526 (2022).

68 Albert, V. A. & Krabbenhoft, T. J. in Polyploidy: Methods and Protocols (ed Yves Van de Peer) 19-45 (Springer US, 2023).

69 Tomlin, C. M. et al. Allopolyploid origin and diversification of the Hawaiian endemic mints. Nature Communications 15, 3109 (2024).

70 Chrtek, J. & Slavíková, Z. Comments on the families Drosophyllaceae and Droseraceae. *Journal of the National Museum (Prague)*, Natural History Series 165, 139-141 (1996).

71 Gibson, R. Drosera arcturi in Tasmania and a comparison with Drosera regia. Carnivorous Plant Newsletter 28, 76–80 (1999).

72 Poppinga, S., Masselter, T., Hartmeyer, I., Hartmeyer, S. & Speck, T. Trap diversity and evolution in the family Droseraceae. Plant Signaling & Behavior 8, e24685 (2013).

