## Supplementary Materials for "Centromere divergence and allopolyploidy reshape carnivorous sundew genomes"

**This file includes:**

Materials and Methods

Figs. S1 to S10

Tables S1 to S2

Supplementary Notes 1, 2 and 3

### Materials and methods

***Plant materials and genome sequencing.*** In vitro cultures of *Drosera regia* and *Drosera capensis* were initiated from seeds derived from plants maintained in the carnivorous plant collection at the Red Manejo Biotecnológico de Recursos, Instituto de Ecología, A.C., Xalapa, Veracruz, Mexico. For superficial disinfection, seeds were placed in filter paper envelopes (Whatman No. 1, 110 mm diameter) and submerged in sterile distilled water for 30 minutes. They were then soaked in a 10% (v/v) commercial bleach solution (1.8% NaOCl) containing two drops of Tween-80 per 100 mL (Sigma, St. Louis, MO) for 10 minutes. This was followed by four rinses with sterile distilled water under aseptic conditions. The disinfected seeds were sown in 125-mL baby food jars containing 25 mL of half-strength MS medium[1], supplemented with 30 $g \cdot L^{-1}$ sucrose. The pH was adjusted to 5.0 ± 0.1 using 0.5 N NaOH and 0.5 N HCl prior to the addition of 7.5 $g \cdot L^{-1}$ Agar, Plant TC (Caisson A111), followed by autoclaving at 1.2 $kg \cdot cm^{-2}$ and 120 °C for 15 minutes. Cultures were incubated in a growth chamber at 25 ± 1 °C under a 16-hour photoperiod provided by LED lamps (50 $\mu mol \cdot m^{-2} \cdot s^{-1}$). Plants obtained from germinated seeds were subcultured every 3-4 months on the same medium to promote growth and multiplication.

High-molecular-weight DNA was extracted from nuclei isolated from the young leaves of *Drosera* species, following the protocol by Steinmüller and Apel[2]. To reduce contamination from chloroplast and mitochondrial DNA, nuclei were first collected from a 60% Percoll (Invitrogen) density gradient after centrifugation at 4000g for 10 minutes at 4°C. Next, high-quality megabase-sized DNA was obtained using the MagAttract HMW DNA Kit (Qiagen). Before library preparation for sequencing, the integrity of the high molecular weight (HMW) DNA was confirmed using pulsed-field gel electrophoresis (CHEF-DRIII system, Bio-Rad), as described elsewhere[3]. For library preparation, 10 µg of DNA were sheared to a fragment size range of 10-40 kb using a Covaris g-TUBE. The resulting fragment distribution was verified by pulsed-field gel electrophoresis. The sheared DNA was purified using 0.45× AMPure PB beads (Pacific Biosciences) following the manufacturer's protocol.

Library preparation for the PacBio RS II instrument was carried out according to the PacBio 20 kb SMRTbell Template Preparation Protocol, using 5 µg of the sheared DNA as input material. After preparation, the library size distribution was analyzed on an Agilent DNA 12000 Bioanalyzer chip to determine the appropriate size-selection cut-off. Libraries were size-selected with a Sage Science BluePippin system, employing a dye-free 0.75% agarose cassette and a 15 kb cut-off. The selected libraries were reanalyzed on the Bioanalyzer to confirm size distribution. Two libraries per species were prepared for *D. capensis* and *D. regia*. The *D. capensis* library was sequenced on two SMRT cells of the PacBio RSII single-molecule sequencing platform at loading concentrations of 0.15 nM and 0.2 nM, respectively. The *D. regia* library was sequenced on eight SMRT cells at a loading concentration of 0.2 nM. For Illumina sequencing of *Nepenthes gracilis*, *D. capensis*, *D. regia* and the ten additional *Drosera* species, the Illumina HiSeq 2500 (rapid run, 2x250bp; https://www.illumina.com/documents/products/datasheets/datasheet_hiseq2500.pdf) was employed.

***CHiCAGO and Hi-C library preparation and sequencing.*** ~500 ng of high-molecular-weight genomic DNA (mean fragment length = 150 kb) was reconstituted into chromatin *in vitro* for the ChiCAGO library[4]. Chromatin was fixed with formaldehyde for the CHiCAGO library. For the HiRise Hi-C library[5], chromatin was fixed in place with formaldehyde in the nucleus. Fixed chromatin was digested with DpnII, 5′ overhangs filled in with biotinylated nucleotides, and free blunt ends were ligated. After ligation, cross-links were reversed and the DNA was purified from protein. Purified DNA was treated to remove biotin that was not internal to the ligated fragments. The DNA was then sheared to a mean fragment size of ~350 bp, and sequencing libraries were generated using NEBNext Ultra enzymes and Illumina-compatible adaptors. Biotin-containing fragments were isolated using streptavidin beads before PCR enrichment of each library.

***Genome assembly and Hi-C scaffolding.*** The genomes were assembled using a hybrid strategy because the PacBio data were relatively low coverage and generated on the RS II platform, consisting of continuous long reads with relatively high raw error rates. The total unfiltered PacBio yield was 12 Gb for *D. capensis* and 6.6 Gb for *D. regia*, corresponding to approximately 42× and 23× genome coverage, respectively. We tested long-read-only assembly with Canu[6] for *D. regia*, but this produced a less contiguous assembly than the MaSuRCA[7] hybrid assembly[8]. Specifically, the Canu assembly was approximately 250 Mb with a contig N50 of 193 kb, whereas the MaSuRCA hybrid assembly was 282 Mb with a contig N50 of 287 kb. We therefore used the MaSuRCA hybrid pipeline for both species to improve assembly contiguity and maintain methodological comparability between the two genomes. Independent short-read genome-size estimates were generated from Illumina reads using KmerGenie[9], which identified predicted best k-mer sizes of 127 for *D. capensis* and 141 for *D. regia*, corresponding to estimated genome sizes of 286,140,113 bp and 348,732,445 bp, respectively.

For the hybrid assembly, the 2x250bp Illumina reads were first filtered for adapter sequences using Trimmomatic v0.36[10]. The hybrid assembly including the filtered reads and PacBio RS II reads was then carried out using MaSuRCA v3.2.1[8]. Assembly completeness was evaluated with BUSCO v4.0.5[11] using the embryophyta_odb10 lineage dataset, with an additional assessment using the eudicots_odb10 lineage dataset reported in **Supplementary Materials Table S1**. Pre-scaffolding contig counts and contig N50 values were calculated from the draft assemblies. QUAST v4.3[12] was subsequently run on the final scaffold-level assemblies; accordingly, the QUAST sequence counts, largest-sequence values, and N50, N75, L50, and L75 statistics reported in **Supplementary Materials Table S1** describe scaffolds rather than pre-scaffolding contigs.

The input *de novo* assembly, CHiCAGO library reads, shotgun reads, and Dovetail Hi-C library reads were then used as input data for HiRise[5], a software pipeline designed specifically for using proximity ligation data to scaffold genome assemblies. An iterative analysis was conducted. First, shotgun and CHiCAGO library sequences were aligned to the draft input assembly using a modified SNAP read mapper (http://snap.cs.berkeley.edu). The separation between CHiCAGO read pairs mapping within draft scaffolds was analyzed by HiRise to produce a likelihood model for the genomic distance between read pairs, and the model was used to identify and break putative misjoins, to score prospective joins, and to make joins above a selected threshold. After aligning and scaffolding CHiCAGO data, Dovetail Hi-C library sequences were aligned and scaffolded following the same method. After scaffolding, shotgun and PacBio sequences were used to close gaps between contigs.

Assembly-quality statistics were reported directly rather than converted into formal Earth BioGenome Project quality classes. The final *D. capensis* assembly spans 284,038,097 bp, with a contig N50 of approximately 1.0 Mb, a scaffold N50 of 12.65 Mb, and 94.8%

complete embryophyta_odb10 BUSCOs. The final *D. regia* assembly spans 282,006,236 bp, with a contig N50 of 287 kb, a scaffold N50 of 15.76 Mb, and 96.7% complete embryophyta_odb10 BUSCOs. Relative to Earth BioGenome Project reference benchmarks, the *D. capensis* assembly approximates the megabase-scale contig-continuity target, while both assemblies show chromosome-scale scaffold continuity and high recovery of conserved gene space. However, formal 6.C.Q40 or 5.C.Q40 designations were not assigned because all required metrics, including consensus quality value, k-mer completeness, and the percentage of assembled sequence assigned to chromosome-scale scaffolds, were not evaluated within a unified Earth BioGenome Project assessment. The lower contig-level continuity of the *D. regia* assembly likely resulted in part from the considerably lower long-read coverage available for this species.

***Transcriptome sequencing and genome annotation.*** Total RNA was extracted from *D. capensis* leaf tissues and petioles using the Spectrum™ Plant Total RNA Kit (Sigma-Aldrich. For RNA-seq, sample preparation employed a single stranded mRNA library kit, and libraries were subsequently sequenced using an Illumina HiSeq instrument. To generate our *D. capensis* transcriptome assembly used to annotate both the *D. capensis* and *D. regia* genomes, we first merged the *D. capensis* RNA-seq data from the two tissues sequenced. This amounted to a total of 105,784,845 read pairs. We then assembled one de novo transcriptome using transAbyss v2.0.1[13]. In transAbyss, we used multiple k-mers (33-75) in steps of 2 to generate multiple assemblies before merging them all into a single large set. We then assembled a second de novo transcriptome using Trinity v2.6.6[14] using the default k-mer size of 25. We generated one reference-guided transcriptome by first mapping the RNA-seq reads against the reference *D. capensis* genome using HISAT2 v.2.1.0[15] and then assembling the transcriptome using StringTie v.1.3.4c[16]. We then passed the three transcriptome assemblies to EvidentialGene v2017.12.21[17] which produced a final high confidence transcriptome assembly.

For repeat masking, we first generated a de novo repeat library for the *D. capensis* genome using RepeatModeler v1.0.9[18] and then masked the genome using RepeatMasker v4.0.7[19]. For gene model prediction, the transcriptome assembly was first splice-aligned against the unmasked reference *D. capensis* genome using PASA v2.2.0[20] to generate ORFs. Secondly, ab initio gene model prediction was carried out using BRAKER v.2.0.3[21], which internally used the reference aligned RNA-seq data and Genemark-ET to train AUGUSTUS v3.3[21,22] for its final prediction. Additionally, the self-training Genemark-ES v.4.33[23] was run independently to generate a second set of predictions. Finally the 2 prediction tracks and two spliced-alignment tracks were passed to the combiner tool Evidence Modeler v1.1.1[20] with highest weights to transcriptome evidence and lowest weights to Genemark-ES to generate a final high confidence gene prediction set. This final prediction set was re-run through PASA to update the gene models, add UTRs and identify and generate alternate spliced models.

For *D. regia*, the *D. capensis* transcriptome assembly was used as evidence for training purposes. First, the assembly was splice-aligned against the *D. regia* genome using PASA v.2.2.0[20] to generate ORFs. Additionally, *Arabidopsis thaliana* gene models were aligned against the genome using exonerate v2.2.0[24]. Genemark-ES v4.33[23] was then used to generate one set of predictions. BRAKER v2.0.3[21] was used in the protein mode where the *D. capensis* gene models were aligned against the *D. regia* genome using GenomeThreader v1.7.1 (https://genomethreader.org/), and that set was used to train BRAKER v2.0.3 to generate a second set of gene models. AUGUSTUS v3.3[21,22] was run independently using *D.*

*capensis* parameters to generate a third set of predictions for *D. regia*. All of these gene predictions were then passed to Evidence Modeler v1.1.1[20] to generate a single high confidence gene prediction set. This set was once again passed through PASA to update the gene models with UTR regions and also to generate the splice variants.

***Synteny analysis.*** Synteny analyses were performed with GENESPACE (https://github.com/jtlovell/GENESPACE)[25]. To identify possibly shared chromosomal rearrangements, we first identified all end-to-end fusions of non-homoeologous chromosomes within the *D. regia* genome. For each such fusion event, we then extracted corresponding genomic regions from *D. capensis* that involved the same ancestral *Nepenthes*-like chromosomes. Candidate shared rearrangements were identified where breakpoint boundaries in *D. regia* corresponded to syntenic block boundaries in *D. capensis* involving the same ancestral *Nepenthes*-like chromosome compartments. These candidates were subsequently evaluated through the formal structural comparisons described in **Supplementary Note 2**, below, rather than being treated as independent proof of a shared rearrangement event.

Further synteny analyses were performed using the CoGe SynMap platform (https://genomevolution.org/coge/SynMap.pl)[26]. Synteny plots were obtained using the following steps: (1) using the Last tool, (2) synteny analysis was performed using DAGChainer, using 20 genes as the maximum distance between two matches (-D) and 5 genes as the minimum number of aligned pairs (-A). Then (3) either default (no syntenic depth) or Quota Align (with syntenic depth) was used with overlap distance of 40 genes, and (4) orthologous and paralogous blocks were differentiated according to the synonymous substitution rate (Ks) using CoGe-integrated CodeML, and represented with different colors in the dot plot. FractBias[27] was run using Quota Align window size of 100 for all genes in the target genome, with a syntenic depth ratio of 12:12 with maximum query and target chromosome numbers of 64 each for the concatenated *D. regia-Nepenthes* genome assembly.

For the characterization of regions involved in fusions, we followed Hofstatter et al, 2022[28]. The syntenic alignment obtained in GENESPACE between the *D. regia* genome and the *N. gracilis* dominant subgenome allowed us to pinpoint regions around the borders of proposed homologous fusion events. To evaluate homology, we loaded and compared annotation features for genes, TEs, and tandem repeats along the syntenic alignments using Geneious (https://www.geneious.com).

For some structural analyses, given the unusually clear breakpoints between ancestral chromosomes apparent in *D. regia*'s GENESPACE alignment (see, e.g., **Fig. 5 a,d**), we manually segregated ancestral chromosomal blocks to simulate the primordial, pre-fusion *D. regia* karyotype architecture. For example, our origin scenario for the modern *D. regia* genome (as shown in **Fig. 7**) depended on resolving eight post-allohexaploid chromosomal fusion events. SubPhaser results (see **Fig. 5b** and ***Subgenome-aware phasing of the* D. regia *and* Dionaea muscipula *genomes***, immediately below) also depended on the manually generated ancestral, pre-fusion *D. regia* genome.

***Subgenome-aware phasing of the* D. regia *and* Dionaea muscipula *genomes.*** We used SubPhaser[29] (default parameters) to phase and partition the subgenomes of *D. regia* and *Dionaea muscipula* genomes simultaneously by assigning chromosomes to subgenomes based on differential repetitive k-mers. Additionally, Ks distributions of homoeologous duplicate gene pairs were extracted from CoGe SynMap calculations (above) to generate density plots for each triplet of *D. regia* ancestral chromosomes. Density plots were generated in R using the tidyverse[30], ggplot2[31], RColorBrewer[32], ggridges[33], and ggpmisc[34] packages.

**ksrates *analysis.*** *ksrates* version 1.1.4[35] was used to position species splits relative to polyploidy events. Coding sequence (CDS) fasta files were extracted using AGAT version 1.4.0[36]. Paralogous Ks (synonymous substitutions per synonymous site) peaks were generated for the following focal species: *Ancistrocladus abbreviatus* (this study), *Dionaea muscipula* (this study), *D. capensis* (this study), *D. regia* (this study), *Nepenthes gracilis*[37], and *Triphyophyllum peltatum* (this study). *Beta vulgaris* (GCA_026745355.1), *Coffea canephora*[38], *Gelsemium elegans* (CoGe id64491), and *Spinacia oleracea* (GCA_020520425.1) were also included in the analysis, but not as focal species. Orthologous Ks peaks were generated for each required species pair.

For the tree topology used in *ksrates*, OrthoFinder v2.5.5[39] was run with default settings to generate a tree for all species listed above. Each annotation was reduced to the longest isoform and proteins were extracted using AGAT version 1.4.0[36]. The species tree inference was performed with STAG[40], which uses the proportion of species trees derived from single-locus gene trees supporting each bipartition as its measure of support, and rooted using STRIDE[41].

***Repeat characterization.*** DANTE and DANTE-LTR retrotransposon identification (Galaxy Version 3.5.1.1) pipelines[42] were used to identify full-length LTR retrotransposons in the assembled genomes of *D. capensis* and *D. regia*, using a set of protein domains from REXdb[43]. All complete LTR-RTs contain GAG, PROT, RT, RH and INT domains, including some lineages encoding additional domains, such as chromodomains (CHD and CHDCR) from chromoviruses or ancestral RNase H (aRH) from Tat elements. DANTE_LTR retrotransposon filtering (Galaxy Version 3.5.1.1) was used to search for high quality retrotransposons, those with no cross-similarity between distinct lineages. This tool produced a GFF3 output file with detailed annotations of the LTR-RTs identified in the genome and a summary table with the numbers of the identified elements. Overall repeat composition was calculated excluding clusters of organelle DNA (chloroplast and mitochondrial DNA).

Tandem repeat sequences were identified using RepeatExplorer2 (https://repeatexplorer-elixir.cerit-sc.cz/) and further verified using the TideCluster pipeline (https://github.com/kavonrtep/TideCluster)[44]. All putative tandem sequences were compared for homology using DOTTER. These tandem sequences were individually mapped to the genome by BLAST[45], with 95% similarity in Geneious (https://www.geneious.com). The mapped sequence files were converted to BED and used as an input track for a genome-wide overview with ShinyCircos[46] using a 100kb window.

Comparative repeatome analysis of *Drosera* genomes was performed using Illumina reads from the species listed in **Supplementary Materials Table S2**. Analyses were carried out using RepeatExplorer2 on the public RepeatExplorer Galaxy server (https://repeatexplorer-elixir.cerit-sc.cz/). Reads were first quality-filtered within the RepeatExplorer2 workflow, retaining reads for which at least 95% of bases had a quality score of 10 or higher. Because genome sizes were unavailable for some of the analyzed species, each species-specific read dataset was downsampled to 1,000,000 reads to provide equal representation among taxa in the comparative analysis. Reads from all species were then analyzed jointly using the comparative clustering workflow in RepeatExplorer2. Clustering was performed using the standard RepeatExplorer2 settings of 90% sequence similarity over at least 55% of the read length, with other parameters left at their default values. Each resulting cluster therefore represents a group of sequence-similar reads recovered across the combined multispecies dataset, while retaining information on the contribution of each species to that cluster. For each species, repeat abundance was estimated from the proportion of that species' reads assigned to each cluster. Automated repeat annotations generated by RepeatExplorer2 were manually inspected and revised when necessary, followed by recalculation of the genomic proportion assigned to the corresponding repeat classes. Sequence-divergent clusters assigned to the same repeat lineage were retained as separate clusters rather than merged, allowing divergent subfamilies, lineage-specific variants, and independent repeat expansions to remain distinguishable in the comparative analysis. Computational resources for RepeatExplorer analysis were provided by the ELIXIR-CZ project (LM2023055), part of the international ELIXIR infrastructure.

To provide phylogenetic context for the comparative repeatome analysis, an independent assembly- and alignment-free phylogenetic analysis was performed using AAF (Assembly and Alignment Free[47]) on the Illumina short-read datasets. Genomic distances among species were estimated from shared k-mer profiles using a k-mer length of 21 bp and a minimum k-mer occurrence threshold of one. Singleton k-mer counts were incorporated into distance estimation. Phylogenetic support was assessed using 100 nonparametric bootstrap replicates generated by resampling the shared k-mer table and recalculating genomic distances for each replicate. The resulting bootstrap trees were summarized to obtain branch-support values for the AAF phylogeny shown alongside the comparative RepeatExplorer analysis in **Supplementary Fig. S5c**.

***Immunostaining and FISH.*** Flower buds of *D. capensis* and *D. regia* at various developmental stages were harvested and immediately fixed in freshly prepared 4% formaldehyde in Tris buffer (10 mM Tris-HCl (pH 7.5), 10 mM EDTA, and 0.5% Triton X-100), when immunostaining was intended, or ethanol-acetic acid (3:1 v/v) when only FISH was performed.

For immunostaining, fixation was carried out under vacuum infiltration for at least 15 minutes at room temperature, followed by an additional 45 minutes of incubation without vacuum. Fixed tissues were washed twice in 1× phosphate-buffered saline (1x PBS) at 4 °C until further processing. For enzymatic digestion, individual fixed buds were incubated in a solution containing 2% (w/v) cellulase (Onozuka R-10) and 2% (w/v) pectinase (Sigma) prepared in 1× PBS. Digestion was performed at 37 °C for 1 hour to facilitate cell wall degradation and release of nuclei. Following digestion, the softened tissue was gently macerated on a clean slide and squashed under the coverslip. Chromosome spreads were washed in 1× PBS for 5 minutes, followed by incubation in PBS-T1 buffer (1× PBS, 0.5% Triton X-100, pH 7.4) for 25 minutes. After two additional 5-minute washes in PBS, slides were incubated in PBS-T2 buffer (1× PBS, 0.1% Tween 20, pH 7.4) for 30 minutes. Primary antibody incubation was performed overnight at 4 °C using rabbit anti-CENH3 (specific for each species; **Supplementary Materials Fig. 2**), rabbit anti-KNL1[48] (GenScript, NJ, USA) and mouse anti α-tubulin (Sigma-Aldrich, St. Louis, MO; catalog number T6199) diluted 1:1,000 in blocking buffer (3% BSA, 1x PBS, 0.1% Tween 20, pH 7.4). Slides were then washed twice in 1x PBS (5 minutes each) and once in PBS-T2 (5 minutes), followed by incubation with secondary antibodies for at least 1 hour at room temperature. As the secondary antibody, goat anti-rabbit IgG antibody conjugated with Alexa Fluor 488 (Invitrogen; catalog number A27034), goat anti-rabbit conjugated with Rhodamine Red X (Jackson ImmunoResearch, catalog number: 111-295-144) or goat anti-mouse conjugated with Alexa Fluor 488 (Jackson ImmunoResearch; catalog number 115-545-166) were used in a 1:500 dilution. Final washes included two rounds in PBS and one in PBS-T2, each for 5 minutes. Slides were then mounted for fluorescence microscopy. Microscopic images were recorded using a Zeiss Axiovert 200M microscope equipped with a Zeiss AxioCam CCD. Images of at least 5 cells were analyzed using the ZEN software (Carl Zeiss GmbH).

For combined immuno-FISH, α-tubulin immunostaining and *Regi90* FISH were performed sequentially on the same *D. regia* metaphase-chromosome preparations. Immunostaining was first carried out using the mouse anti-α-tubulin primary antibody and Alexa Fluor-conjugated secondary antibody described above. The same slides were then processed for FISH using the Cy3-labelled *Regi90* oligonucleotide probe and the hybridization and washing conditions described below. Fluorescence channels corresponding to DAPI, α-tubulin, and *Regi90* were recorded from the same chromosome preparations and combined after channel registration. Because the spatial resolution of these preparations is insufficient to resolve direct molecular colocalization, the combined images were used only to assess broad spatial proximity between α-tubulin and chromosome-wide *Regi90* signals.

For FISH experiments, material fixation was performed for 2 days at 4 °C, washed in ice-cold water, and digested in a solution of 4% cellulase (Onozuka R10, Serva Electrophoresis, Heidelberg, Germany), 2% pectinase, and 0.4% pectolyase Y23 (both MP Biomedicals, Santa Ana, CA) in 0.01 M citrate buffer (pH 4.5) for 60 min at 37 °C. The digested material was transferred to a drop of 45% acetic acid, macerated and squashed under a coverslip. FISH was performed using species-specific oligonucleotide probes that were 5′-labeled with Cy3 during their synthesis:

*D. regia*__Regi90 (Cy3)-
CAAGTATTTCAATGGAAATGGTGAAATAACATGTTTTACACCTATTTCC;

*D. capensis*__Cape71 (Cy3)-
CCCTTTAAATGAGCTTAAAACACTCAAAACCCCTTGAAAAGGCTAAAAAC

FISH was performed as described in Macas, et al. 2007[49], with hybridization and washing temperatures adjusted to account for AT/GC content and hybridization stringency allowing for 10-20% mismatches. The slides were counterstained with 2 µg/mL DAPI in Vectashield (Vector) mounting medium. The images of at least 10 cells were captured as described above.

***Satellite DNA phylogeny.*** Satellite-repeat tracks were generated using the consensus sequences of the dominant satellite families identified with TideCluster, and the coordinates of *Cape71* and *Regi90* satDNA polymers were used to guide sequence extraction in Geneious Prime. (https://www.geneious.com). In total, 5,371 *Cape71* and 18,178 *Regi90* were extracted from *D. capensis* and *D. regia*, respectively. The collected satellite-repeat monomers were aligned using MAFFT v7.490[50]. Phylogenetic trees were inferred using FastTree v2.1.11[51] under the generalized time-reversible (GTR) model. Resulting trees were midpoint rooted and visualized and annotated using the Interactive Tree of Life (iTOL) web server[52], with colors corresponding to their chromosome of origin.

The phylogenetic trees were imported into R using the ape package[53]. Tip labels included chromosome ID and genomic coordinates. Genomic positions of sampled tips were plotted using ggplot2[31] as normalized positions (relative to chromosome length) in order to show the spatial distribution of tips along individual chromosomes. Tips were plotted using ggplot2, with color representing the order of appearance in the tree (NodeOrder), allowing assessment of how phylogenetic relationships correlate with genomic location. All analyses and plotting were performed in R (version 4.1.2) using the packages ape[53], dplyr[54], ggplot2[31], and viridis (https://sjmgarnier.github.io/viridis/).

***Species tree inference with BUSCO genes.*** To extract BUSCO genes, we collected coding sequence sets from sequenced genomes as well as previously assembled transcriptomes[55] (**Supplementary Materials Table S1**). We identified eudicot-conserved single-copy genes with BUSCO v5.3.2 (https://gitlab.com/ezlab/busco). Genes classified as single-copy (S) or fragmented (F) were retained, whereas those classified as duplicated (D) or missing (M) were considered absent, as described previously. Protein sequences were aligned with MAFFT v7.508 (https://mafft.cbrc.jp/alignment/server/index.html), trimmed with ClipKIT v2.1.1 (https://github.com/JLSteenwyk/ClipKIT), and back-translated to codons with CDSKIT v0.10.10 (https://github.com/kfuku52/cdskit) to produce in-frame nucleotide alignments. For each gene, nucleotide and protein maximum-likelihood trees were inferred with IQ-TREE v2.2.5 using GTR+R4 and LG+R4 models, respectively. Individual gene trees were then used for coalescent species-tree inference with ASTRAL v5.7.8[56]. Concatenated nucleotide and protein alignments, generated by catfasta2phyml v2018-09-28 (https://github.com/nylander/catfasta2phyml), served as additional input to IQ-TREE[57] with the same substitution models. *Amborella trichopoda* was specified as the outgroup.

***DensiTree visualization of gene-tree conflict.*** For the focal cloudogram analysis, maximum-likelihood gene trees were restricted to single-copy orthologs containing one representative from each of the eight sampled carnivorous Caryophyllales species and the non-carnivorous relative *Ancistrocladus*. This complete-tip filtering yielded 563 gene trees. Each tree was temporarily outgroup-rooted using *Anicistrocladus*, and taxon labels were standardized so that all trees contained the same tip set and could be displayed in a common tip order. The trees were imported into DensiTree v3.1 without time calibration or ultrametric transformation. Inferred branch lengths from the individual maximum-likelihood trees were retained. The complete set of gene trees was displayed as an overlaid cloudogram together with the consensus topologies calculated by DensiTree, with line intensity reflecting the frequency of coincident or similar tree paths. Variation in root-to-tip depth among trees was retained as gene-tree branch-length heterogeneity and was not interpreted as a common temporal scale. The cloudogram was used only as a qualitative visualization of gene-tree discordance and not as a statistical test distinguishing hybridization from incomplete lineage sorting.

### Supplementary Materials: Figures

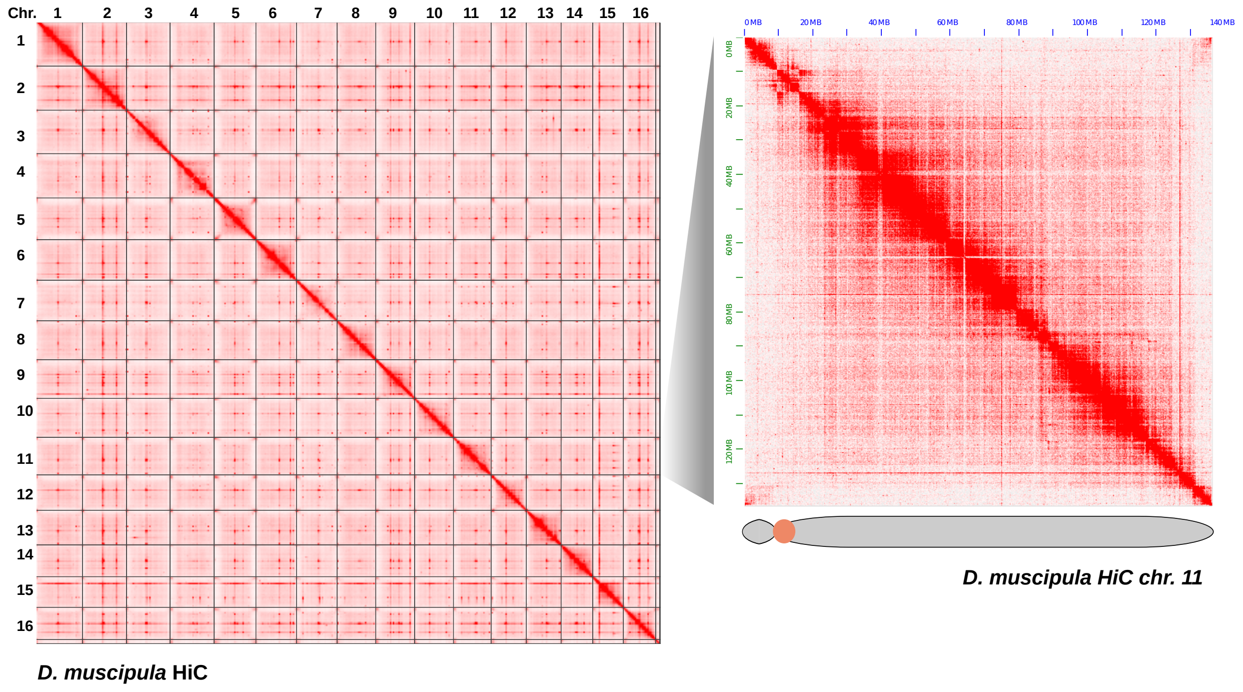


**Supplementary Materials Fig. S1. *Dionaea muscipula* Hi-C contact map.** Hi-C contact map showing every *Dionaea muscipula* chromosome, plus a zoom-in to chromosome 11.

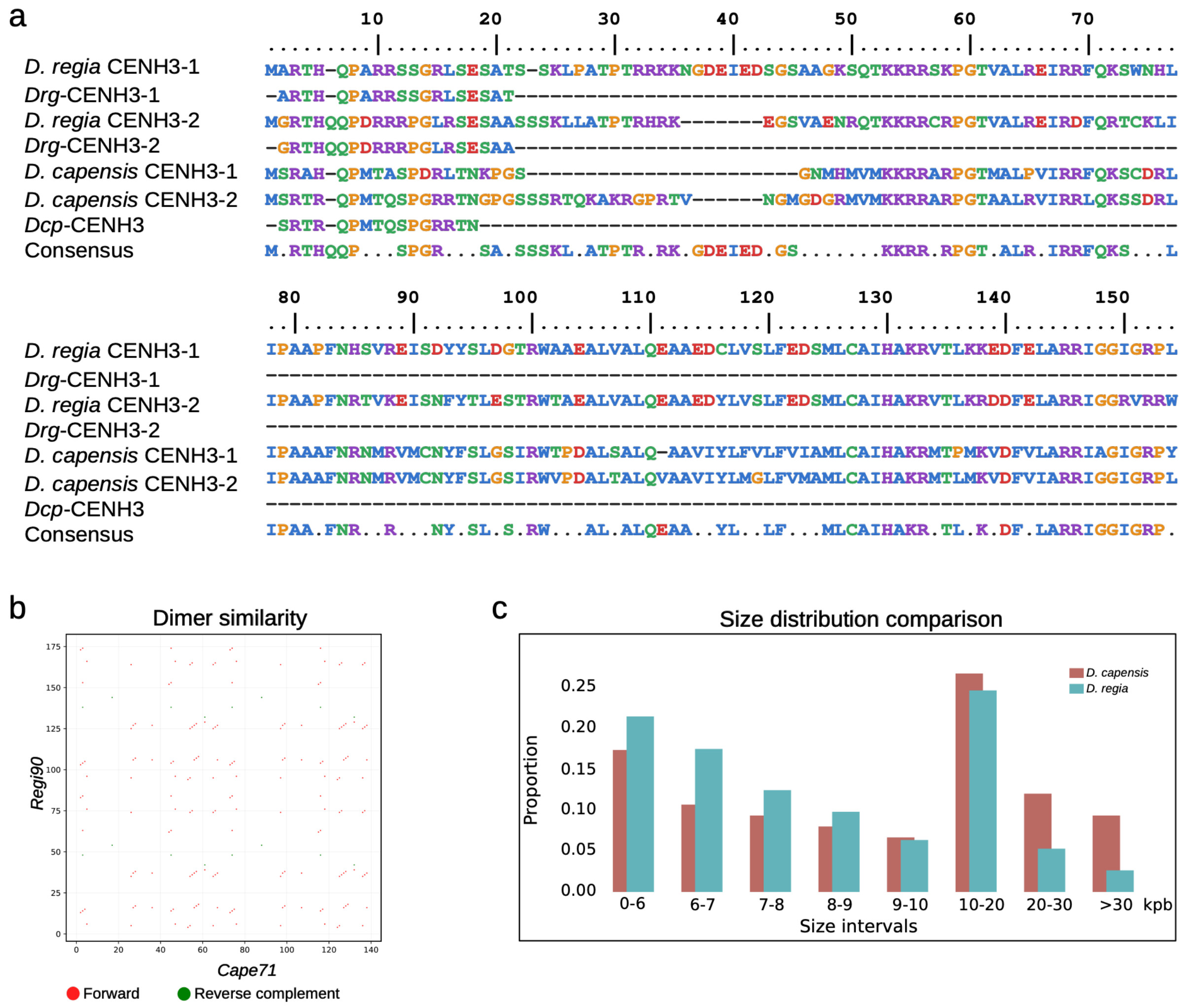


**Supplementary Materials Fig. S2. Characterization of CENH3 sequence and centromeric repeats in *D. regia* and *D. capensis*. (a)** Sequence alignment of inferred CENH3 proteins found in *D. regia* and *D. capensis*. The peptide/epitope used to raise the CENH3 antibody in each species is shown below each CENH3 sequence. **(b)** Dot plot highlighting short stretches of identity between the two satDNA families *Cape71* in *D. capensis* and *Regi90* in *D. regia*. **(c)** Size distribution comparison of *Cape71* and *Regi90*. Notice the prevalent array size of 10-20 kbp for both species.

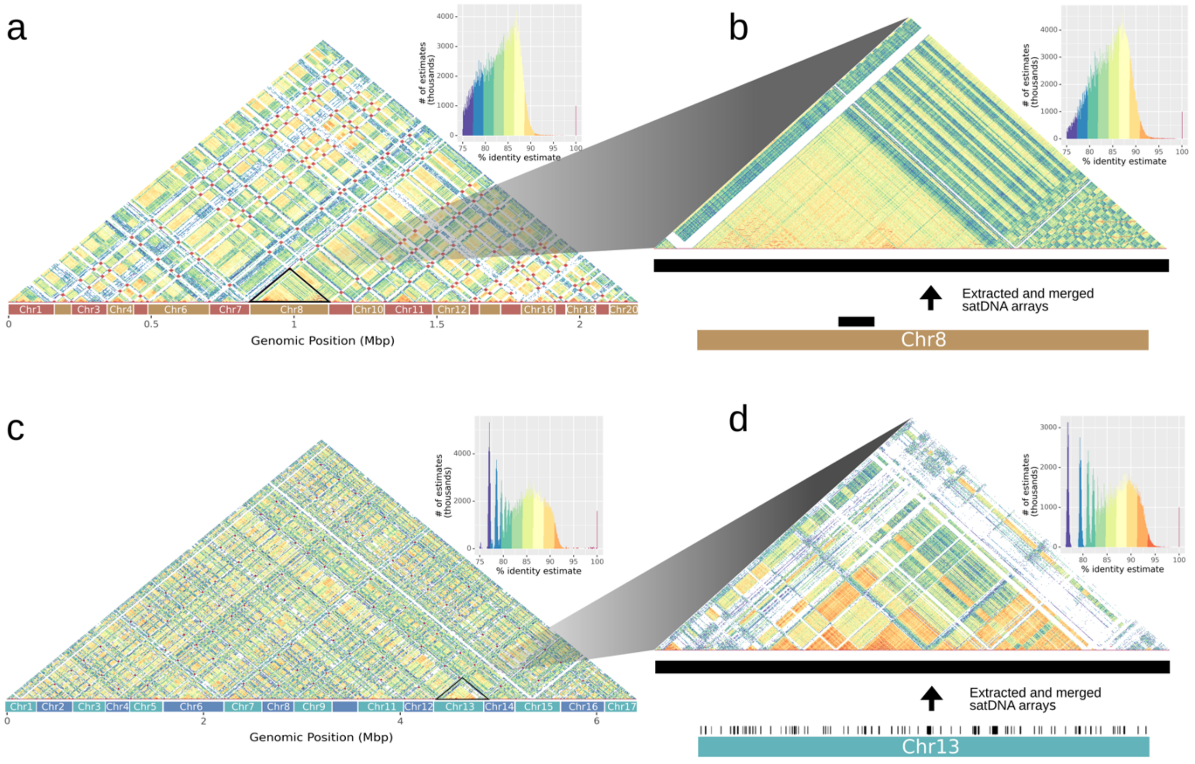


**Supplementary Materials Fig. S3. Comparison of satellite-repeat sequence homogenization in *D. capensis* and *D. regia*.** **(a)** Genome-wide ModDot plot histograms of concatenated isolated satellite arrays of all *D. capensis* chromosomes. **(b)** Close-up view of satellite array homogenization in chromosome 8 centromeric arrays. **(c)** Genome-wide ModDot plot histograms of concatenated isolated *Regi90* arrays of all *D. regia* chromosomes. **(d)** Close-up view of satellite array homogenization in chromosome 13 centromeric arrays.

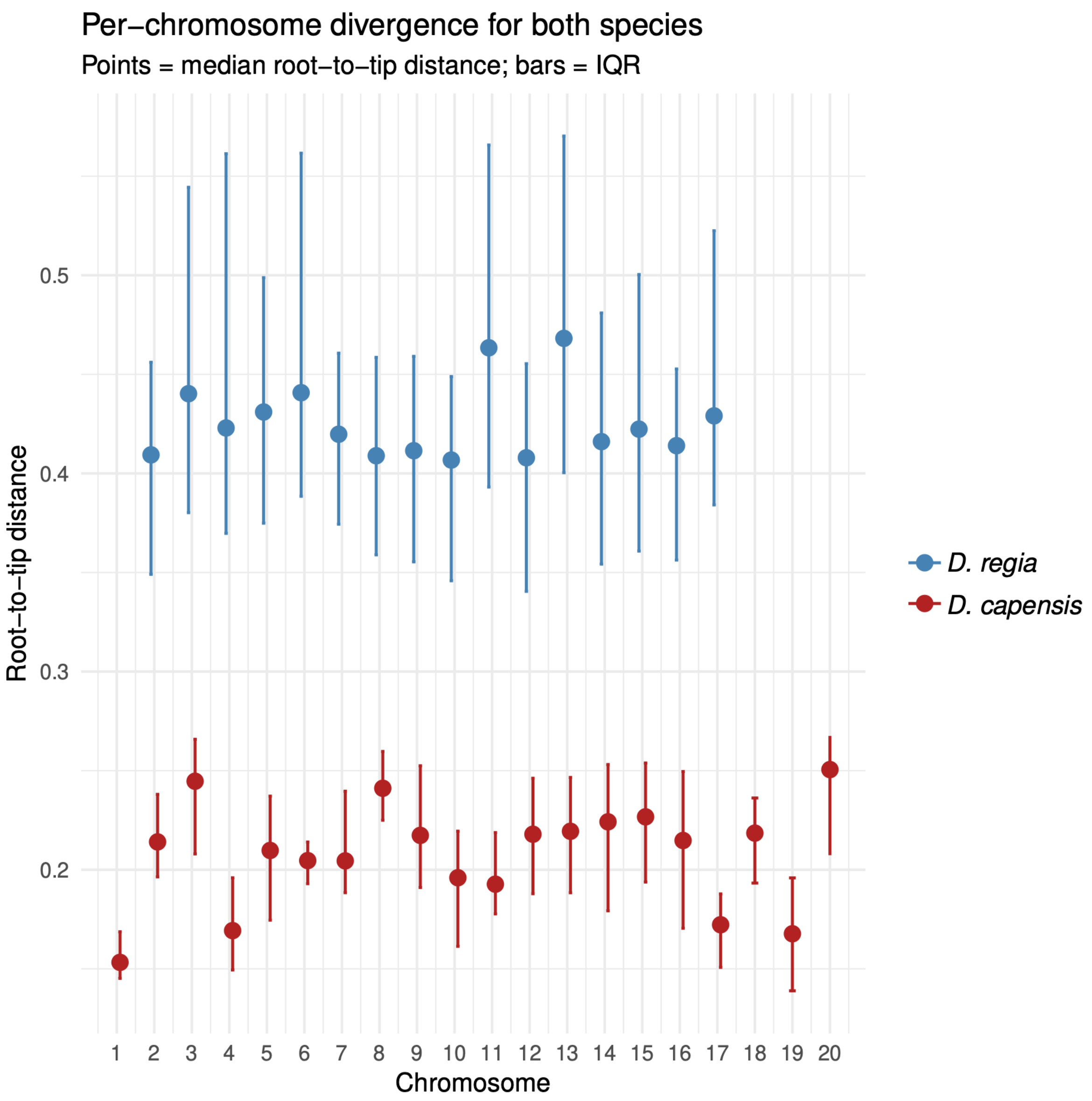


**Supplementary Materials Fig. S4. Per-chromosome divergence of candidate centromere-associated satellite monomers in *D. regia* and *D. capensis*.** Root-to-tip distances were calculated from midpoint-rooted satellite-monomer phylogenies and summarized separately for each pseudochromosome. Plotted along the same y-axis, points show the median root-to-tip distances for monomers assigned to each chromosome, and vertical bars show the interquartile ranges. *Regi90* monomers from *D. regia* show consistently higher root-to-tip distances across pseudochromosomes than *Cape71* monomers from *D. capensis*, whereas *Cape71* values remain lower and more tightly distributed. This chromosome-level comparison supports the pooled root-to-tip analysis in **Fig. 3g** and indicates that the greater divergence of *Regi90* reflects a genome-wide pattern rather than an outlier chromosome or localized repeat subset.

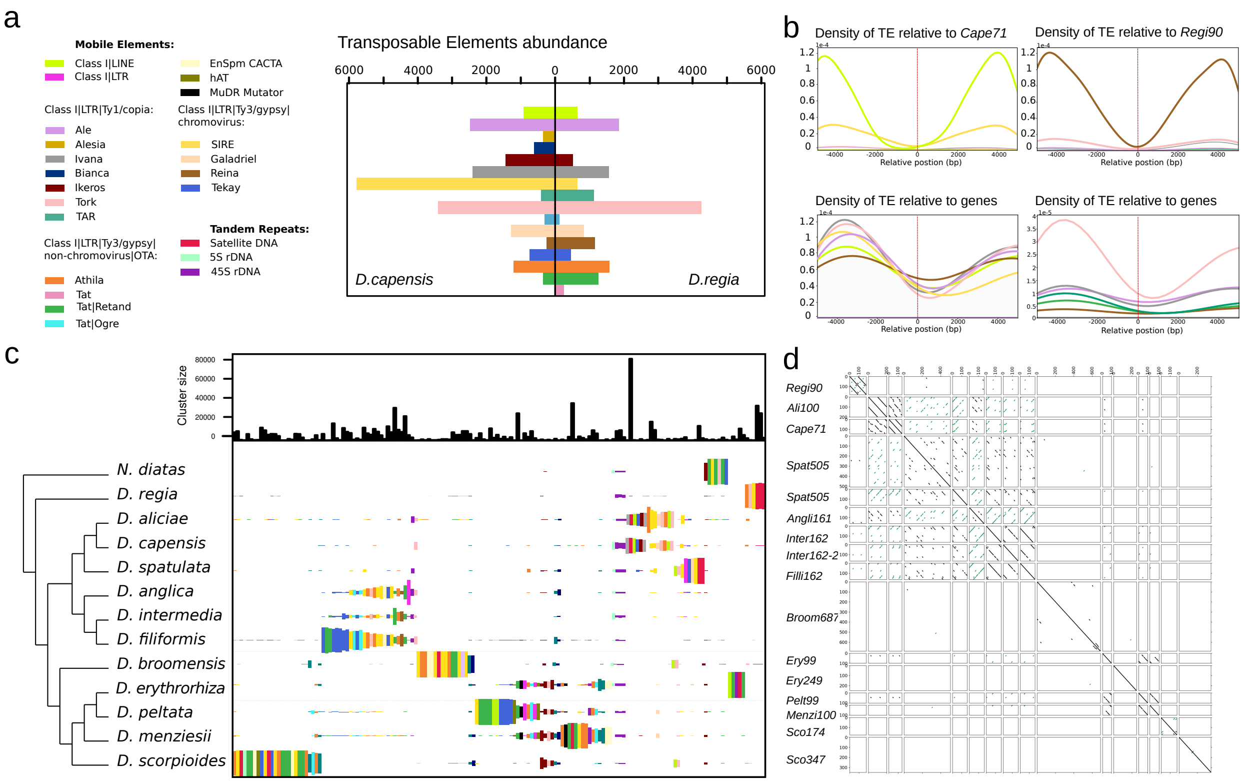


**Supplementary Materials Fig. S5. Repeat composition and interspecific repeatome comparison across *Drosera*.** (**a**) Abundance differences of major transposable-element (TE) lineages between *D. capensis* and *D. regia*. Bar heights indicate the estimated genomic proportion represented by each TE lineage. (**b**) Genomic distribution of selected TE lineages relative to the nearest satellite DNA locus, shown within ±5 kb windows, and relative to genes. (**c**) Comparative repeatome analysis of 12 *Drosera* species and *Nepenthes*. Each column represents an individual repeat cluster recovered by comparative RepeatExplorer clustering, ordered by decreasing total cluster size in the combined dataset. Black bars at the top of the panel indicate the total size of each cluster across the combined analysis. Within each species row, colored bars show the percentage of reads from that species assigned to the corresponding cluster, representing the relative abundance of that cluster in that species. Colors indicate the repeat annotation assigned to each cluster, not a unique cluster identity. Therefore, the same color may appear in multiple columns when several sequence-divergent clusters are annotated as the same TE lineage or repeat class. Adjacent columns with the same or related colors represent separate repeat clusters, often corresponding to divergent subfamilies, lineage-specific variants, or independent repeat expansions. (**d**) Dot-plot comparison of the main satellite DNA families identified among the 12 sampled *Drosera* species.

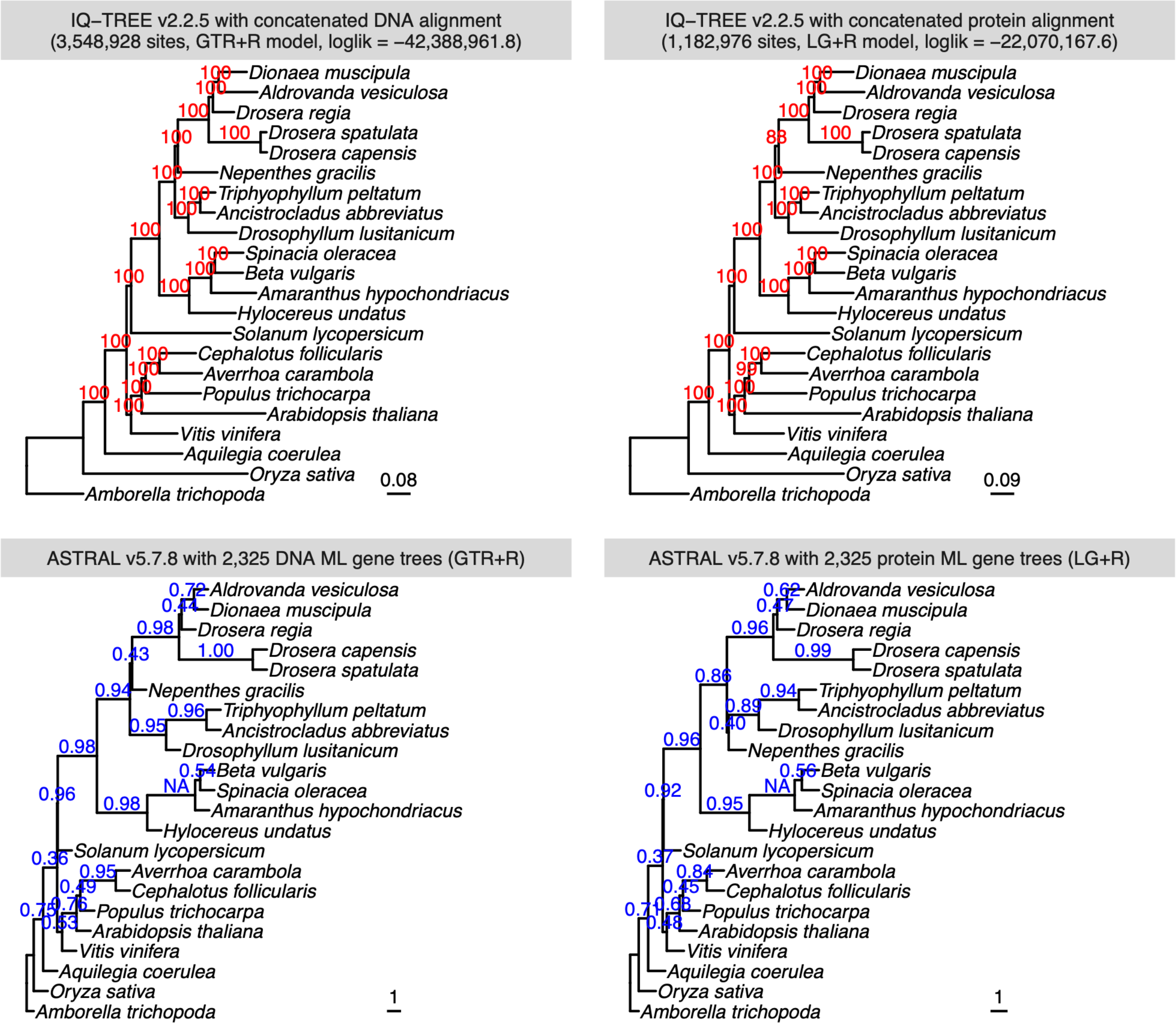
IQ–TREE v2.2.5 with concatenated DNA alignment
(3,548,928 sites, GTR+R model, loglik = –42,388,961.8)
Dionaea muscipula
Aldrovanda vesiculosa
Drosera regia
Drosera spatulata
Drosera capensis
Nepenthes gracilis
Triphyophyllum peltatum
Ancistrocladus abbreviatus
Drosophyllum lusitanicum
Spinacia oleracea
Beta vulgaris
Amaranthus hypochondriacus
Hylocereus undatus
Solanum lycopersicum
Cephalotus follicularis
Averrhoa carambola
Populus trichocarpa
Arabidopsis thaliana
Vitis vinifera
Aquilegia coerulea
Oryza sativa
Amborella trichopoda
0.08
IQ–TREE v2.2.5 with concatenated protein alignment
(1,182,976 sites, LG+R model, loglik = –22,070,167.6)
Dionaea muscipula
Aldrovanda vesiculosa
Drosera regia
Drosera spatulata
Drosera capensis
Nepenthes gracilis
Triphyophyllum peltatum
Ancistrocladus abbreviatus
Drosophyllum lusitanicum
Spinacia oleracea
Beta vulgaris
Amaranthus hypochondriacus
Hylocereus undatus
Solanum lycopersicum
Cephalotus follicularis
Averrhoa carambola
Populus trichocarpa
Arabidopsis thaliana
Vitis vinifera
Aquilegia coerulea
Oryza sativa
Amborella trichopoda
0.09
ASTRAL v5.7.8 with 2,325 DNA ML gene trees (GTR+R)
Aldrovanda vesiculosa
Dionaea muscipula
Drosera regia
Drosera capensis
Drosera spatulata
Nepenthes gracilis
Triphyophyllum peltatum
Ancistrocladus abbreviatus
Drosophyllum lusitanicum
Beta vulgaris
Spinacia oleracea
Amaranthus hypochondriacus
Hylocereus undatus
Solanum lycopersicum
Averrhoa carambola
Cephalotus follicularis
Populus trichocarpa
Arabidopsis thaliana
Vitis vinifera
Aquilegia coerulea
Oryza sativa
Amborella trichopoda
1
ASTRAL v5.7.8 with 2,325 protein ML gene trees (LG+R)
Aldrovanda vesiculosa
Dionaea muscipula
Drosera regia
Drosera capensis
Drosera spatulata
Triphyophyllum peltatum
Ancistrocladus abbreviatus
Drosophyllum lusitanicum
Nepenthes gracilis
Beta vulgaris
Spinacia oleracea
Amaranthus hypochondriacus
Hylocereus undatus
Solanum lycopersicum
Averrhoa carambola
Cephalotus follicularis
Populus trichocarpa
Arabidopsis thaliana
Vitis vinifera
Aquilegia coerulea
Oryza sativa
Amborella trichopoda
1

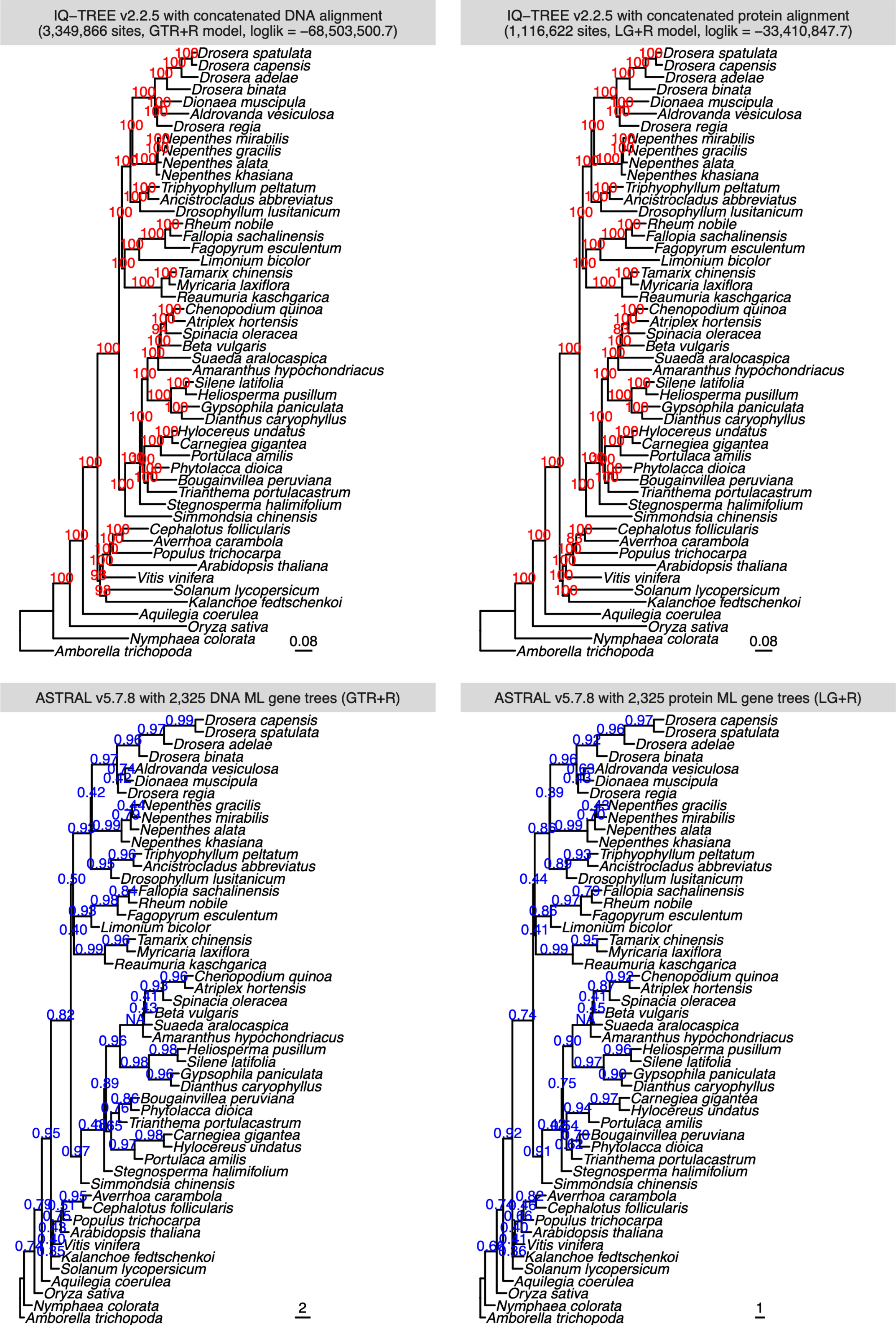
IQ–TREE v2.2.5 with concatenated DNA alignment
(3,349,866 sites, GTR+R model, loglik = –68,503,500.7)
0.08
IQ–TREE v2.2.5 with concatenated protein alignment
(1,116,622 sites, LG+R model, loglik = –33,410,847.7)
0.08
ASTRAL v5.7.8 with 2,325 DNA ML gene trees (GTR+R)
2
ASTRAL v5.7.8 with 2,325 protein ML gene trees (LG+R)
1

**Supplementary Materials Fig. S6. BUSCO gene and species trees.** Two datasets were examined: nucleotide coding sequences or inferred amino acids from 22 species or 50 species. Phylogenetic relationships were reconstructed using two different approaches: (1) maximum likelihood (ML) analysis based on concatenated BUSCO gene alignments (**top**), and (2) ASTRAL species tree inference summarizing gene trees from individual BUSCOs (**bottom**). Both methods consistently placed *D. regia* as the sister group to *Aldrovanda* plus *Dionaea*, rather than to other *Drosera* species. The placement of *Nepenthes* varied among the reconstructions, with some analyses recovering it as sister to *Drosera* plus the snap-trapping lineages, and others placing it with *Drosophyllum* and the *Triphyophyllum-Ancistrocladus* lineage. Notably, the protein-based ASTRAL reconstruction recovered the latter topology. These differences illustrate phylogenetic discordance associated with the alternative datasets, taxonomic sampling, and inference methods.

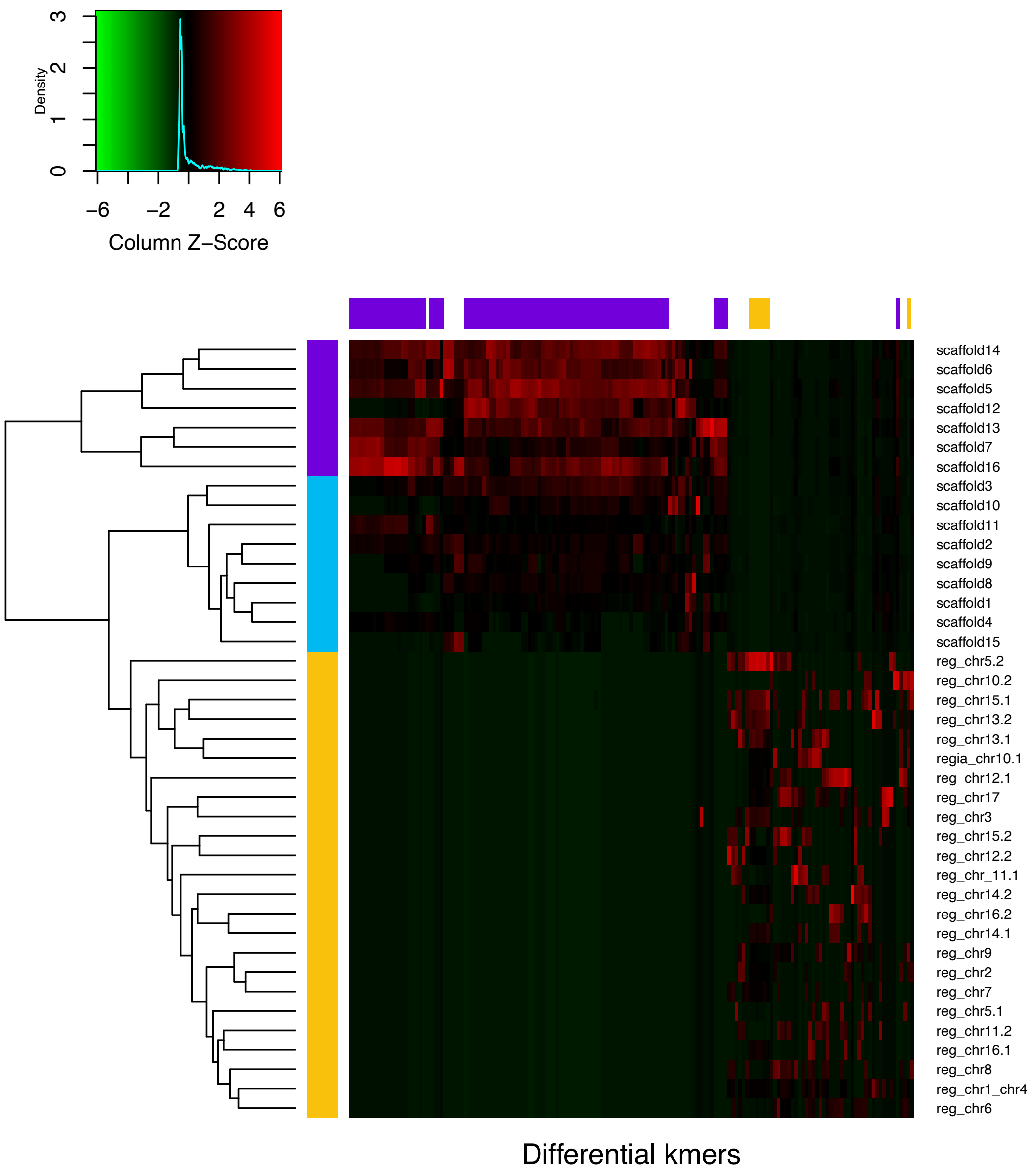


**Supplementary Materials Fig. S7. Subgenome-diagnostic k-mer enrichment in *Dionaea muscipula* and pre-fusion chromosomes of *D. regia*.** Heatmap showing z-scaled relative abundance of SubPhaser diagnostic k-mer windows across chromosomes. For *D. regia*, the inferred pre-fusion chromosomes were used (see ***Synteny analysis***, above). Rows represent chromosomes assigned to inferred genome components; columns represent k-mer windows or k-mer feature groups. The left color bar indicates row-level SubPhaser assignments matching those in main-text **Fig. 5b**: *D. regia* (in yellow) displays no substructuring among its triplet homoeologs, while *Dionaea* is separated into discrete components (sky blue and purple). The top color bar indicates the genome component for which each k-mer window is most diagnostic. Fragmentation and gaps in this top annotation reflect local windows with variable densities of subgenome-diagnostic k-mers, including windows that do not meet assignment thresholds, rather than continuous chromosome-wide subgenome assignment. While *Dionaea* shows two clear chromosome-scale k-mer partitions, only the purple partition contains highly diagnostic k-mers. The sparse *D. regia*-specific signal apparent in the heatmap and top color bar therefore indicates limited local enrichment of *D. regia*-diagnostic k-mers, not absence of *D. regia* sequence or a failed assignment.

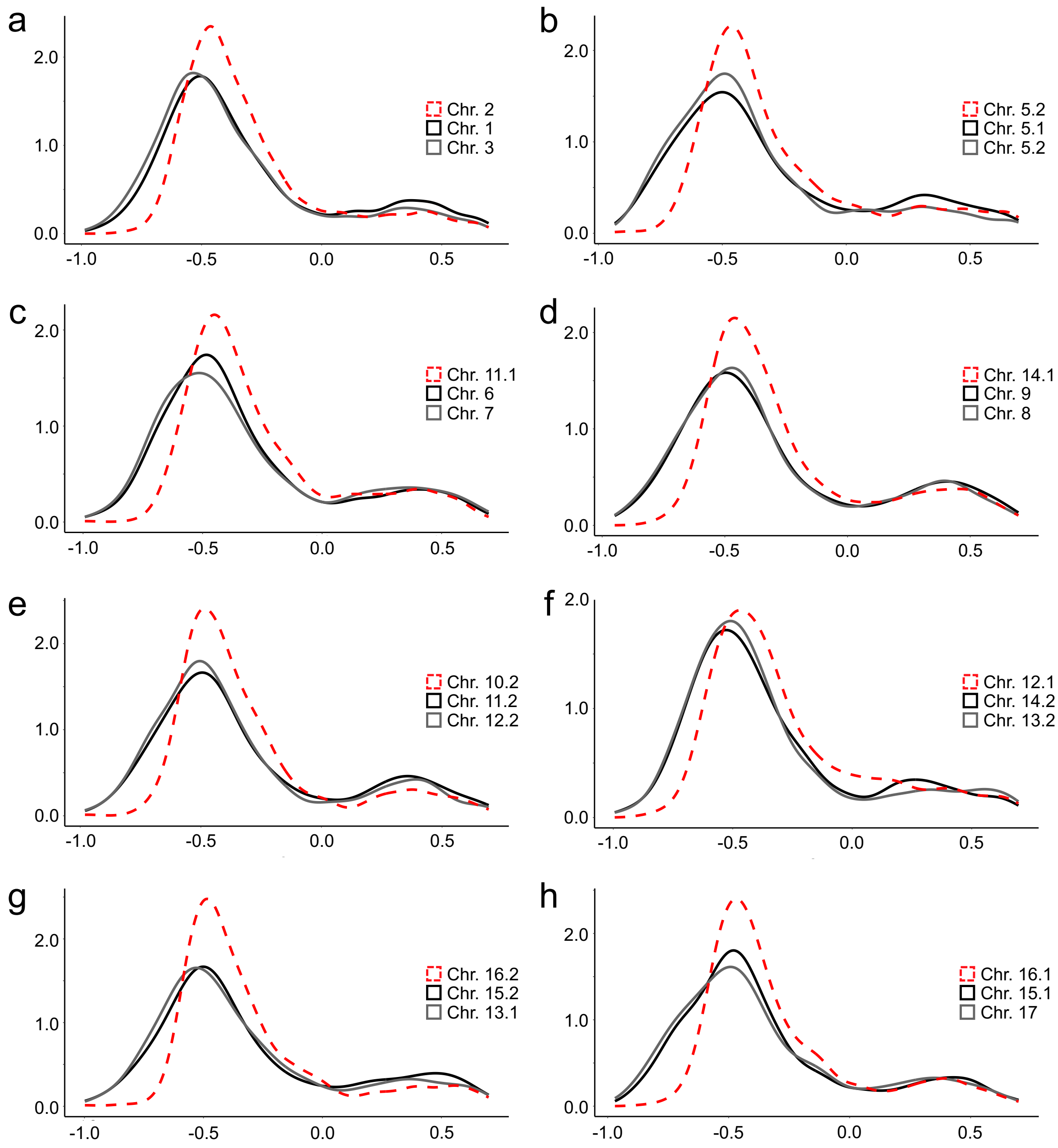


**Supplementary Materials Fig. S8. Ks divergence reveals a 2:1 relationship among the three homoeologous chromosome sets of *D. regia*.** (**a-h**) Ks density plots of homoeologous gene pairs in *D. regia* reveal a clear triplicate subgenomic structure that exhibits a 2:1 configuration of pre-fusion, ancestral homoeologous chromosomes (see ***Synteny analysis***, above). Two of each triplet of homoeologous chromosomes show similar Ks distributions (gray), while the third (dashed red lines) shows higher Ks values, consistent with greater sequence divergence (x-axis = $\log_{10}$ Ks; y-axis = density).

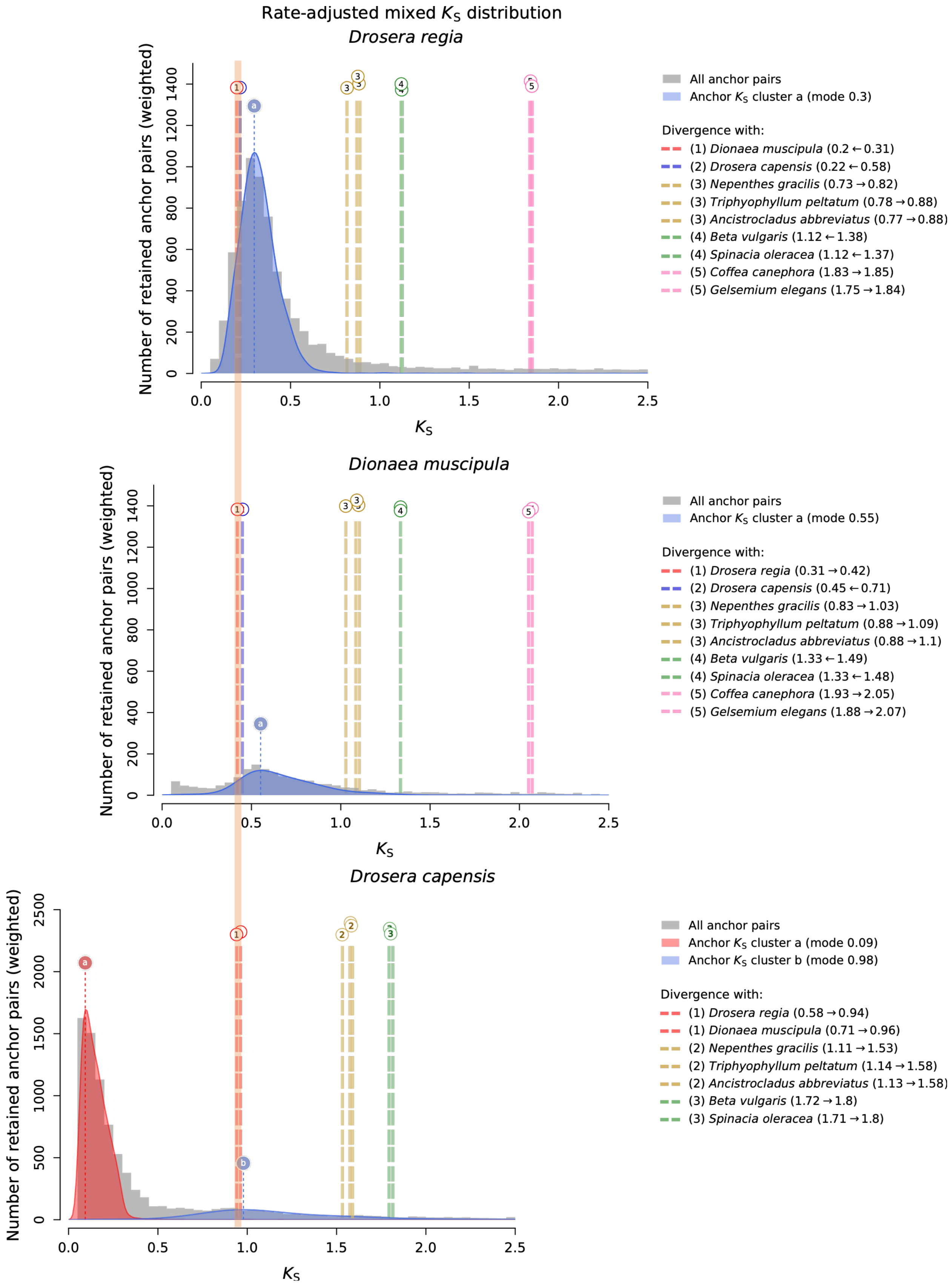

Rate-adjusted mixed $K_S$ distribution
Drosera regia
Number of retained anchor pairs (weighted)
$K_S$
All anchor pairs
Anchor $K_S$ cluster a (mode 0.3)
Divergence with:
(1) Dionaea muscipula (0.2 ← 0.31)
(2) Drosera capensis (0.22 ← 0.58)
(3) Nepenthes gracilis (0.73 → 0.82)
(3) Triphyophyllum peltatum (0.78 → 0.88)
(3) Ancistrocladus abbreviatus (0.77 → 0.88)
(4) Beta vulgaris (1.12 ← 1.38)
(4) Spinacia oleracea (1.12 ← 1.37)
(5) Coffea canephora (1.83 → 1.85)
(5) Gelsemium elegans (1.75 → 1.84)
Dionaea muscipula
All anchor pairs
Anchor $K_S$ cluster a (mode 0.55)
Divergence with:
(1) Drosera regia (0.31 → 0.42)
(2) Drosera capensis (0.45 ← 0.71)
(3) Nepenthes gracilis (0.83 → 1.03)
(3) Triphyophyllum peltatum (0.88 → 1.09)
(3) Ancistrocladus abbreviatus (0.88 → 1.1)
(4) Beta vulgaris (1.33 ← 1.49)
(4) Spinacia oleracea (1.33 ← 1.48)
(5) Coffea canephora (1.93 → 2.05)
(5) Gelsemium elegans (1.88 → 2.07)
Drosera capensis
All anchor pairs
Anchor $K_S$ cluster a (mode 0.09)
Anchor $K_S$ cluster b (mode 0.98)
Divergence with:
(1) Drosera regia (0.58 → 0.94)
(1) Dionaea muscipula (0.71 → 0.96)
(2) Nepenthes gracilis (1.11 → 1.53)
(2) Triphyophyllum peltatum (1.14 → 1.58)
(2) Ancistrocladus abbreviatus (1.13 → 1.58)
(3) Beta vulgaris (1.72 → 1.8)
(3) Spinacia oleracea (1.71 → 1.8)

**Supplementary Materials Fig. S9. *ksrates* synonymous-substitution calibration of lineage splits and polyploidy-associated paralog peaks in *D. regia*, *Dionaea*, and *D. capensis*.** Focal species are shown from top to bottom: *D. regia*, *Dionaea*, and *D. capensis*. Grey histograms show paralogous Ks distributions for each focal genome, and smoothed colored curves mark inferred mixture components corresponding to polyploidy-associated duplicate-gene peaks. Vertical colored bands indicate calibrated orthologous divergence ranges among the focal and outgroup taxa. In the *D. regia* and *Dionaea* panels, the blue duplicate peak corresponds to an older polyploidy signal that is similar in age between these two analyses. This event is interpreted as the *Dionaea*-visible allotetraploidy and as approximately time-coincident with the event contributing to the three-subgenome structure of *D. regia*. In this interpretation, one inferred *D. regia* subgenomic component shows affinity to the *Dionaea* lineage, consistent with the structural evidence for a *Dionaea*-like contribution or retained *Dionaea*-like ancestral component. The orange vertical band marks the calibrated divergence of the focal *Drosera*/*Dionaea* lineages and falls after the older blue duplicate peak in the *D. regia* and *Dionaea* panels. Thus, the duplicate peak predates or is approximately coincident with the deeper lineage history that separates the focal genomes, whereas the species divergences occur later. The *D. capensis* panel appears different because the plotting colors are assigned by the focal-species *ksrates* output rather than by a universal event-color scheme across panels. In the *D. capensis* panel, the prominent low-Ks red component represents the youngest, *D. capensis*-specific duplication, which occurred after the divergence marked by the orange band. The broader blue component in *D. capensis* lies at higher Ks and is therefore older; this is the component most comparable in age to the older blue polyploidy signal visible in the *D. regia* and *Dionaea* panels. Thus, the apparent color mismatch across panels reflects plotting convention and event ordering rather than a biological contradiction: *D. capensis* retains evidence of an older shared or related polyploidy signal, but this signal is visually and biologically complicated by a much stronger, more recent *D. capensis*-specific duplication.

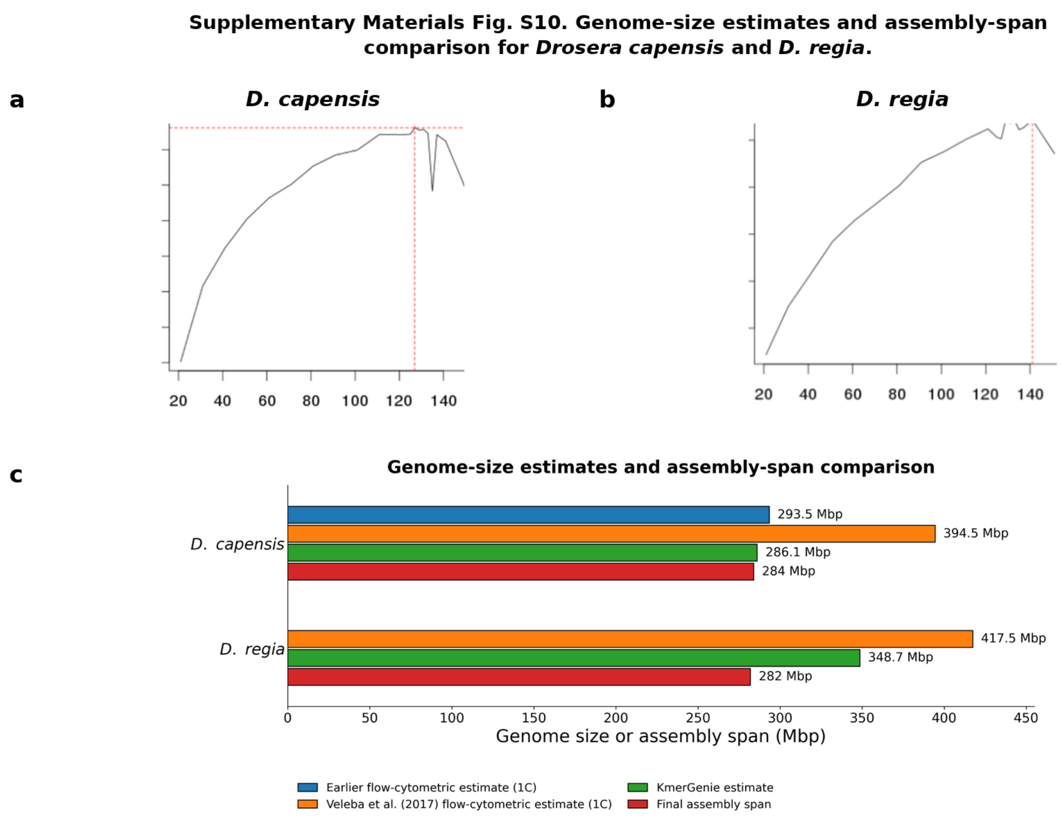


**Supplementary Materials Fig. S10. Genome-size estimates and assembly-span comparison for *D. capensis* and *D. regia*.** (**a**,**b**) KmerGenie genome-size estimates from Illumina short reads for *D. cap*ensis (**a**) and *D. regia* (**b**), with best *k* values of 127 and 141, respectively. Red dashed lines indicate the predicted best k-mer size and corresponding predicted assembly size: 286,140,113 bp for *D. capensis* and 348,732,445 bp for *D. regia*. (**c**) Comparison of an earlier published flow-cytometric 1C estimate cited by Veleba et al.[58], Veleba et al.'s own flow-cytometric 1C estimates, KmerGenie estimates, and final assembly spans. For *D. capensis*, Veleba et al. reported 2C = 789 Mbp and ploidy level 4 (Cx = 197 Mbp), corresponding to 1C = 394.5 Mbp; they also cited an earlier estimate of 2C = 587 Mbp, corresponding to 1C = 293.5 Mbp. For *D. regia*, Veleba et al. reported 2C = 835 Mbp and ploidy level 2, corresponding to 1C = Cx = 417.5 Mbp. The *D. capensis* assembly span agrees closely with the earlier 1C estimate and the KmerGenie estimate, but is smaller than Veleba et al.'s own 1C estimate. The *D. regia* assembly span is smaller than both its flow-cytometric and KmerGenie estimates. These comparisons indicate uncertainty in the expected *D. capensis* genome size and underrepresentation of repetitive or otherwise difficult-to-assemble sequence in *D. regia*.

### Supplementary Materials: Tables

| | BUSCO lineage dataset: eudicots_odb10 | *Drosera capensis* | *Drosera regia* | *Triphyophyllum peltatum* | *Ancistrocladus abbreviatus* | *Dionaea muscipula* |
|---|---|---|---|---|---|---|
| BUSCO | Complete BUSCOs (C) | 2,086 (89.7%) | 2,179 (93.7%) | 2,255 (96.9%) | 2,240 (96.3%) | 2,128 (91.5%) |
| | Complete and single-copy BUSCOs (S) | 1,471 (63.2%) | 1,859 (79.9%) | 2,089 (89.8%) | 2,033 (87.4%) | 1,995 (85.8%) |
| | Complete and duplicated BUSCOs (D) | 615 (26.4%) | 320 (13.8%) | 166 (7.1%) | 207 (8.9%) | 133 (5.7%) |
| | Fragmented BUSCOs (F) | 37 (1.6%) | 27 (1.2%) | 18 (0.8%) | 29 (1.2%) | 37 (1.6%) |
| | Missing BUSCOs (M) | 203 (8.7%) | 120 (5.2%) | 53 (2.3%) | 57 (2.5%) | 161 (6.9%) |
| | Total BUSCO groups searched | 2,326 (100.0%) | 2,326 (100.0%) | 2,326 (100.0%) | 2,326 (100.0%) | 2,326 (100.0%) |
| | Number of scaffolds in final assembly | 1,089 | 1,338 | 429 | 131 | 3,706 |
| | Number of contigs before scaffolding | 1,431 | 3,275 | 468 | 653 | 11,425 |
| | Total length | 284,038,097 | 282,006,236 | 552,408,181 | 1,187,329,241 | 2,551,530,229 |
| | Scaffold N50, final assembly | 12 MB | 15 MB | 26 MB | 64 MB | 144 MB |
| | Contig N50, before scaffolding | 1 MB | 287 KB | 14 MB | 4 MB | 432 KB |

| | BUSCO lineage dataset: embryophyta_odb10 | *Drosera capensis* | *Drosera regia* | *Triphyophyllum peltatum* | *Ancistrocladus abbreviatus* | *Dionaea muscipula* |
|---|---|---|---|---|---|---|
| BUSCO | Complete BUSCOs (C) | 1,530 (94.8%) | 1,561 (96.7%) | 1,578 (97.8%) | 1,572 (97.4%) | 1,540 (95.4%) |
| | Complete and single-copy BUSCOs (S) | 1,115 (69.1%) | 1,390 (86.1%) | 1,507 (93.4%) | 1,459 (90.4%) | 1,466 (90.8%) |
| | Complete and duplicated BUSCOs (D) | 415 (25.7%) | 171 (10.6%) | 71 (4.4%) | 113 (7.0%) | 74 (4.6%) |
| | Fragmented BUSCOs (F) | 23 (1.4%) | 11 (0.7%) | 15 (0.9%) | 18 (1.1%) | 25 (1.5%) |
| | Missing BUSCOs (M) | 61 (3.8%) | 42 (2.6%) | 21 (1.3%) | 24 (1.5%) | 49 (3.0%) |
| | Total BUSCO groups searched | 1,614 (100.0%) | 1,614 (100.0%) | 1,614 (100.0%) | 1,614 (100.0%) | 1,614 (100.0%) |
| | Number of scaffolds in final assembly | 1,089 | 1,338 | 429 | 131 | 3,706 |
| | Number of contigs before scaffolding | 1,431 | 3,275 | 468 | 653 | 11,425 |
| | Total length | 284,038,097 | 282,006,236 | 552,408,181 | 1,187,329,241 | 2,551,530,229 |
| | Scaffold N50, final assembly | 12 MB | 15 MB | 26 MB | 64 MB | 144 MB |
| | Contig N50, before scaffolding | 1 MB | 287 KB | 14 MB | 4 MB | 432 KB |

| | | | | | | |
|---|---|---|---|---|---|---|
| QUAST | Number of sequences (scaffolds) | 1,089 | 1,338 | 429 | 131 | 3,706 |
| | Largest scaffold | 17,145,115 | 20,363,681 | 31,991,434 | 85,057,085 | 169,349,142 |
| | Total scaffold-assembly length | 284,038,097 | 282,006,236 | 552,408,181 | 1,187,329,241 | 2,551,530,229 |
| | GC content (%) | 37 | 37 | 40 | 36 | 44 |
| | Scaffold N50 | 12,650,043 | 15,761,547 | 26,241,775 | 64,216,489 | 144,508,999 |
| | Scaffold N75 | 10,814,855 | 12,243,869 | 24,909,253 | 58,859,448 | 126,729,089 |
| | Scaffold L50 | 10 | 8 | 10 | 9 | 9 |
| | Scaffold L75 | 16 | 13 | 16 | 14 | 13 |
| | Ns per 100 kbp | 34 | 120 | 1 | 4 | 30 |

**Supplementary Materials Table S1. Assembly-quality assessment of the genomes.** BUSCO completeness was evaluated using both the eudicots_odb10 and embryophyta_odb10 lineage datasets; percentages cited in the main text and response letter refer to the embryophyta_odb10 results. Pre-scaffolding contig counts and contig N50 values are reported separately from final scaffold-level statistics. QUAST was run on the final scaffold-level assemblies; therefore, the QUAST sequence count, largest-sequence value, N50, N75, L50, and L75 statistics refer to scaffolds.

| Species | Total reads analyzed | Read length (bp)[1] | Reads: repetitive elements | % reads repetitive | Data source | Raw-data accession(s) |
|---|---|---|---|---|---|---|
| *Nepenthes diatas* | 236096 | 150 | 78331 | 33.18 | Generated in this study | ERR18101751 |
| *D. regia* | 235232 | 251 | 161825 | 68.79 | Generated in this study | ERR18094178 |
| *D. aliciae* | 236124 | 251 | 156822 | 66.42 | Generated in this study | ERR18094174 |
| *D. capensis* | 235410 | 251 | 178862 | 75.98 | Generated in this study | ERR18094177 |
| *D. spatulata* | 235912 | 145 | 172730 | 73.22 | Public data; processed in this study | PRJDB9009 |
| *D. anglica* | 235654 | 101 | 136961 | 58.12 | Public data; processed in this study | ERR5555107 |
| *D. intermedia* | 235694 | 151 | 203669 | 86.41 | Public data; processed in this study | SRR12432545 |
| *D. filiformis* | 234720 | 251 | 203766 | 86.81 | Generated in this study | ERR18094168; ERR18094169 |
| *D. broomensis* | 235806 | 251 | 158050 | 67.03 | Generated in this study | ERR18094175 |
| *D. erythrorhiza* | 235186 | 151 | 113059 | 48.07 | Public data; processed in this study | SRR7072766; SRR7072767 |
| *D. peltata* | 235214 | 251 | 195065 | 82.93 | Generated in this study | ERR18094170; ERR18094171 |
| *D. menziesii* | 235850 | 251 | 184643 | 78.29 | Generated in this study | ERR18094172; ERR18094173 |
| *D. scorpioides* | 234890 | 251 | 172536 | 73.45 | Generated in this study | ERR18094176 |

**Supplementary Materials Table S2. Short reads analyzed in the repeatome comparative analysis.** Raw sequencing datasets generated in this study are deposited in the European Nucleotide Archive under **BioProject PRJEB128437**; previously published datasets are indicated by their corresponding public accession numbers.

**Supplementary Note 1: RNA-seq evidence for transcriptional and structural integrity of kinetochore-associated genes in *D. capensis* and *D. regia***

To assess whether the kinetochore-associated targets examined in our cytological analyses are present and transcribed in both species, we evaluated RNA-seq assemblies, genome-aligned read coverage, predicted open reading frames (ORFs), epitope recovery, and transcript abundance for *KNL1*, *CENH3*, and *NDC80* (**Supplementary Note 1 Table 1**; **Supplementary Note 1 Figs. 1-3**). These analyses were intended to distinguish simple gene absence from more limited cases of transcript fragmentation, ORF disruption, or chimeric transcription.

***KNL1***. In *D. capensis*, *KNL1* is recovered as a continuous transcript encoding a full-length ORF consistent in size and overall architecture with reference orthologs (**Supplementary Note 1 Fig. 1A**). RNA-seq reads show coverage across the coding region, and the antibody epitope is represented within the assembled transcript (**Supplementary Note 1 Fig. 1B**; **Supplementary Note 1 Table 1**). Together, these data indicate that *KNL1* is intact and expressed in *D. capensis*.

In *D. regia*, *KNL1* is also clearly transcribed, with a summed transcript abundance higher than that recovered in *D. capensis* (**Supplementary Note 1 Table 1**). However, Trinity reconstruction yielded multiple transcript fragments rather than a single contiguous ORF. These fragments align to non-overlapping conserved portions of *KNL1* orthologs and collectively span much of the expected protein length (**Supplementary Note 1 Fig. 1A**). RNA-seq reads map robustly to the recovered fragments, including the epitope-encoding region (**Supplementary Note 1 Fig. 1C**), but only limited bridging read support was recovered between fragments. We therefore interpret *KNL1* in *D. regia* as expressed, but with full-length transcript reconstruction remaining ambiguous.

Overall, the RNA evidence supports the presence and transcription of *KNL1* in both species, while indicating that transcript reconstruction is substantially more fragmented in *D. regia* than in *D. capensis*.

***CENH3* paralogs**. Both species contain two *CENH3* loci, but the two copies are not equally supported as intact genes (**Supplementary Note 1 Table 1**; **Supplementary Note 1 Fig. 2**).

In *D. capensis*, *CENH3-2* encodes a full-length protein with a conserved histone fold domain and is transcriptionally supported (**Supplementary Note 1 Fig. 2A, B**). By contrast, *CENH3-1* contains a premature stop codon that truncates the predicted protein before completion of the canonical conserved region, consistent with compromised or absent function (**Supplementary Note 1 Fig. 2A**). Although expression evidence for this locus is weak, RNA-seq coverage is present in the corresponding genomic region (**Supplementary Note 1 Fig. 2C**), indicating that the locus is transcribed at least at low level. We therefore interpret *D. capensis CENH3-2* as the clearly intact and expressed paralog, whereas *CENH3-1* appears to be structurally compromised.

In *D. regia*, *CENH3-1* is recovered as a full-length, transcriptionally supported paralog with a conserved histone fold domain (**Supplementary Note 1 Fig. 2A, D**; **Supplementary Note 1 Table 1**). In contrast, *CENH3-2* is recovered as a chimeric transcript, in which the *CENH3*-like coding region occurs as a downstream ORF. The CENH3-homologous portion aligns to orthologs, but RNA-seq reads support the fusion structure itself and no separate monocistronic *CENH3* transcript was detected (**Supplementary Note 1 Fig. 2E**). Thus, although transcription is evident, the transcript structure is inconsistent with straightforward production of a canonical CENH3 protein from this paralog.

Taken together, these results indicate that in each species only one *CENH3* paralog is clearly structurally intact, while the second shows features consistent with compromised functionality: premature truncation in *D. capensis* and chimeric transcript structure in *D. regia*.

***NDC80***. *NDC80* is supported as present and transcribed in both species, although transcript reconstruction differs between them (**Supplementary Note 1 Table 1**; **Supplementary Note 1 Fig. 3**).

In *D. capensis*, Trinity reconstruction fragments the *NDC80* transcript across multiple contigs and incompletely recovers the epitope-containing region in the transcript assembly. However, genome-aligned RNA-seq reads span the relevant genomic interval (**Supplementary Note 1 Fig. 3B**), indicating that the epitope-encoding region is transcribed despite incomplete transcript assembly. Together with the overall ORF structure inferred from genome-based evidence (**Supplementary Note 1 Table 1**), these data support the interpretation that *NDC80* is present and expressed in *D. capensis*, and that the apparent deficiency reflects transcript misassembly rather than true absence of the gene.

***Conclusions***. Across all three loci, the RNA-seq evidence argues against a simple explanation based on wholesale loss of kinetochore-associated genes in either species. Instead, the two *Drosera* genomes differ mainly in the quality and structure of transcript recovery. *KNL1* and *NDC80* are transcriptionally supported in both taxa, although *KNL1* in *D. regia* and *NDC80* in *D. capensis* are affected by transcript fragmentation. For *CENH3*, both species retain two paralogs, but only one copy in each species is clearly intact, whereas the second shows features consistent with reduced or uncertain functionality. These results provide a gene-level framework for interpreting the cytological observations: the contrasting immunostaining patterns, including weak or absent *KNL1* and *CENH3* signal in *D. regia* and robust signal in *D. capensis*, are not readily explained by gene absence or lack of transcription, and instead point to differences in protein integrity, epitope accessibility, localization, abundance, or centromere-associated chromatin context.

### Supplementary Note 1: Figures and Tables

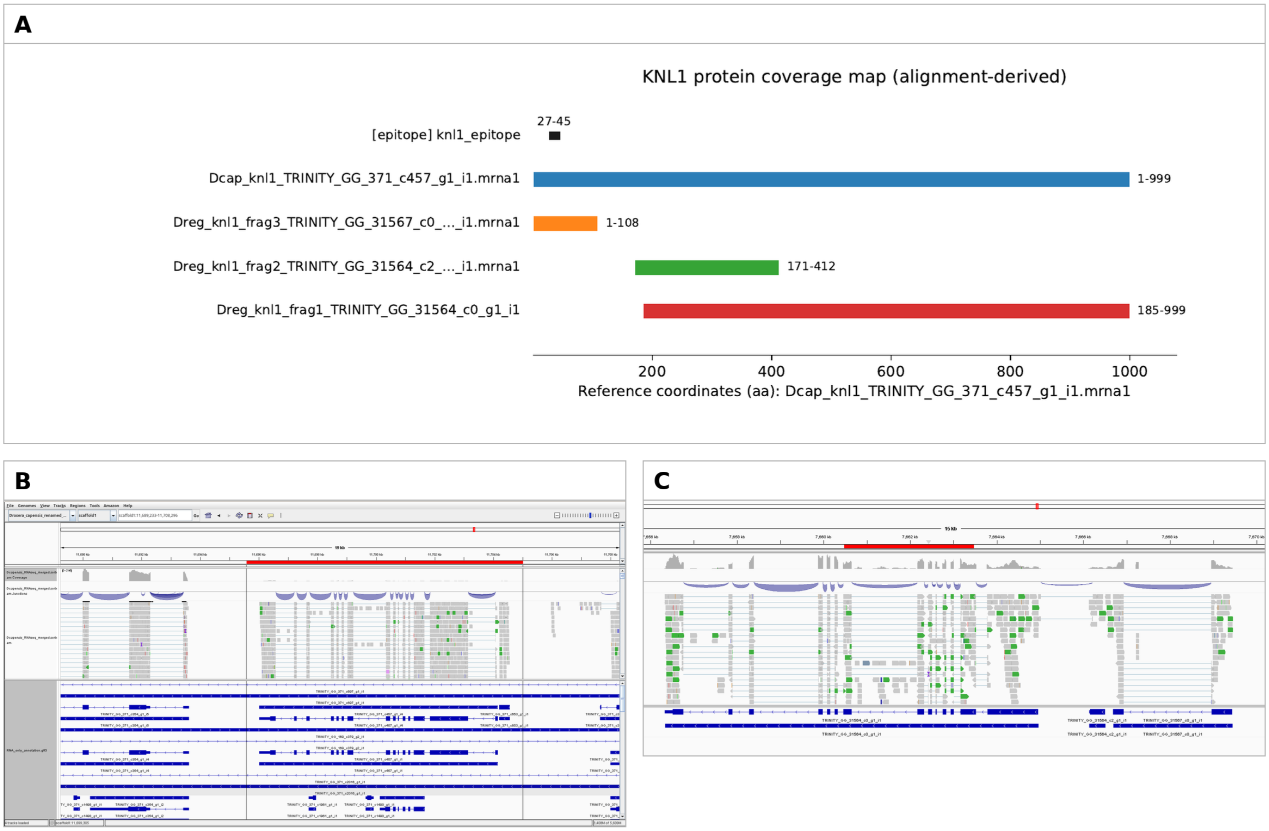


**Supplementary Note 1 Fig. 1. RNA-seq evidence for *KNL1* in *D. capensis* and *D. regia*.** (**A**) Protein-level comparison and coverage summary for *KNL1*, showing recovery of a continuous full-length ORF in *D. capensis* and multiple non-overlapping transcript fragments in *D. regia* that together span much of the expected protein length. (**B**) IGV[59] view of the *KNL1* locus in *D. capensis*, showing RNA-seq read coverage across the coding region and support for the epitope-containing interval. (**C**) IGV view of *KNL1*-associated transcript fragments in *D. regia*, showing RNA-seq support across recovered fragments, including the epitope-encoding region, despite incomplete full-length transcript reconstruction.

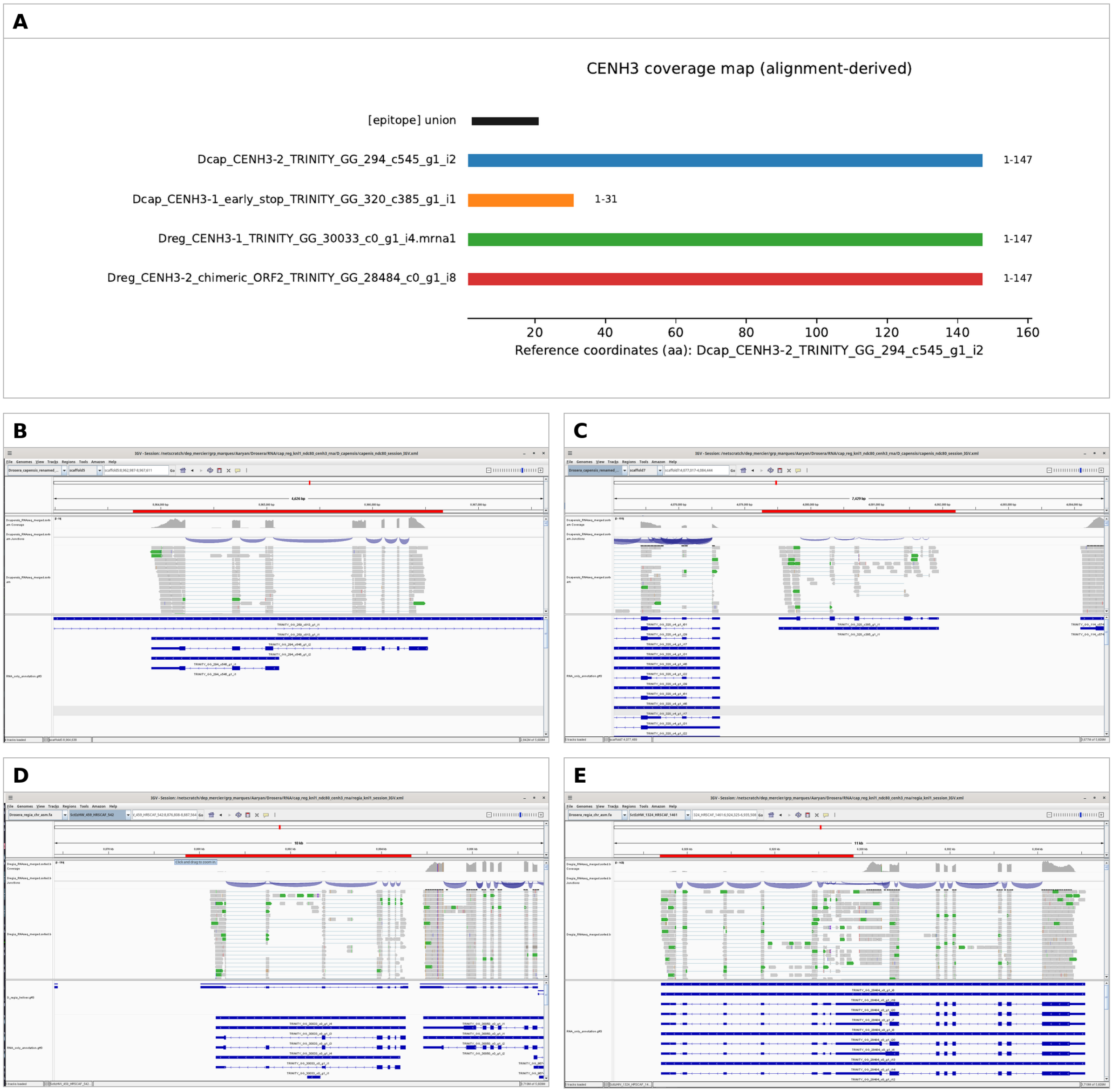


**Supplementary Note 1 Fig. 2. RNA-seq and structural evidence for *CENH3* paralogs in *D. capensis* and *D. regia*.** (**A**) Alignment-based comparison of predicted CENH3 proteins, highlighting conserved regions and structural differences among paralogs. (**B**) IGV view of *D. capensis CENH3-2*, the intact and expressed paralog. (**C**) IGV view of *D. capensis CENH3-1*, showing transcriptional support for a paralog containing an early stop codon and therefore predicted to be structurally compromised. (**D**) IGV view of *D. regia CENH3-1*, showing transcriptional support for a full-length paralog. (E) IGV view of *D. regia CENH3-2*, showing read support for a chimeric transcript in which the *CENH3*-like ORF occurs downstream within a fused transcript structure. Together, these data indicate that only one *CENH3* paralog in each species is clearly intact.

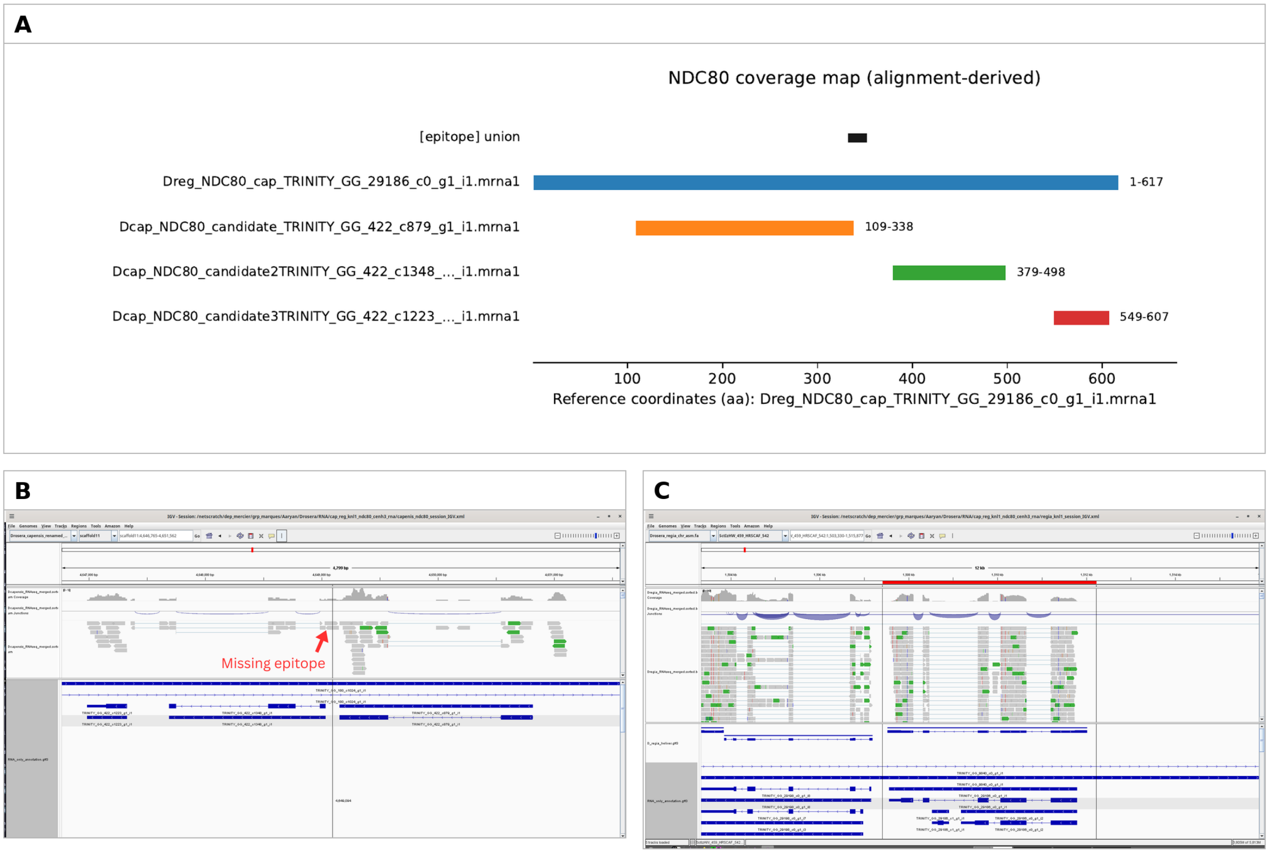


**Supplementary Note 1 Fig. 3. RNA-seq evidence for *NDC80* in *D. capensis* and *D. regia*.** (**A**) Alignment-based and transcript-structure summary for *NDC80*, showing recovery of a full-length ORF in *D. regia* and fragmented transcript reconstruction in *D. capensis*. (**B**) IGV view of the *NDC80* locus in *D. capensis*, showing genome-aligned RNA-seq reads spanning the epitope-encoding region despite incomplete Trinity reconstruction. (**C**) IGV view of the *NDC80* locus in *D. regia*, showing RNA-seq support across a single full-length transcript containing the epitope region.

| Gene | Species | Copy | Assembly status | Intact ORF? | Transcribed? | TPM | Epitope | Interpretation |
|---|---|---|---|---|---|---|---|---|
| *KNL1* | *capensis* | - | assembled | yes | yes | 1.87 | yes | intact and expressed |
| *KNL1* | *regia* | - | fragmented | uncertain | yes | 5.75* | yes | expressed; full-length reconstruction ambiguous |
| *CENH3* | *capensis* | 1 | assembled | no; early stop | yes, very low | 0.00† | yes | transcribed but structurally compromised |
| *CENH3* | *capensis* | 2 | assembled | yes | yes | 0.46 | yes | intact and expressed |
| *CENH3* | *regia* | 1 | assembled | yes | yes | 5.75 | yes | intact and expressed |
| *CENH3* | *regia* | 2 | chimeric | no; canonical functional ORF unlikely | yes | 0.85 | yes | chimeric transcript; canonical CENH3 translation unlikely |
| *NDC80* | *capensis* | - | fragmented; epitope not fully recovered in Trinity | likely yes | yes | 1.19 | yes, supported by genome-mapped reads | expressed; likely intact despite transcript misassembly |
| *NDC80* | *regia* | - | single contig | yes | yes | 8.33 | yes | intact and expressed |

* Summed across recovered transcript fragments.
† TPM was reported as 0.00, but RNA-seq coverage is visible in the corresponding IGV view, indicating at least low-level transcription.

**Supplementary Note 1 Table 1. Summary of RNA-seq support, ORF integrity, and epitope recovery for *KNL1*, *CENH3*, and *NDC80* in *D. capensis* and *D. regia*.** For each locus or paralog, the table summarizes assembly status, predicted ORF integrity, transcriptional support, transcript abundance, recovery of the antibody epitope region, associated IGV and alignment evidence, and the resulting interpretation. For *CENH3*, each paralog is listed separately because the two copies differ markedly in transcript structure and inferred functional integrity.

**Methods for Supplementary Note 1**

For *D. capensis* analyses, we used the transcriptome described above under **Transcriptome sequencing and genome annotation**, which was otherwise used for genome annotation. For *D. regia* analyses, we extracted total RNA of *D. regia* using the Spectrum™ Plant Total RNA Kit (Sigma-Aldrich). The RNA-seq library was sequenced on a NextSeq 2000 instrument, for a total of 31,650,905 read pairs. RNA-seq reads were trimmed with fastp, aligned to the reference genomes with STAR, and assembled in genome-guided mode with Trinity. For locus-level inspection, reads were also mapped to the genomes with HISAT2 and visualized as sorted/indexed BAM files. Trinity transcripts were aligned back to the genomes with GMAP, and RNA-supported gene models were reconstructed using PASA. Predicted proteins were extracted with gffread, and manually curated transcripts were translated with TransDecoder. Candidate *KNL1*, *CENH3*, and *NDC80* loci were identified by BLAST searches against Trinity/PASA-derived transcript and protein sets using reference proteins, previously predicted *Drosera* sequences, and antibody epitopes as queries. Expression was quantified with Salmon (-l A --validateMappings) and reported as TPM.

**Supplementary Note 2: Extended structural and topological analyses of higher-order genome history in *Drosera regia*, *D. capensis*, and *Dionaea muscipula***

***Multispecies syntenic block framework.*** All structural analyses described here were based on a multispecies syntenic block coordinate table generated from a comparative GENESPACE-based synteny framework. In this table, collinear blocks descending from a shared ancestral genomic architecture were projected into a common reference namespace defined by the eight-chromosome dominant subgenome of *Nepenthes gracilis*. In this context, the dominant *Nepenthes* subgenome serves as an operational proxy for the ancestral karyotypic framework of the clade under study. Each row of the table records a pairwise mapping between this reference and one extant genome included in the analysis (*Ancistrocladus abbreviatus*, *Dionaea muscipula*, *Drosera capensis*, *Drosera regia*, or *Triphyophyllum peltatum*), including genome identities on both sides, chromosome identities, block order along chromosomes, genomic spans, and block support. This representation makes it possible to compare chromosome architecture among taxa within a shared label space even when the focal genomes differ substantially in ploidy, chromosome structure, and annotation completeness. The framework is particularly useful here because it captures the ordered distribution of inferred ancestral compartments along extant chromosomes while remaining independent of direct gene-order reconstruction in each focal lineage.

The framework sdescribed above separates broad syntenic relatedness from higher-order structural equivalence. Two genomes may share many of the same ancestral compartments and still differ substantially in the way those compartments were assembled into present-day chromosome-scale paths. By placing all taxa into the same dominant-*Nepenthes* scaffold label space, the block table permits direct comparison not only of block content, but also of (1) adjacency structure, (2) ordered run paths, (3) local motifs, (4) weighted breakpoint graphs, and (5) chromosome-level feature vectors. Definitions used are:

1. *Adjacency* is defined at the level of inferred syntenic block runs rather than individual genes. After ordering block-table entries along each chromosome and collapsing consecutive entries with the same dominant reference scaffold label into a single run, each transition between neighboring runs with different labels is recorded as one adjacency. Adjacencies therefore represent joins between distinct ancestral block compartments along extant chromosomes.

2. A *run* is defined as a contiguous block-table segment along a chromosome in which adjacent entries share the same dominant ancestral scaffold label; adjacent entries with identical labels are collapsed into one run.

3. *Weighted motifs* are defined as short, ordered sub-sequences of consecutive labeled runs along a chromosome, typically spanning two or three adjacent runs, in which each motif is assigned a weight based on the support associated with its constituent runs rather than being treated as a simple binary occurrence. Weighted motif comparisons therefore capture not only the recurrence of local structural patterns, but also the extent to which those patterns are supported by larger or more strongly represented segments.

4. *Weighted breakpoint graphs* are constructed by representing each distinct ancestral scaffold label as a *node* and each observed adjacency between neighboring labeled runs along extant chromosomes as an *edge*. Edge weights record the strength of support for each adjacency, such as its frequency, cumulative support, or cumulative span. These graphs therefore

summarize higher-order chromosome structure as a weighted network of inferred joins among ancestral block compartments.

5. *Chromosome-level feature vectors* are constructed by summarizing each chromosome as a multivariate numerical profile derived from its ordered sequence of labeled runs in the shared reference scaffold space. Here, a vector denotes an ordered list of quantitative feature values, with each position in the list corresponding to one predefined chromosome property. Features include chromosome-wide quantities: run number, total support, and total span, together with normalized ancestral-label composition and adjacency-transition frequencies. These vectors provide a common structural feature space for comparing chromosomes across taxa and for identifying inferred structural channels. Here, a *channel* denotes a data-derived structural grouping recovered from chromosome-level feature space; it may serve as a proxy for a major subgenomic component, but is not assumed a priori to represent a fully validated biological subgenome.

We used the multispecies syntenic block coordinate table to test whether *D. regia* and *D. capensis* share a specific higher-order structural history, rather than merely broad macrosyntenic continuity. For all downstream analyses, rows linking a focal extant genome to the dominant *Nepenthes* subgenome were extracted and ordered by chromosome and ordinal midpoint. Consecutive rows carrying the same dominant-*Nepenthes* scaffold label were then collapsed into contiguous runs. Each run therefore represents a structurally coherent chromosome segment from a given extant genome that maps consistently to one reference scaffold label. These run tables formed the common substrate for all subsequent analyses. In practice, this converts the original pairwise block-coordinate output into an ordered chromosome-path representation that can be queried at multiple structural scales.

***Two-stage analytical design.*** We analyzed the multispecies block table in two stages. The first stage asked whether the shared chromosomal rearrangement signal among *D. regia*, *D. capensis*, and *Dionaea* supports one shared, higher-order hexaploid ancestry for the two *Drosera* genomes. This step combined a formal shared-adjacency analysis of the focal triplet with a phylogenomic analysis of orthogroup copy-number classes and gene-tree topologies, used here to quantify alternative topological support rather than to infer a unique reticulation mechanism. In this setting, gene-tree discordance (as evidenced by the DensiTree results reported in the main text) was interpreted only as qualitative evidence of mixed histories, compatible with hybridization and/or incomplete lineage sorting (ILS).

The second stage of the analysis added the other carnivorous Caryophyllales in the block table and asked whether the apparent *D. regia*-*D. capensis* similarity remains exceptional once placed into a wider comparative context, and whether one or more inferred structural channels in *D. regia* show preferential affinity to *Dionaea* rather than to other comparator lineages. Because both stages use the same block-coordinate table, the broader comparisons are directly comparable with the focused triplet analysis.

**Stage 1. Focused analysis of shared rearrangements and topology mixtures**

1. **Shared-adjacency test.** The focused structural analysis for stage 1 began by reconstructing ancestral block adjacencies along chromosomes of extant taxa. Ordered block runs were converted into adjacency pairs by recording each transition between adjacent runs carrying different dominant-*Nepenthes* scaffold labels. Adjacencies were treated as undirected so that

a transition between two scaffold labels was counted as the same structural join regardless of orientation along a chromosome. The central question was whether *D. regia* and *D. capensis* share more such inferred ancestral joins than expected by chance.

Two null models were used. Under the global null, dominant-*Nepenthes* scaffold labels were randomly reassigned across all segments in a genome while preserving the number of segments, the number of segments per chromosome, and the overall frequency of each ancestral label. Under the chromosome-constrained null, labels were shuffled only within extant chromosomes, thereby preserving chromosome-level ancestral composition as well as segment counts. The first model tests whether the observed overlap exceeds a genome-wide random arrangement of ancestral compartments; the second asks whether internal ordering of ancestral segments is more similar than expected once chromosome composition is held constant.

This analysis recovered seven shared adjacency types between *D. regia* and *D. capensis*. Although this was the largest raw overlap among the focal structurally informative pairs, it was not enriched beyond either null expectation. Under the global null, the expected overlap was approximately 8.30 and the empirical probability of observing at least seven shared adjacencies was 0.889. Under the chromosome-constrained null, the expected overlap was approximately 6.31 and the empirical probability of observing at least seven shared adjacencies was 0.426. Thus, the observed overlap is real in a descriptive sense, but it does not exceed the amount of shared structure expected once block inventory and chromosome context are taken into account. The adjacency-overlap result and its global-null permutation test are summarized in **Supplementary Note 2 Fig. 1b,c.**

Several of the shared adjacencies were exclusive to the *D. regia-D. capensis* pair within the sampled set. These pair-specific joins identify shared structural features not observed in the other sampled taxa, but do not establish statistically exceptional sharing or provide sufficient evidence for a shared higher-order hexaploidy. Taxa with richer segment inventories are also more likely to share some adjacent run-to-run joins between ancestral block labels that are not seen in taxa with smaller inventories. Once the relevant constraints are imposed in the null models, the total extent of *D. regia-D. capensis* overlap is not exceptional. Selected candidate shared rearrangements that motivated this focused comparison are illustrated in **Supplementary Note 2 Fig. 1a.**

2. **Orthogroup copy number and topology distributions.** To evaluate whether gene-tree signal supports a uniquely shared *Drosera* hexaploidy, orthogroup inference was rerun using the Orthofinder framework, and gene trees were reconstructed from nucleotide alignments, with branch support assessed by SH-aLRT and ultrafast bootstrapping. Support-filtered trees were defined by simultaneous thresholds of at least 70 for both statistics. Confidence intervals on topology frequencies were estimated by nonparametric bootstrap resampling of orthogroups.

   Orthogroup copy-number classes did not reveal a modal pattern uniquely diagnostic of retained shared hexaploid structure. The most frequent copy-number class was the single-copy configuration, but substantial numbers of orthogroups fell into other classes consistent with lineage-specific duplication and fractionation. This result indicates that the copy-number landscape is heterogeneous and does not by itself support a simple one-event model for the focal taxa.

Among support-filtered single-copy gene trees, all three alternative topologies relating *D. regia*, *D. capensis*, and *Dionaea* were well represented. The *Dionaea-D. regia* grouping was modestly most frequent at 41.6%, followed by *D. regia-D. capensis* at 32.8% and *Dionaea-D. capensis* at 25.6%. When the analysis was expanded using a draw-one synthetic single-copy strategy for multicopy orthogroups, the same pattern persisted. In the expanded support-filtered set, *Dionaea-D. regia* accounted for 42.3%, *D. regia-D. capensis* for 30.9%, and *Dionaea-D. capensis* for 26.8%. The striking feature of both datasets is not a single dominant topology, but rather the persistence of all three alternatives at substantial frequencies. These rooted topology frequencies are summarized in **Supplementary Note 2 Fig. 1d.**

In summary, these topology mixtures do not support a simple bifurcating history that uniquely unites *D. regia* and *D. capensis*. Instead they show substantial gene-tree discordance, compatible with hybridization, incomplete lineage sorting, or both, without distinguishing among those causes. The cloudogram shown in the main text is fully consistent with this interpretation and should be considered a qualitative visualization of that discordance rather than as a formal partitioning of its causes.

#### Stage 2. Broader outgroup-informed structural analysis

**Multispecies adjacency comparison.** The second-stage structural analysis generalized the shared-adjacency framework to the broader block-table comparison taxon set. Ordered runs of dominant-*Nepenthes* scaffold labels were reconstructed for each extant taxon represented in the comparison, and pairwise overlap in adjacency inventories was evaluated under the same global and chromosome-constrained null models described above. This extension changes the interpretive basis of the analysis; instead of asking only whether *D. regia* and *D. capensis* share some ancestral joins, it asks whether that overlap is unusually strong relative to the structural opportunity space created by the full taxon comparison set.

In this broader comparative framework, *D. regia* and *D. capensis* again shared seven unique adjacency types and remained each other's closest meaningful structural comparison in broad terms. The overlap again failed to exceed null expectation, and adding the outgroups did not reveal a uniquely shared *D. regia-D. capensis* higher-order architecture, and it did not materially change the underlying interpretation of the *D. regia-D. capensis-Dionaea* genome-structure problem. Rather, the broader sampling clarified that the *D. regia-D. capensis* resemblance is best understood as evidence of close structural relationship and shared ancestral substrate, not as evidence that the two genomes preserve the same higher-order rearrangement logic. This provides the comparative baseline for the analyses below: *D. regia* and *D. capensis* are close structural relatives and share broad syntenic continuity, but the evidence does not support the specific expectation that capensis is simply a doubled derivative of the extant *D. regia*-like three-channel system. This broader outgroup-informed comparison is summarized in **Supplementary Note 2 Fig. 2.**

**D. capensis *collapse and duplication structure*.** A more targeted test asked whether the six-subgenome architecture of *D. capensis* reduces naturally to the same three-channel organization inferred in *D. regia*. To do this, each chromosome was encoded by a feature vector derived from the ordered run table. Features included normalized ancestral-label composition and adjacency content, thereby summarizing both chromosome composition and local transitions among ancestral compartments. *D. capensis* chromosomes were then clustered across a range of *k* values to assess whether a three-group representation is naturally favored in this feature space.

*D. capensis* showed strong internal duplication structure. Duplicate run signatures were abundant, several chromosomes belonged to duplicated signature classes, and permutation tests that preserved the number of runs per chromosome together with the global *D. capensis* label multiset confirmed that this degree of redundancy was far above null expectation. In this broad sense, *D. capensis* clearly contains repeated higher-order structural units. To test whether these repeated units reduce naturally to a small number of recurrent chromosome classes, chromosomes were grouped in feature space using agglomerative clustering with average linkage and cosine distance, matching the implemented analysis. The key result, however, was that these repeated units did not collapse cleanly into a *D. regia*-like three-channel structure: a three-cluster solution was not preferred. Six clusters outperformed three, and still higher values of *k* yielded stronger *silhouette support*, where silhouette support measures how well chromosomes group within their assigned cluster relative to their separation from other clusters. When duplicated *D. capensis* signatures were compared directly with *D. regia* signatures, the best matches were only partial. *D. capensis* is therefore internally repeat-structured, but not in a way that straightforwardly reproduces the extant *D. regia* architecture. The *D. capensis* duplication-without-clean-collapse result is summarized in **Supplementary Note 2 Fig. 3.**

***Ordered path, motif, graph, and reuse analyses.*** Because simple adjacency counts may miss higher-order architectural structure, additional analyses treated each chromosome as an ordered path of ancestral labels after collapsing identical consecutive runs. Path similarity was quantified using normalized longest-common-subsequence style measures summarized across chromosomes. Shorter local path motifs were also counted, and adjacency structure was represented as weighted graphs in which nodes corresponded to ancestral scaffold labels and edges corresponded to observed adjacencies. These complementary representations test whether a shared ancient structural scaffold might persist at the level of whole paths, local fragments, graph topology, or edge reuse even if raw shared-adjacency counts are not striking.

None of these analyses supported a uniquely shared *D. regia-D. capensis* higher-order architecture. Ordered path similarity was only modestly above the null mean and not significant. Shared three-label motifs were not enriched beyond null expectation. Weighted adjacency graph similarities also fell near or below null expectation. Breakpoint reuse spectra, measured as the similarity of edge-frequency profiles across genomes, did not show unusually high correspondence between *D. regia* and *D. capensis*. Likewise, correlations in the contributions of individual ancestral scaffold labels to the adjacency landscape were mild at most and not exceptional. The different structural representations gave the same qualitative result: *D. regia* and *D. capensis* share substantial substrate, but they do not share an unusually conserved higher-order architectural logic.

***Resampling, localization, and optimized capensis pairing.*** To test whether the *D. regia-D. capensis* signal might be driven disproportionately by a small number of chromosomes or by localized structural hotspots, chromosome-level jackknife and bootstrap resampling analyses were performed, together with localization statistics that compared the chromosomal distribution of shared edges with null expectations. These analyses showed that the *D. regia-D. capensis* signal is reasonably stable but quantitatively limited. Shared-edge counts and path-similarity estimates varied only within narrow ranges under resampling, indicating that the overall result is not an artifact of one or two unusual chromosomes. At the same time, the signal remained limited in magnitude: it did not intensify under resampling into the level of excess overlap, path concordance, or chromosomal concentration expected for a strong shared-architecture result. Shared edges were not concentrated in a single exceptional core, but instead were distributed more diffusely across the genome. These resampling, localization, and optimized-pairing analyses are summarized in **Supplementary Note 2 Fig. 4.**

We next searched explicitly for the best partition of *D. capensis* chromosomes into three paired units, as predicted by a doubled-three-channel model. Within *D. capensis*, this optimization recovered very strong pair structure, confirming that the genome contains internally duplicated structural units. However, when those optimized *D. capensis* chromosome pairs were matched against *D. regia* chromosome signatures, the resulting alignment score was not significantly greater than expected under the random-pair benchmark. The distinction is straightforward: *D. capensis* does show strong internal duplication structure; it does not, however, behave simply like a doubled version of the extant *D. regia*-like three-channel genome architecture.

***Support-weighted edge overlap.*** Support weighting produced the only mildly positive *D. regia-D. capensis* result among the structural tests. Here, similarity was calculated from shared inferred block joins, each defined by neighboring labeled runs on a chromosome. Similarity was quantified using the Jaccard index[60], which measures the fraction of shared joins relative to the total set of joins observed across the two genomes. In the unweighted version, each occurrence of a join contributes equally, so repeated occurrences of the same adjacency type are retained in the comparison. Weighted versions instead scale adjacency contributions by the supporting block evidence or ordinal span.

Two weighting schemes were examined: in the first, joins were weighted by block support, meaning that joins built from runs supported by more underlying syntenic blocks contributed more strongly than joins supported by fewer blocks, in the second, joins were weighted by ordinal span, meaning that joins derived from runs extending across larger ordered portions of the chromosome contributed more strongly than joins derived from shorter segments. When block-support weighting was used, the weighted Jaccard similarity between *D. regia* and *D. capensis* exceeded the mean of a permutation-based null distribution constructed specifically for this pair. The observed value was 0.3018, compared with a null mean of 0.2027, with a nominal empirical probability of $P = 0.0346$ based on 5,000 permutations. This indicates modest support-weighted adjacency enrichment under the specified null model, although the probability was not adjusted for the multiple structural similarity measures examined and does not independently establish a shared higher-order polyploidization event.

Other summary measures did not show the same pattern: the unweighted Jaccard index, in which all joins are counted equally regardless of support, was not unusually elevated; nor were path similarity, which compares the ordered sequences of labeled runs along chromosomes, or edge-reuse correlation, which compares how similarly the two genomes reuse particular joins across their architectures. We therefore interpret the results as evidence that *D. regia* and *D. capensis* share a subset of relatively robust and well-supported structural joins, but do not preserve the same overall higher-order breakpoint architecture. They share some durable structural substrate without retaining the same whole-genome rearrangement logic. An ordered anchor-track representation in the shared dominant-*Nepenthes* label space illustrates the same combination of broad shared substrate and divergent higher-order organization **(Supplementary Note 2 Fig. 5).**

***Channel-specific structural affinity between* D. regia *and* Dionaea.** The negative results for a simple shared *D. regia-D. capensis* higher-order architecture motivated a more focused question: rather than asking whether *D. regia* matches another genome as a whole, we asked whether one or more inferred *D. regia* channels may show preferential affinity to *Dionaea*. We used the same run tables to construct chromosome-level feature vectors. For each chromosome, the vector included total run count; total hit support - meaning the summed support values of all contributing syntenic block mappings on that chromosome - and total ordinal span, meaning the summed extent of the chromosome path measured in ordered block-table units rather than physical base pairs. The vector also included normalized ancestral-label composition by span and by support. These quantities describe how much of the chromosome is assigned to each ancestral scaffold label, scaled either by the relative span occupied by that label or by the relative support associated with it, so that chromosomes can be compared on the same proportional scale. Finally, the vector included normalized adjacency-edge frequencies between consecutive distinct labels, that is, the relative frequencies of joins between neighboring runs carrying different ancestral labels along the chromosome. Together, these quantities combined chromosome composition, local transitions among ancestral compartments, and the overall ordered path of labels into a common structural feature space for comparison across chromosomes.

*D. regia* chromosomes were grouped into three structural channels by hierarchical agglomerative clustering after feature standardization, using Ward linkage[61], which merges chromosomes so as to minimize the increase in within-group variance at each step and is therefore well suited to identifying compact structural groups in a continuous multivariate space. Comparator taxa were clustered at fixed values of $k$ chosen to match the design of the comparison. *Dionaea* was treated differently because its block-table representation supports a natural two-channel interpretation, but the specific assignment of chromosomes to those two channels is not uniquely determined in advance. All admissible two-channel assignments consistent with the observed block structure were therefore enumerated. Equivalent assignments that differed only by swapping the labels of the two inferred *Dionaea* channels were counted only once, which was achieved by fixing the orientation of one ancestral label and thereby removing label-exchange symmetry that carries no biological meaning. For each assignment, the resulting channel profiles and ordered path signatures were compared with the three *D. regia* channels. Affinity was scored by a composite of ancestral-label cosine similarity and ordered-path similarity. Cosine similarity was used because it measures the similarity in relative composition of multicomponent profiles while being less sensitive to absolute magnitude[62], making it appropriate for comparing chromosomes or channels that may differ in total support or total span but still share similar proportional ancestry-label structure.

We asked whether the best structural match between a given *D. regia* channel and *Dionaea* was stronger than the best match that same *D. regia* channel could achieve with channels from any non-*Dionaea* comparator taxon. The difference between these two scores was termed the *exclusivity margin*. The exclusivity margin therefore measures how much better *Dionaea* explains a *D. regia* channel than the best alternative available elsewhere in the sampled comparison set. A positive margin means that *Dionaea* is the single best match for that *D. regia* channel within the analyzed structural space, whereas a negative margin means that some non-*Dionaea* comparator provides an equal or better explanation of the observed resemblance.

We benchmarked this result against every two-channel combination drawn from the non-*Dionaea* taxa. Every channel pair from the non-*Dionaea* comparison set was treated as a hypothetical two-channel *Dionaea*-like comparator and analyzed using the same matching procedure. For each pseudo-comparator, the best match to each *D. regia* channel was identified and its exclusivity margin was calculated against the remaining comparison pool. The resulting distribution shows how large an apparent *Dionaea*-like advantage could arise purely from arbitrary two-channel subsets elsewhere in the dataset. The observed *D. regia*-*Dionaea* exclusivity could then be compared against this empirical background to test whether it was genuinely unusual.

The strongest interpretation was not a broad two-channel sharing pattern between *Dionaea* and *D. regia*. Instead, under the selected overall two-channel assignment for *Dionaea*, two *D. regia* channels showed positive exclusivity, meaning that *Dionaea* provided their best match within the sampled comparison set. However, this pattern was weak overall: the number of positively matching *D. regia* channels did not exceed the empirical benchmark at the conventional 0.05 threshold (P = 0.0667), and, in a separate maximum-over-assignments benchmark, the largest summed positive margin across all admissible assignments (0.084710) was not greater than expected from arbitrary two-channel comparator sets drawn from the non-*Dionaea* taxa (P = 0.244).

By contrast, one *D. regia* channel showed a distinctly stronger signal than the others. *D. regia* channel R2 had the clearest and most benchmark-supported *Dionaea*-like affinity, with a best exclusivity margin of 0.084710 and an empirical benchmark probability of P = 0.022. This indicates that the resemblance between R2 and one inferred *Dionaea* channel is stronger than would usually be produced by arbitrary two-channel subsets elsewhere in the dataset. The other *D. regia* channels did not show comparably strong evidence. R1 appeared weakly *Dionaea*-favoring under some assignments, but only with a very small advantage, whereas R3 showed no *Dionaea*-specific signal. The corresponding channel-affinity comparisons are shown in **Supplementary Note 2 Fig. 6.**

These results do not support the view that *D. regia* broadly shares two channels, or a complete higher-order polyploid architecture, with *Dionaea*. Instead, one major *D. regia* structural channel, used here as a proxy for one subgenomic component, shows preferential *Dionaea*-like affinity. This is more consistent with partial subgenome sharing, or with one *Dionaea*-like contributor to the *D. regia* genome history, than with a model in which *D. regia* as a whole derives from the same overall polyploid ancestry as *Dionaea*.

***Interpretation*****.** Overall, the structural and topological analyses point in the same direction: *D. regia* and *D. capensis* are clearly related and share broad syntenic continuity. *D. capensis* also contains strong internal duplication structure, consistent with further genome multiplication or repeated higher-order structural units, although these units do not reduce cleanly to the extant three-channel organization inferred in *D. regia.* However, the strongest tests of a uniquely shared higher-order architecture are negative: *D. regia*-*D. capensis* adjacency overlap is not enriched beyond null expectation, whole-path similarity is not unusually high, and motif and graph analyses do not reveal a concealed shared higher-order backbone. Likewise, the two genomes do not show unusually similar patterns in how often particular block joins are reused across their chromosome structures, and even the *D. capensis* chromosome pairings selected to test the doubled-three-channel hypothesis did not show unusually strong correspondence to individual *D. regia* chromosome signatures relative to randomly selected chromosome pairings. The only mildly positive result was an excess of shared joins when those joins were weighted by the strength of the underlying block evidence. This pattern is consistent with some shared robust structural substrate, but it does not extend to a genome-wide signal of the same higher-order rearrangement architecture.

The gene-tree analyses are consistent with this picture: alternative topologies relating *D. regia*, *D. capensis*, and *Dionaea* are all well represented, and the cloudogram in the main text also shows the corresponding mixture qualitatively. These data show substantial gene-tree discordance and argue against a single simple species-tree history for these lineages. Again, this discordance is taken as qualitative evidence compatible with hybridization, incomplete lineage sorting, or both, but not as a formal distinction among those processes.

Within that broader context, the channel-specific *D. regia*-*Dionaea* result is particularly informative. One inferred *D. regia* channel, especially R2, shows a preferential and benchmark-supported affinity to *Dionaea* that is not mirrored evenly across the rest of the *D. regia* system. The analyses suggest that *D. regia* and *D. capensis* descend from related ancestral genomic substrate but experienced largely independent higher-order restructuring histories, while *D. regia* additionally preserves a channel-specific structural signal of affinity to a *Dionaea*-like lineage. This pattern is more consistent with a mosaic or reticulate higher-order genome history than with a single shared higher-order *Drosera* hexaploidy followed by simple *D. capensis* doubling, although these analyses alone do not distinguish direct reticulation from differential retention of deeper shared genomic components.

### Supplementary Note 2: Figures

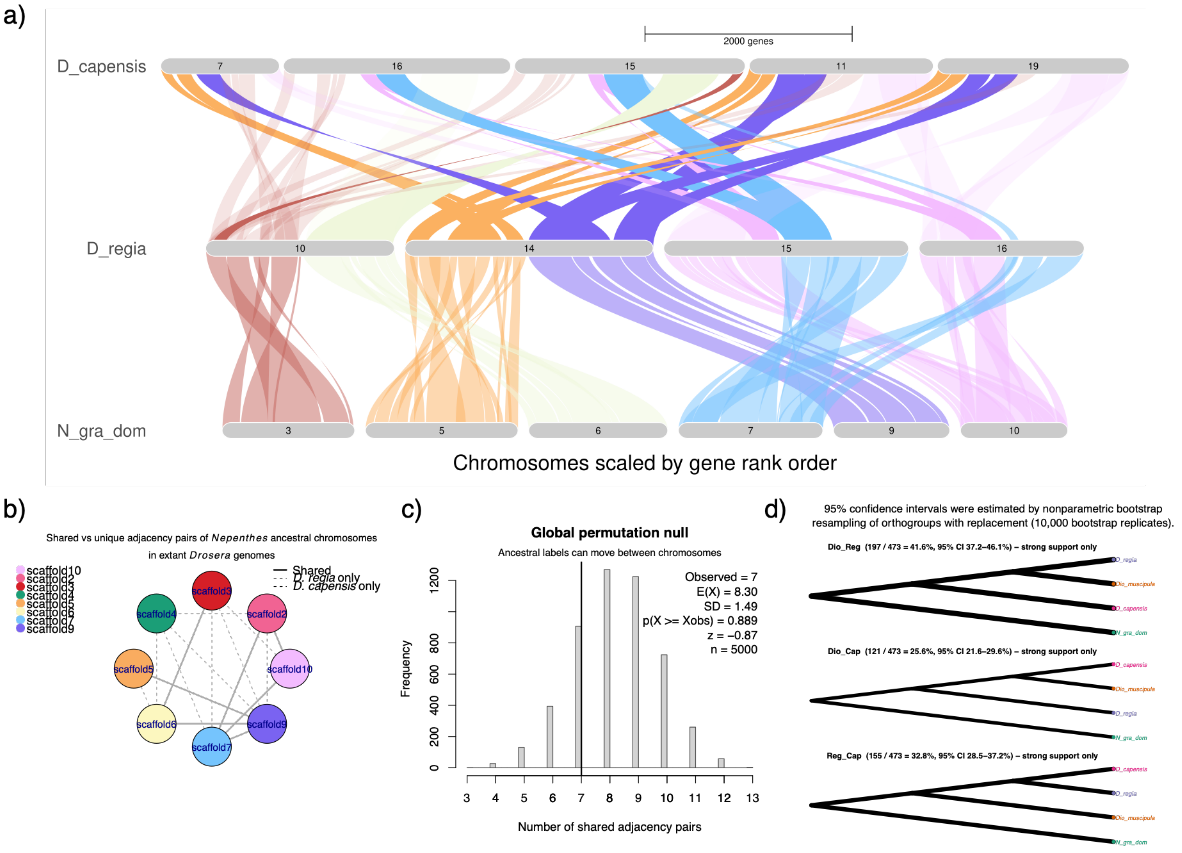


**Supplementary Note 2 Fig. 1. Focused evidence for and against a shared hexaploidy in *D. capensis* and *D. regia*.** **(a)** GENESPACE riparian plot illustrating candidate shared chromosomal rearrangements among selected chromosomes from the *Nepenthes gracilis* dominant subgenome (N. gra_dom)*, D. regia*, and *D. capensis*. This subset highlights examples that initially appeared consistent with shared ancestral restructuring, including the orange:purple fusion in *D. regia* chromosome 14 and apparently corresponding regions in *D. cap*ensis chromosomes 7, 11, and 19, together with the pink:sky-blue fusion shared by *D. regia* chromosomes 15 and 16 and reflected in *D. capensis* chromosomes 15 and 16. **(b)** Network plot of inferred ancestral chromosome adjacencies in *D. regia* and *D. capensis*; solid edges indicate shared adjacency pairs and dashed edges indicate species-specific pairs. **(c)** Distribution of the number of shared adjacency pairs under a global permutation null model in which ancestral chromosome labels were randomly reassigned across segments while preserving global label frequencies. The vertical line marks the observed overlap. **(d)** Frequencies of the three resolved rooted gene-tree topologies among single-copy orthogroups for N. gra_dom, *Dionaea*, *D. regia*, and *D. capensis*, with support-filtered trees defined by SH-aLRT ≥ 70 and UFBoot ≥ 70. Trees were rooted with N. gra_dom, and only gene trees with strong support for the focal sister pairing were retained. Line thickness is proportional to the number of orthogroups supporting each topology, with labels indicating counts, proportions, and 95% confidence intervals from bootstrap resampling.

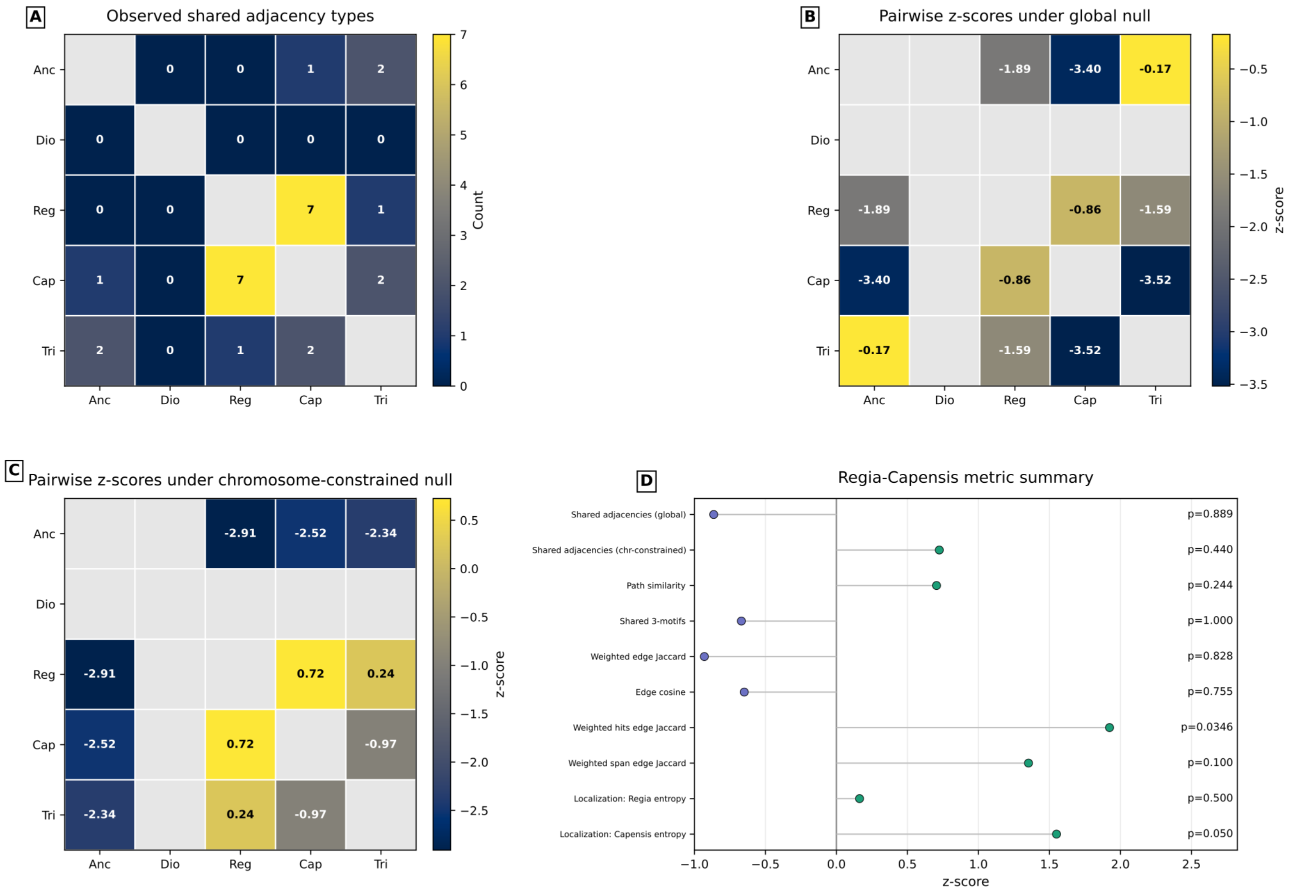


**Supplementary Note 2 Fig. 2. Broader outgroup-informed structural comparison. (a)** Heat map showing the observed number of shared adjacency types for pairwise comparisons among *Dionaea* (Dio), *D. regia* (Reg), *D. capensis* (Cap) and the broader comparison taxa, *Triphyophyllum* (Tri) and *Ancistrocladus* (Anc), all represented in the shared dominant-*Nepenthes* scaffold-label space (N_gra_dom). Here, an adjacency type refers to a connection between neighboring syntenic block identities in the inferred structural representation. Cell values and the color scale indicate the number of shared adjacency types. In this panel, strongly yellow cells indicate pairs with high raw observed overlap in adjacency types. **(b)** Heat map showing pairwise z-scores under a global permutation null model for the same taxon pairs. In this null model, ancestral or chromosomal labels were randomized across the genome while preserving their overall frequencies. Cell values are z-scores, with more positive values indicating greater structural similarity than expected by chance. Axes list the taxa being compared; the color bar shows z-score. In this panel, strongly yellow cells indicate pairwise overlap that is less depleted, or more enriched, relative to expectation under genome-wide randomization that preserves overall label frequencies; such cells therefore reflect deviation from the global null rather than raw overlap alone. **(c)** Heat map showing pairwise z-scores under a chromosome-constrained null model, in which randomization was restricted within chromosomes rather than across the whole genome. Values are again shown as z-scores. This tests whether apparent similarity remains unusual after preserving chromosome-level composition. In this panel, strongly yellow cells indicate pairwise overlap that is relatively higher than expected after chromosome-level label composition has been preserved, and therefore point more specifically to similarity in within-chromosome adjacency structure rather than simply shared overall label usage. **(d)** Summary plot of the *D. regia-D. capensis* comparison across multiple structural metrics. Each point gives the z-score for one metric, with the corresponding permutation P value shown at right. Metrics include shared adjacencies under the global and chromosome-constrained null models, path similarity, shared three-step motifs, weighted edge Jaccard similarity, edge cosine similarity, weighted-hit edge Jaccard similarity, weighted-span edge Jaccard similarity and localization statistics for *D. regia* and *D. capensis*. The x-axis shows z-score and the y-axis lists the metric. Across these broader outgroup-informed comparisons, *D. regia* and *D. capensis* remain structurally related, but the analyses do not reveal hidden support for a uniquely shared higher-order *D. regia-D. capensis* architecture. The most conservative interpretation is therefore broad syntenic continuity on a shared ancestral substrate, rather than preservation of a uniquely shared rearrangement system.

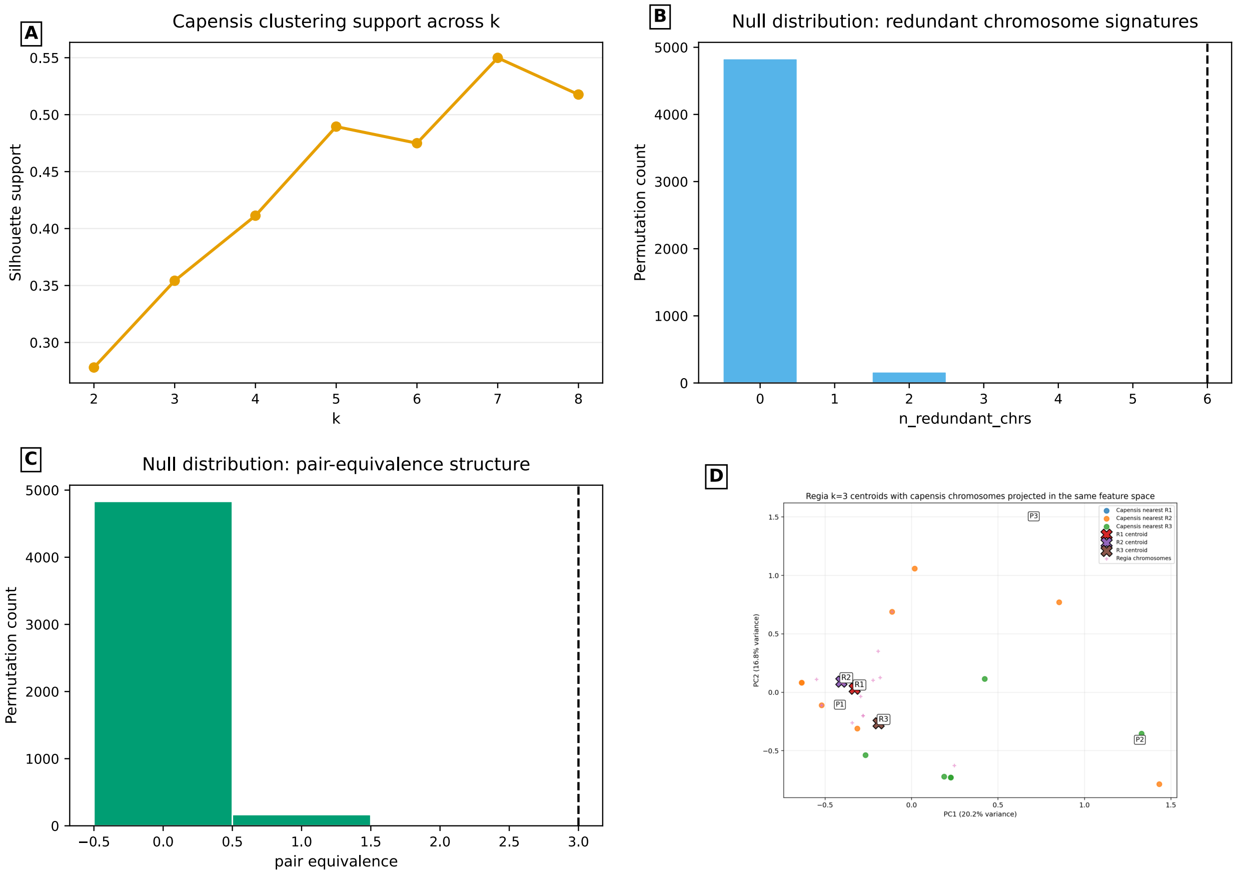


**Supplementary Note 2 Fig. 3. *D. capensis* duplication without clean collapse to a *D. regia*-like three-channel organization. (a)** Line plot showing clustering support across increasing values of *k* for *D. capensis* chromosomes. The x axis shows the number of clusters tested (*k*) and the y-axis shows silhouette support, a unitless measure of how well chromosomes group together. with the highest silhouette support at seven clusters in the tested range, indicating substantial internal structure within *D. capensis*, but not by itself identifying a *D. regia*-like three-group solution. **(b)** Histogram showing the null distribution for the number of redundant D. capensis chromosome signatures under 5,000 randomizations. The x-axis gives the number of redundant chromosomes, and the y-axis gives the permutation count. The dashed vertical line marks the observed value of six. **(c)** Histogram showing the null distribution for the pair-equivalence statistic. The x-axis shows the pair-equivalence statistic (unitless), which summarizes how strongly chromosome pairs in *D. capensis* behave as duplicated structural counterparts, and the y axis shows the number of permutations. The dashed vertical line marks the observed value. **(d)** Projection of *capensis* chromosomes into the same feature space as the *D. regia* *k* = 3 centroids. The x- and y -axes show principal component 1 (PC1) and principal component 2 (PC2), respectively, with the percentage of variance explained indicated on the axes. Point colors indicate, for each *capensis* chromosome, the nearest *D. regia* centroid in feature space; pair labels P1, P2 and P3 identify the three strongest internal *D. capensis* chromosome pairs, while X markers indicate the three reference centroids (R1, R2 and R3). These pair labels mark inferred *capensis* duplicated chromosome pairs. Their positions relative to R1, R2 and R3 show which *D. regia* centroid they most closely approach in the reduced structural feature space; however, the *D. capensis* pair/groups are dispersed across the plot, with different pairs/groups lying nearest different *D. regia* centroids rather than collapsing into three compact *capensis* clusters centered on the three *D. regia* centroids. This panel asks whether the repeated structure within *capensis* collapses naturally into the same three-group pattern observed in extant *D. regia*, which it does not. Together, these analyses show that *D. capensis* contains strong internal duplication structure, but that structure does not reduce cleanly to a *D. regia*-like three-channel organization. Thus, *D. capensis* is not simply *D. regia* doubled.

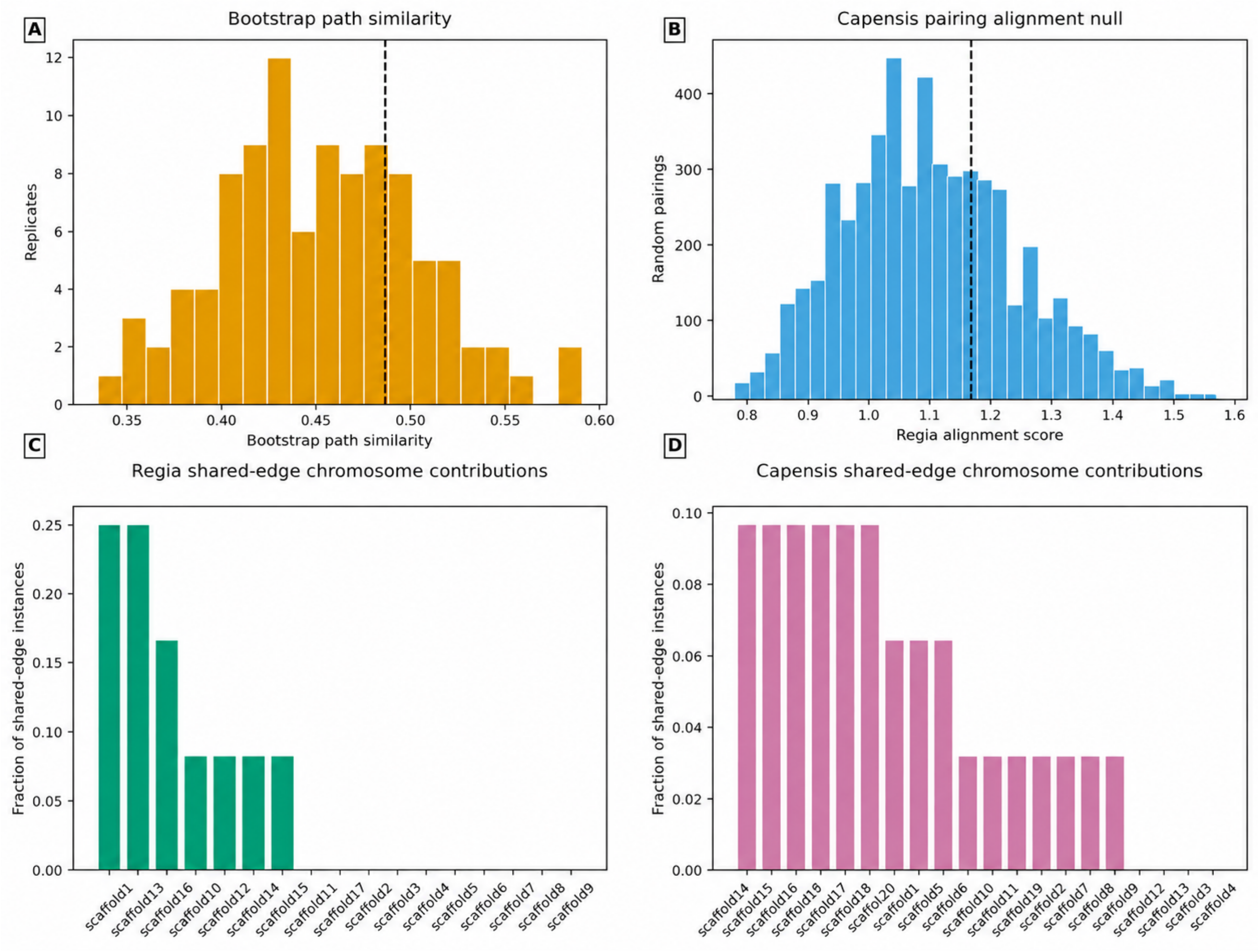


**Supplementary Note 2 Fig. 4. Resampling, optimized-pairing, and chromosome-contribution analyses. (a)** Histogram showing bootstrap path-similarity values across resampled replicates for the *D. regia-D. capensis* comparison. The x axis shows bootstrap path similarity (unitless) and the y axis shows the number of replicates. The dashed vertical line marks the observed value. This tests whether the path-based similarity between *D. regia* and *D. capensis* is robust to resampling. **(b)** Histogram showing the null distribution for the *capensis* pairing-alignment score. The x-axis shows the *D. regia* alignment score (unitless) and the y-axis shows the number of random pairings. The dashed vertical line marks the observed score for the optimized *D. capensis* pairing. This tests whether the duplicated chromosome structure within *capensis* aligns to *D. regia* more strongly than expected by chance. **(c)** Bar plot showing *D. regia* chromosome contributions to shared-edge signal. The x-axis lists *D. regia* chromosomes or scaffolds and the y-axis shows the fraction of shared-edge instances attributable to each chromosome. This assesses whether the shared structural signal is dominated by a single *D. regia* chromosome or distributed more broadly. **(d)** Bar plot showing *D. capensis* chromosome contributions to shared-edge signal. The x-axis lists *capensis* chromosomes or scaffolds and the y axis shows the fraction of shared-edge instances attributable to each chromosome. As in **(c)**, this tests whether the signal is concentrated in one chromosome or spread more broadly across the genome. Together, these analyses show that the modest *D. regia-D. capensis* structural signal is distributed across multiple chromosomes rather than driven by a single outlier chromosome. They also reveal strong internal duplicate structure within *D. capensis*, while showing that the optimized *D. capensis* chromosome pairing does not align unusually well with individual *D. regia* chromosome signatures.

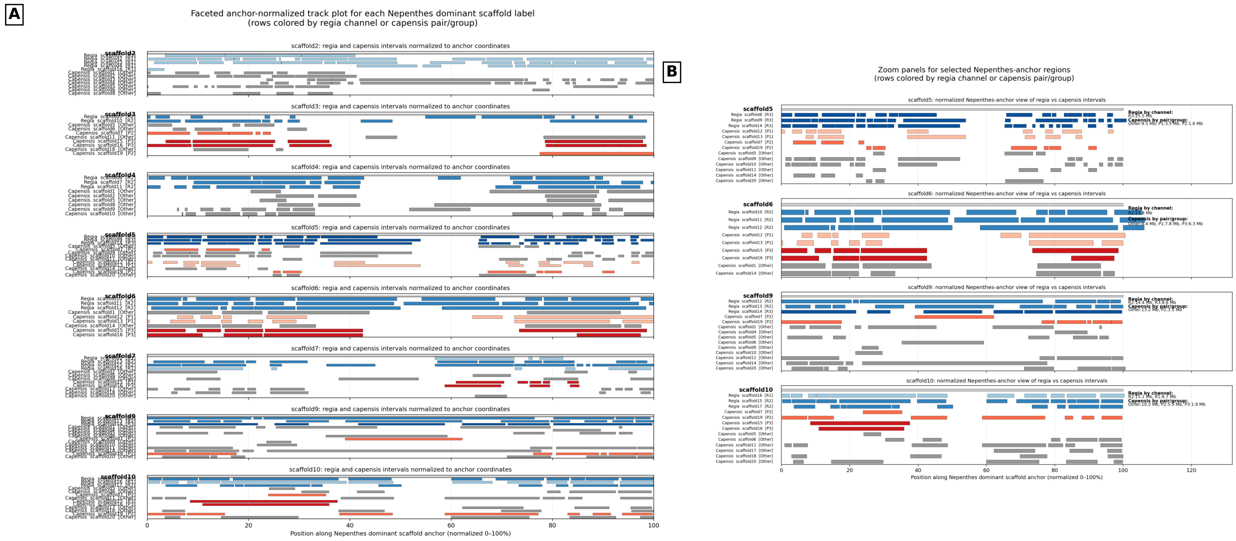


**Supplementary Note 2 Fig. 5. Ordered anchor-track comparison of *D. regia* and *D. capensis* in shared dominant-*Nepenthes* label space. (a)** Faceted anchor-normalized track plots for each dominant *Nepenthes* scaffold label, showing corresponding intervals from *D. regia* and *D. capensis* in a shared coordinate system. Each facet represents one dominant *Nepenthes* scaffold label. The x-axis shows position along the anchor track normalized from 0 to 100% of scaffold span, rather than physical chromosome coordinates in base pairs. Rows are colored by inferred *D. regia* channel or *capensis* pair or group. This panel provides a genome-wide visual summary of how *D. regia* and *capensis* intervals occupy the same inferred ancestral-label space. **(b)** Zoomed views of selected dominant-*Nepenthes* scaffold regions from **(a)**. The x-axis again shows normalized position along the anchor track, and rows show the corresponding *D. regia* and *D. capensis* intervals colored by channel or pair or group assignment. These zoomed views highlight local similarities in ancestry-label usage and order, while also showing that *D. capensis* does not resolve visually as a simple doubled version of the extant *D. regia*-like three-channel pattern. Together, these anchor-track plots provide an intuitive summary of shared substrate together with different higher-order organization. The coordinates represent ordered anchor-block position in a shared dominant-label space, not physical base-pair position, so the visual similarity reflects shared use and ordering of ancestral labels rather than literal chromosome-scale collinearity in base pairs.

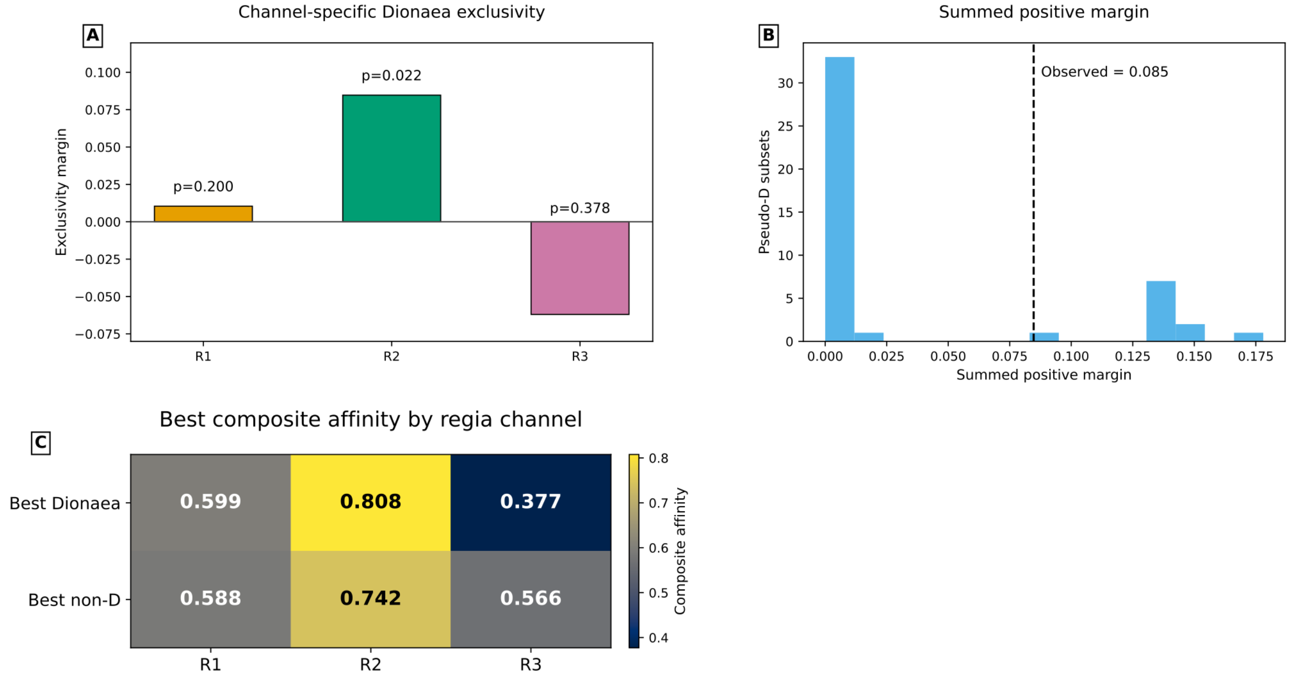


**Supplementary Note 2 Fig. 6. Channel-specific structural affinity between *D. regia* and *Dionaea*.** (a) Bar plot showing the *Dionaea* exclusivity margin for each inferred *D. regia* channel (R1, R2 and R3). The x-axis lists *D. regia* channels and the y-axis shows the exclusivity margin (unitless), where more positive values indicate stronger preferential similarity to *Dionaea* than to non-*Dionaea* comparison channels. Empirical P values are shown above the bars. Among the three channels, R2 shows the clearest positive *Dionaea*-specific signal. **(b)** Histogram showing the null distribution of the summed positive margin across pseudo-*Dionaea* subsets. The x-axis shows the summed positive margin (unitless) and the y-axis shows the number of pseudo-*Dionaea* subsets. The dashed vertical line marks the maximum summed positive margin obtained across all admissible assignments (0.084710). This maximum was not greater than expected under the pseudo-*Dionaea* benchmark (P = 0.244). **(c)** Heat map showing the best composite affinity score for each *D. regia* channel. Columns correspond to *D. regia* channels (R1-R3), and rows compare the best *Dionaea* composite with the best non-*Dionaea* composite. Cell values are composite affinity scores (unitless), shown numerically and by color scale. These comparisons show that the strongest positive signal is channel specific rather than genome wide: *D. regia* channel R2 shows the clearest benchmark-supported *Dionaea* affinity, whereas broader two-channel sharing is weaker and more sensitive to assignment. The figure therefore supports a restrained interpretation of partial channel-level structural affinity, rather than whole-genome equivalence or proof of a shared polyploid event.

**Methods for Supplementary Note 2**

***Input block table and run representation.*** All structural analyses were based on a multispecies syntenic-block coordinate table generated from GENESPACE, in which extant chromosomes were projected into the dominant-*Nepenthes* scaffold-label namespace used as the shared ancestral reference space. For each focal taxon, rows were sorted by chromosome and ordinal-coordinate midpoint; where ordinal coordinates were unavailable in earlier focused visualizations, base-pair midpoint order was used instead. Consecutive rows carrying the same dominant ancestral scaffold label were collapsed into runs, each representing a contiguous chromosome segment assigned to a single ancestral scaffold label.

***Focused shared-rearrangement screening.*** For **Supplementary Note 2 Fig. 1a**, candidate shared rearrangements were identified by first locating end-to-end fusions of non-homoeologous ancestral compartments in *D. regia* and then examining the corresponding *D. capensis* regions containing the same ancestral *Nepenthes*-like compartments. Candidates were retained when the *D. regia* breakpoint boundaries corresponded to syntenic-block boundaries representing the same inferred fusion in *D. capensis*. These examples motivated the subsequent formal tests and were not treated as independent evidence for a shared higher-order history.

***Adjacency inference and permutation testing.*** Adjacencies were defined from the run tables as undirected pairs of neighboring runs assigned to different ancestral scaffold labels. Unique adjacency pairs were extracted for each species and visualized as a network in which nodes represented ancestral scaffold labels and edges represented inferred adjacencies.

Observed overlap in adjacency sets was tested by Monte Carlo permutation. In the global null model, ancestral scaffold labels were shuffled across all segments without replacement while preserving the overall label multiset. In the chromosome-constrained null model, labels were shuffled only within chromosomes, preserving chromosome-level composition while randomizing internal order. In the implemented focused and multispecies adjacency analyses, 5,000 permutations were performed with a fixed random seed. For each replicate, adjacency sets were recalculated and the number of shared unique adjacency pairs was recorded. In the focused analysis, empirical one-sided probabilities used an add-one Monte Carlo correction; in the multispecies analysis, they were estimated as the fraction of null replicates with values greater than or equal to the observed value. Z-scores were calculated from the empirical null mean and standard deviation.

***Gene-tree topology summary for the focused four-taxon comparison.*** For the topology component of **Supplementary Note 2 Fig. 1**, orthogroups were inferred with OrthoFinder v3.1.2, CDS sequences were aligned with MAFFT, and gene trees were reconstructed using IQ-TREE v3.0.1 with the dominant *Nepenthes* reference channel as outgroup. For each orthogroup, the ingroup topology was assigned according to which of the three possible species pairs formed a monophyletic clade. Support for the defining node was extracted from SH-aLRT and UFBoot values. Trees were summarized both before and after applying a joint node-support filter requiring SH-aLRT $\geq 70$ and UFBoot $\geq 70$. The strict summary shown in **Supplementary Note 2 Fig. 1d** used orthogroups with exactly one tip per species. For the expanded analysis reported in the text, multi-copy trees containing all four species were pruned to one randomly sampled tip per species with fixed seed 1 and pooled with the strict single-copy set. Uncertainty in topology proportions was estimated by nonparametric bootstrap resampling of orthogroups with replacement using 10,000 bootstrap replicates. To assess sensitivity to random paralog selection in the expanded dataset, we repeated the one-tip-per-species sampling across 100 independent random seeds, retaining the strict single-copy trees in every replicate. This sensitivity analysis used the full expanded set of classified gene trees, without applying the SH-aLRT and UFBoot strong-support thresholds. The *Dionaea-D. regia* topology remained the most frequent in all 100 replicates, with proportions ranging from 0.3989 to 0.4271 (mean 0.4102). The corresponding ranges were 0.2933-0.3118 for *D. regia-D. capensis* and 0.2787-0.2965 for *Dionaea-D. capensis*. The latter two topologies exchanged rank in only one replicate. These results indicate that the predominant *Dionaea-D. regia* signal is robust to alternative paralog selections, although the relative frequencies of the two less common topologies are more similar.

***Chromosome feature vectors and clustering.*** For higher-order structural analyses, each chromosome was represented by a feature vector derived from the run table. These vectors included chromosome-wide ancestral-label composition and adjacency content; in weighted analyses they additionally incorporated total hit support and total ordinal span. Internal structure within *D. capensis* was evaluated by agglomerative clustering. In the *D. capensis*-collapse analysis, silhouette support was evaluated across $k = 2$ to 8 (or to $n - 1$ if fewer chromosomes were available), using average linkage with cosine distance. A fixed $k = 6$ ordering was then used for the capensis feature heat map, whereas *D. regia* was represented by a fixed $k = 3$ clustering for the extant three-channel comparison.

***Duplicate-signature permutation test in D. capensis.*** To test whether *D. capensis* contained more repeated chromosome-scale structure than expected by chance, per-chromosome run signatures were compared exactly. The observed statistics were the number of redundant chromosomes, the number of duplicated signatures, the maximum signature multiplicity, and the pair-equivalence count. Null signatures were generated by reshuffling the global ancestral-label multiset without replacement while preserving the observed number of runs per chromosome. The implemented analysis used 5,000 permutations with a fixed random number seed. This test asked whether repeated chromosome-scale signatures in *D. capensis* exceeded random expectation under fixed chromosome complexity.

***Path, motif, and graph-based comparisons.*** To test for hidden similarity beyond simple adjacency overlap, chromosomes were also represented as ordered label paths after collapsing repeated consecutive runs. Pairwise comparisons then used several complementary metrics: symmetric mean best normalized longest common subsequence (nLCS) for whole-path similarity; counts of shared ordered 3-motifs; weighted Jaccard similarity of adjacency-edge reuse; and cosine similarity of edge-count vectors. Here, normalized longest common subsequence measures how much of the ordered label sequence in one chromosome can be matched in the same order to another chromosome, without requiring uninterrupted contiguity. The score was normalized for path length so that chromosomes of different sizes could be compared on the same scale, and the "symmetric mean best" form means that the best match from genome A to genome B and from genome B to genome A were both calculated and then averaged, reducing directional bias. High nLCS values therefore indicate that two chromosomes preserve a similar overall order of ancestral labels across much of their length, even if local interruptions, insertions, or rearrangements are present. In the implemented path/motif/graph phase, generated by permuting the pooled label tokens while preserving the observed path lengths of each chromosome. These analyses used 1,000 permutations with a fixed random number seed.

***Channel-affinity analysis.*** For the channel-affinity analysis, *D. regia* chromosomes were clustered into three inferred channels. Comparator taxa were grouped from the same feature-space framework, using two channels for *Ancistrocladus*, six channels for *D. capensis*, and two channels for *Triphyophyllum*. For *Dionaea*, all admissible two-channel assignments consistent with the duplicated ancestral-label structure were evaluated, with one ancestral-label pair fixed to remove channel-swap symmetry. Affinity between channels was quantified from normalized ancestral-label composition, ordered-path similarity, and a composite score equal to the mean of label-cosine similarity and path similarity. For each *D. regia* channel, a *Dionaea* exclusivity margin was calculated as the best *Dionaea* composite score minus the best score obtained from non-*Dionaea* comparator channels. Benchmarking used all possible pseudo-*Dionaea* two-channel subsets drawn from the non-*Dionaea* comparator pool.

***Support-weighted and localization analyses.*** In the later refinement phase, edges were quantified in both unweighted and weighted forms. Weighted joins were calculated from the mean of the two flanking runs, using either summed block hits or summed ordinal span as the weight. Pairwise comparisons included occurrence-count-based (multiset) edge Jaccard similarity, weighted-hit edge Jaccard similarity, weighted-span edge Jaccard similarity, edge-based Spearman and Pearson correlations, label-incidence correlations, and whole-path similarity. Null distributions were generated by independently permuting ancestral-block labels genome-wide within each species, pooling labels across chromosomes and redistributing them among the original run positions. This procedure preserved each species' overall ancestral-label frequencies, the number and positions of runs on each chromosome, chromosome boundaries, and the original block-support weights assigned to run positions, but did not preserve chromosome-specific ancestral-label composition. Support-weighted adjacency scores were recalculated after each permutation using the original weighting definitions. The focal *D. regia-D. capensis* block-support-weighted Jaccard comparison was evaluated using 5,000 permutations with random seed 20261010.

To assess whether shared signal was localized or broadly distributed, chromosome-level shared-edge contributions were summarized and compared with permutation-based null expectations using normalized Herfindahl-Hirschman concentration and Shannon entropy statistics[63-65]. The normalized Herfindahl-Hirschman statistic measures concentration of signal across chromosomes, with higher values indicating that shared-edge support is concentrated in a small number of chromosomes and lower values indicating a more even spread. The normalized entropy statistic measures the converse pattern of dispersion, with higher values indicating that shared-edge support is distributed more evenly across many chromosomes and lower values indicating stronger localization. Together, these complementary statistics distinguish whether apparent similarity is driven by one or a few chromosome outliers or is broadly distributed across the genome. Robustness was further assessed by chromosome jackknifing and by bootstrap resampling of chromosomes with replacement. The implemented bootstrap analysis used 100 replicates.

***Direct test of the doubled-three-channel hypothesis.*** To test whether *D. capensis* behaved like a doubled version of the extant *D. regia* three-channel system, *D. capensis* chromosomes with at least three runs were paired by maximum-weight matching on within-*D. capensis* path similarity, with a small bonus for exact signature identity. The top three pairs were then matched to distinct *D. regia* chromosomes with at least two runs using the Hungarian assignment algorithm[66], which maximizes total pair-to-chromosome similarity rather than matching pairs greedily. Similarity was quantified using normalized longest common subsequence scores, yielding the globally optimal one-to-one assignment under this criterion. Significance was evaluated against a null distribution generated from 5,000 random three-pair selections from *D. capensis*, using a fixed random seed and the same optimal assignment procedure for each replicate. The observed alignment score was 1.1667, compared with a null mean of 1.0873 (empirical $P = 0.270$), indicating that the internally optimized *D. capensis* chromosome pairs did not exhibit unusually strong correspondence to *D. regia* chromosome signatures relative to randomly selected chromosome pairings.

### Supplementary Note 3: Fractionation-scar and hidden-Markov reconstruction of polyploid genomic relationships

Main-text **Fig. 7** summarizes a reticulate genome-history model for the sampled carnivorous Caryophyllales by integrating the sequence-based phylogenetic backbone, syntenic depth, chromosome numbers, $K_S$ relationships, structural rearrangements, and the channel-specific analyses developed in **Supplementary Note 2**. **Supplementary Note 3 Fig. 1** lays out that evidence path explicitly. The two alternatives for the tripartite genome of *D. regia* postulated in **Fig. 7** differ in mechanism: model I invokes a direct *Dionaea*-like contribution to a mosaic allopolyploid, whereas model II explains the same channel-specific signal by differential retention of a deeper shared component. In both models, the critical structural observation is restricted to part of the *D. regia* genome rather than representing a genome-wide equivalence with *Dionaea*.

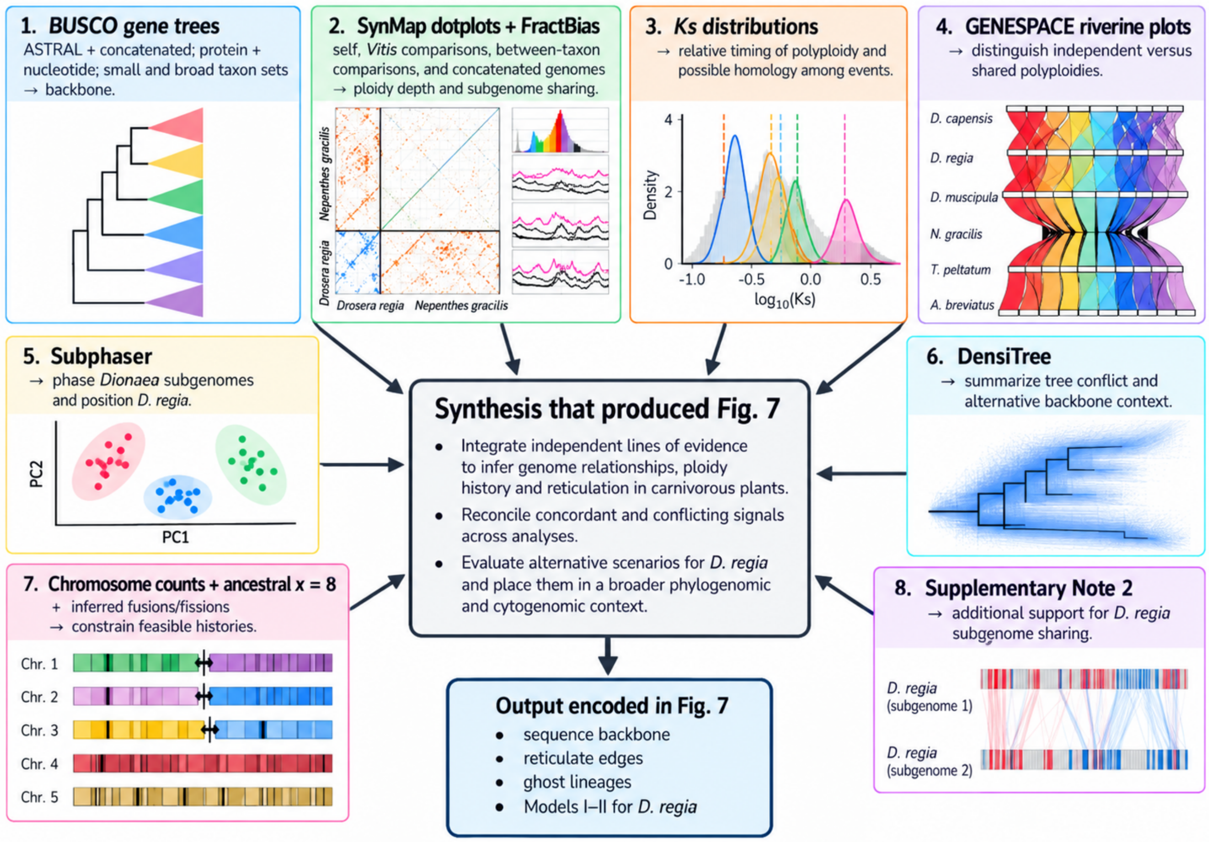


**Supplementary Note 3 Fig. 1**. **Evidence flow behind the genome-history scenarios in main-text Fig. 7**. The figure separates the primary observations from the inferences drawn from them and shows where the sequence-based phylogenetic backbone, synteny/chromosome structure, channel-specific comparisons, Ks results, and chromosome-number information enter the **Fig. 7** synthesis. The fractionation-scar/HMM analysis developed below was not used to construct those scenarios; it provides an independent test of their component structure. The design of this evidence-flow diagram was assisted by ChatGPT (GPT-5.6 Sol); its scientific content and final presentation were reviewed by the authors.

Here, we test the same broader history using post-polyploid patterns of gene retention and loss. The analysis described below remains fully independent of the species tree until the final reconciliation step. Species-tree and gene-tree topologies, $K_S$ distributions, chromosome counts, centromere/repeat evidence, and the event placements in **Fig. 7** do not enter the parental-lineage optimization. Synteny is the only upstream exception because retained duplicates must first be recognized as homologous syntenic copies and placed on a shared ancestral coordinate system. Here, gene-level FractBias output[27] is organized by *Vitis vinifera* chromosomes.

**Analytical logic.** A bifurcating tree captures vertical descent but cannot represent both parental histories of an allopolyploid. We therefore infer relationships among fractionated genomic components as a network without supplying a species tree, and then project the fixed component assignments onto the independently inferred bifurcating backbone. Network representations are well suited to reticulate plant histories[67].

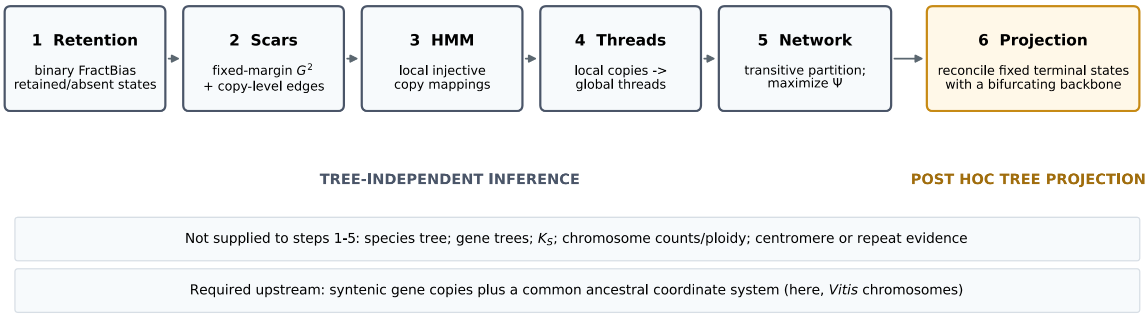


**Supplementary Note 3 Fig. 2. Analytical separation between fractionation-network inference and tree projection.** Steps 1-5 infer genomic-component relationships without a supplied species tree. Step 6 is a constrained projection of the already fixed terminal parental compositions onto the sequence-based phylogenetic backbone. The design of this diagram was assisted by ChatGPT (GPT-5.6 Sol); its scientific content and final presentation were reviewed by the authors.

***Taxa, copy-depth input, and the meaning of operational K.*** The analysis, using a pipeline that we call *FractLineage*, includes seven scored taxa (**Supplementary Note 3 Table 1**). For each species, FractBias/Quota Align provides chromosome-specific retained-copy channels. We call the maximum number available in the input the *FractBias cap*. The operational depth $K$ is the number of these local channels admitted to the scar/HMM pipeline. $K$ is therefore a model-resolution parameter, not a statement of cytological ploidy. For six taxa, the final $K$ equals the available cap. For *Aldrovanda*, an input exposing as many as nine copies allows $K$ = 2-9 to be compared formally, with $K$ = 8 selected by the global criterion described below.

**Supplementary Note 3 Table 1. Taxa and operational copy depths.** The FractBias cap is an input limit; selected ***K*** is the local thread depth used in the final network.

| Taxon | Thread prefix | Family | FractBias cap | selected $K$ |
|---|---|---|---|---|
| *Dionaea muscipula* | D | Droseraceae | 2 | 2 |
| *Drosera regia* | R | Droseraceae | 3 | 3 |
| *Drosera capensis* | C | Droseraceae | 6 | 6 |
| *Nepenthes gracilis* | N | Nepenthaceae | 5 | 5 |
| *Ancistrocladus abbreviatus* | A | Ancistrocladaceae | 2 | 2 |
| *Triphyophyllum peltatum* | T | Dioncophyllaceae | 2 | 2 |
| *Aldrovanda vesiculosa* | L | Droseraceae | 9 | 8 |

***Fractionation scars: shared retention history as a genomic signal.*** After whole-genome multiplication, descendant copies do not retain identical gene sets indefinitely. Genes are lost or retained differentially during diploidization. If a retention/loss decision occurred before two descendant genomic components diverged, both descendants may inherit part of that pattern. We call this inherited correspondence a *fractionation scar*. The informative signal is not simply how many genes two genomes retain, but *which ancestral genes* are retained together in particular copies.

Retained/lost homoeolog patterns have been used previously as historical characters. Reciprocal and differential post-WGD gene loss were established early as lineage-informative genomic characters in yeast[68,69]. POInT subsequently formalized presence/absence evolution across syntenic loci after a shared polyploid event and used those patterns, together with syntenic order, to phase orthologous post-WGD regions across species[70,71]. SCORPiOs provides a close operational analogue: it threads duplicated genomic segments and scores their cross-species syntenic similarity using both orthology information and patterns of gene retention and loss[72]. Treating persistent genomic changes as phylogenetic characters also follows the logic of rare genomic changes[73]. The scar statistic used here differs in purpose and statistical construction: it asks whether exact binary retention patterns are associated between species under a fixed-margin null, without supplying a species phylogeny or assuming that all compared taxa descend from the same polyploid event.

For species $A$ with $m_A$ retained copies over one ancestral chromosome, the state of ancestral gene $g$ is the binary vector

$$\mathbf{x}_g^A = (x_{g1}^A, \ldots, x_{gm_A}^A), \qquad x_{gi}^A \in \{0, 1\}, \tag{1}$$

where 1 denotes retention and 0 denotes absence.

Let $k_g^A = \sum_i x_{gi}^A$ be the number of retained copies. Genes retained in none or all copies of a species are not informative about copy-specific fractionation and are excluded from the exact-pattern statistic.

**Binary example.** Over six ancestral genes, one retained copy might read 1 0 1 1 0 0 and another 1 0 1 0 0 0. Their partly shared pattern contributes to a common fractionation scar; a very different pattern such as 0 1 0 1 1 0 would contribute much less. The method therefore uses the arrangement of 1s and 0s across homologous ancestral genes, not simply overall retention level.

For each species pair, informative genes are stratified jointly by ancestral chromosome $c$ and retained-copy counts $(k_g^A, k_g^B)$. This conditioning prevents high overall retention alone from making two species appear similar. Within a stratum $s = (c, k_A, k_B)$, the observed contingency table $O_{uv}^{(s)}$ counts co-occurring retention patterns $u$ and $v$.

Under independence,

$$E_{uv}^{(s)} = \frac{O_{u+}^{(s)} O_{+v}^{(s)}}{N_s}, \qquad G_{AB}^2 = \sum_s 2 \sum_{u,v:O_{uv}^{(s)}>0} O_{uv}^{(s)} \log\left(\frac{O_{uv}^{(s)}}{E_{uv}^{(s)}}\right). \tag{2}$$

The null distribution is generated by random contingency tables with the observed row and column margins held fixed[74].

With $R = 5{,}000$ randomizations,

$$Z_{AB} = \frac{G_{AB}^{2,\text{obs}} - \mu(G_{AB}^{2,\text{null}})}{\sigma(G_{AB}^{2,\text{null}})}, \qquad P_{AB} = \frac{1 + \sum_{r=1}^{R} \mathbf{1}[G_{AB}^{2,(r)} \geq G_{AB}^{2,\text{obs}}]}{R+1}. \tag{3}$$

A large positive $Z_{AB}$ means that exact retention patterns are associated more strongly than expected from their marginal frequencies alone. In Eq. (3), the null mean and standard deviation are calculated from the $R$ randomized values, with the standard deviation taken as the sample standard deviation (denominator $R-1$).

The genome-wide statistic establishes whether two species share scar structure, but downstream reconstruction also needs to know which copy pairs contribute to it. Within the same strata, each candidate copy pair is therefore tested under a fixed-margin hypergeometric null. If $a$ and $b$ genes are retained in the two copies within a stratum containing $n$ genes, the number $X$ retained in both copies follows

$$\Pr(X = x) = \frac{\binom{a}{x}\binom{n-a}{b-x}}{\binom{n}{b}} \tag{4}$$

The corresponding number of matching retention states is

$$M = n - a - b + 2X \tag{5}$$

Stratum-specific distributions of $M$ are convolved, and positive copy-pair excesses are corrected within species pairs using Benjamini-Hochberg false discovery rate.

**Operationally, t**he genome-wide test asks whether two species share more retention/loss structure than expected, while the copy-level test identifies the retained copies contributing to that signal.

***HMM analysis of spatial copy correspondence.*** A scar statistic is genome-wide and does not identify which copy correspondences hold over particular chromosomal intervals or whether they change along the ancestral coordinate. A hidden Markov model (HMM) is used for this local problem[75].

Related HMM-based spatial phasing has been used in several genomic settings. POInT conditions alternative orthology assignments on neighboring syntenic loci[70,71], while PhyloNet-HMM couples hidden genomic states to a reticulate evolutionary model for introgression[76]. In polyploid genome analysis, Mitros et al.[77] used an HMM to detect local switches in subgenome identity from subgenome-specific 13-mer densities in *Miscanthus*, and this repeat-based strategy was subsequently generalized across allopolyploids[78]. Our HMM differs in its state space and emissions: hidden states are injective copy-to-copy mappings and the emissions summarize local fractionation correspondence rather than sequence-based genealogy or subgenome-specific repetitive content.

For a pair with $m_A \le m_B$, each resolved HMM state is an injective mapping $\pi: \{1, \dots, m_A\} \to \{1, \dots, m_B\}$, so no two copies of $A$ map to the same copy of $B$ within one resolved block.

The number of resolved states is

$$|\mathcal{S}_{AB}| = \frac{m_B!}{(m_B - m_A)!}, \tag{6}$$

plus one unresolved state $U$.

Thus the *D. capensis-Aldrovanda* comparison at $(6,8)$ has 20,160 resolved mappings plus $U$. Genes are ordered by the ancestral *Vitis vinifera* coordinate and divided into consecutive complete 100-gene blocks; a trailing remainder of fewer than 100 genes is discarded.

For block $b$, centered binary-retention correlation between copies $i$ and $j$ is

$$\phi_{bij} = \frac{(\mathbf{x}_{bi}^A - \bar{x}_{bi}^A)^T (\mathbf{x}_{bj}^B - \bar{x}_{bj}^B)}{\|\mathbf{x}_{bi}^A - \bar{x}_{bi}^A\| \, \|\mathbf{x}_{bj}^B - \bar{x}_{bj}^B\|}, \tag{7}$$

with 0 assigned when the denominator is zero.

A mapping score sums these edge similarities, $S_b^\phi(\pi) = \sum_i \phi_{bi,\pi(i)}$. A complementary centered-cosine score $S_b^r(\pi)$ compares copy-wise mean-retention profiles. For this score, one vector contains the blockwise mean retention rates of the A copies and the other the correspondingly ordered B-copy means under the candidate mapping; zero is assigned when the centered-cosine denominator is zero.

After both are standardized across resolved states within the block, the log-emission score (log potential) is

$$\begin{gathered} S_b^r(\pi) = \frac{(\mathbf{a}_b - \bar{a}_b \mathbf{1})^T (\mathbf{b}_{b,\pi} - \bar{b}_{b,\pi} \mathbf{1})}{\|\mathbf{a}_b - \bar{a}_b \mathbf{1}\| \, \|\mathbf{b}_{b,\pi} - \bar{b}_{b,\pi} \mathbf{1}\|}, \\ e_b(\pi) = z\left[S_b^\phi(\pi)\right] + \kappa z[S_b^r(\pi)], \quad \kappa = 1, \qquad \widetilde{p}_b(\pi) \propto \exp\{e_b(\pi)\}, \quad e_b(U) = 0. \end{gathered} \tag{8}$$

For the unresolved state, $e_b(U) = 0$. These log potentials are used directly in the forward-backward and Viterbi recursions; explicit state-wise emission normalization is unnecessary.

The transition model favors continuity without forcing it. A resolved state remains unchanged with probability 0.85, moves to $U$ with probability 0.10, and distributes the remaining 0.05 equally among alternative resolved states. $U$ remains unresolved with probability 0.90 and distributes 0.10 equally across resolved states. Initial probability is 0.5 on $U$ and 0.5 divided equally among resolved states. Forward-backward recursions yield posterior state probabilities; Viterbi decoding yields the maximum-score path[75,79].

**Example:** If one ancestral block is best explained by the local pairing A1↔B2 and A2↔B1, and neighboring blocks favor the same pairing, the HMM reinforces that correspondence. If a long run of later blocks instead favors A1↔B3, the decoded path can switch there, marking a change in local correspondence along the ancestral coordinate.

The HMM is not used as a standalone significance test; rather, it describes local correspondence conditional on the selected copy sets. A species pair that fails the global scar test cannot acquire a direct cross-species ancestry edge merely because its HMM finds a locally coherent mapping.

***From chromosome-specific copies to species-wide threads.*** FractBias copy ranks are local labels. “Copy 1” on one ancestral chromosome need not represent the same biological genomic component as “copy 1” on another. We therefore construct recurrent species-wide *threads*. Each local copy is represented by its scar and HMM relationships to copies in the other species; local FractBias rank itself is omitted from the feature vector in the principal analysis. For a species of depth $K$, each ancestral chromosome must assign its $K$ local copies one-to-one to $K$ global threads.

We begin from multiple initial assignments, calculate thread centroids, and reassign each ancestral chromosome to those centroids using the Hungarian algorithm[66]. Centroids and assignments are iterated to stability; the best multi-start solution defines the global threads. A thread is a recurrent cross-chromosome pattern of fractionation correspondence, not a chromosome-number-defined subgenome.

A conceptual precedent for the threading step is SCORPiOs, which uses an unduplicated outgroup as a proxy for ancestral gene order, identifies duplicated descendant segments, and explicitly “threads” those segments across WGD-derived genomes before matching them by local syntenic similarity[72]. One-to-one graph matching under sequence and genomic-context information provides a related algorithmic precedent[80]. Our assignment differs in object and scope: it does not thread genes or pre-defined WGD segments, but iterates chromosome-local scar/HMM relationship profiles to obtain latent species-wide fractionation components.

For global threads $i$ and $j$ in different species, three quantities are summarized: HMM support $\tilde{h}_{ij}$, recurrence across the 19 ancestral *Vitis* chromosomes $r_{ij}$, and positive copy-level scar support $\tilde{s}_{ij}$. HMM and scar support are normalized to the maximum within each species pair, whereas recurrence is expressed directly as the fraction of the 19 ancestral *Vitis* chromosomes carrying the correspondence.

Their composite correspondence is

$$q_{ij} = g_{AB}\left(0.50\tilde{h}_{ij} + 0.30r_{ij} + 0.20\tilde{s}_{ij}\right), \tag{9}$$

where $g_{AB} = 1$ when the genome-wide scar test for species pair $(A, B)$ has $P < 0.05$ and 0 otherwise.

The weighting places greatest emphasis on spatially coherent HMM support, followed by cross-chromosome recurrence and exact scar support.

***The optimality criterion: a globally coherent parental-lineage partition.*** The central inference is a partition of terminal threads into parental-lineage classes $P1, P2, \ldots$. The classes are not defined by thresholding individual edges. Because "same parental lineage" must be transitive, a strong $i$-$j$ relationship and a strong $j$-$k$ relationship can force $i$ and $k$ to be evaluated together. The optimization rewards strong cross-species evidence, penalizes unsupported forced mergers, and allows repeated ancestry within a species only at an explicit cost.

Let $x_{ij} = 1$ if threads $i$ and $j$ are assigned to the same parental class and 0 otherwise. For cross-species pairs, coassignment contributes $q_{ij} - \tau$, with $\tau = 0.30$. Thus, evidence above 0.30 is positive; evidence below 0.30 is negative. A pair failing the global scar gate has $q_{ij} = 0$ and therefore contributes $-\tau$ if a larger transitive class would force the pair together. Unsupported relationships are consequently not ignored.

Define

$$U = \sum_{\substack{i<j \\ s(i) \neq s(j)}} x_{ij}(q_{ij} - \tau), \qquad D = \sum_{\substack{i<j \\ s(i)=s(j)}} x_{ij}, \tag{10}$$

where $s(i)$ is the species containing thread $i$.

Same-species coassignment is permitted for every taxon, but every such pair receives the common within-species coassignment penalty $\lambda_D = 0.30$. Accordingly, $D$ is a within-species coassignment-pair count and a pairwise regularization term, not a direct count of biological duplication events.

The objective is

$$\boxed{\Psi = U - \lambda_D D} = \sum_{\substack{i<j \\ s(i) \neq s(j)}} x_{ij}(q_{ij} - \tau) - \lambda_D \sum_{\substack{i<j \\ s(i)=s(j)}} x_{ij}. \tag{11}$$

Partition consistency is enforced by triangle inequalities for every triple $(i, j, k)$, for example $x_{ij} + x_{jk} - x_{ik} \leq 1$ and its permutations. Parental components need not correspond to sampled extant taxa. Unsampled or extinct ("ghost") parental sources are allowed: singleton classes can represent ancestry contributed by an unsampled or extinct progenitor, or a component whose fractionation history cannot yet be matched confidently to another sampled genome. A singleton $P$ class therefore must not be interpreted as proof of a parent unique to the extant species in which it is observed.

Mathematically, the coassignment part of this optimization is a weighted clique-partition/correlation-clustering problem: pairwise rewards and penalties are optimized subject to the same transitivity-enforcing triangle inequalities used in clique partitioning[81] and the broader correlation-clustering literature[82]. Transitivity clustering provides a biological-data precedent for enforcing globally consistent clusters from pairwise similarities[83], while plant synteny-network approaches show the broader utility of converting many pairwise syntenic relationships into network structure[84]. Here, edge weights are constructed from scar, HMM and recurrence evidence, with an explicit symmetric penalty for repeated within-species membership.

Pairwise syntenic relationships have also been represented and clustered as networks[84], and formal methods exist for inferring allopolyploid phylogenetic networks from gene-tree or multilocus data[85,86]. The object optimized here is different: its nodes are fractionation threads and its classes are hypotheses of shared parental-component history, not reticulation nodes inferred directly in a species network.

**Optimization behavior:** The criterion is conservative about merging genomic components. A large class is favored only if its positive cross-species evidence outweighs both sub-threshold relationships created by transitivity and any repeated within-species class membership needed to form the class. The same $\tau$ and $\lambda_D$ apply to every taxon; there are no species-specific permissions.

For each set of operational depths, the calculation follows the same sequence: (1) obtain global scar gates; (2) infer local HMM mappings; (3) build global threads; (4) calculate all $q_{ij}$; (5) create all coassignment variables; (6) assign signed cross-species rewards and the symmetric within-species coassignment penalty; (7) enforce transitivity and solve the mixed-integer partition problem; and (8) report the optimal parental classes, $U$, $D$, $\Psi$, and each species' terminal composition. Alternative operational depths are compared only after this complete sequence has been rerun.

***Model selection and pairwise evidence.*** For *Aldrovanda*, the complete pipeline was rerun for $K = 2$ through 9. The maximum objective occurs at $K = 8$: $U = 11.0261$, $D = 1$, and $\Psi = 10.7261$. $K = 6$ is a close secondary solution ($\Delta\Psi = -0.1304$), while $K = 7$ and $K = 9$ are markedly poorer. The near-optimal ($K = 6$) solution differs from the selected ($K = 8$) network in its parental-class partition, including the *D. regia* fractionation thread coassigned with *Dionaea* D1. Because thread construction and parental-class optimization were repeated at each operational depth, the precise thread-level correspondence is conditional on the selected ($K = 8$) model rather than invariant across alternative resolutions. This selects the number of fractionation threads that best supports the seven-taxon network under the same criterion used to infer the parental classes; it does not infer octoploidy.

The pairwise evidence is strong overall (**Supplementary Note 3 Fig. 3**). Twenty of 21 global scar tests are significant at the resolution of 5,000 randomizations. The exception is *Dionaea-Ancistrocladus* ($Z = 0.57$, $P = 0.262$). The strongest scar association is *Ancistrocladus-Triphyophyllum* ($Z = 477.64$), followed by *Dionaea-Aldrovanda* ($Z = 63.03$). Mean posterior probability assigned to resolved HMM states ranges from 0.807 to 0.959, while Viterbi-resolved block fractions range from 0.947 to 1.000. At *Aldrovanda* $K = 8$, posterior resolution remains high against all six other taxa, showing that the additional depth is not represented mainly by unresolved HMM states.

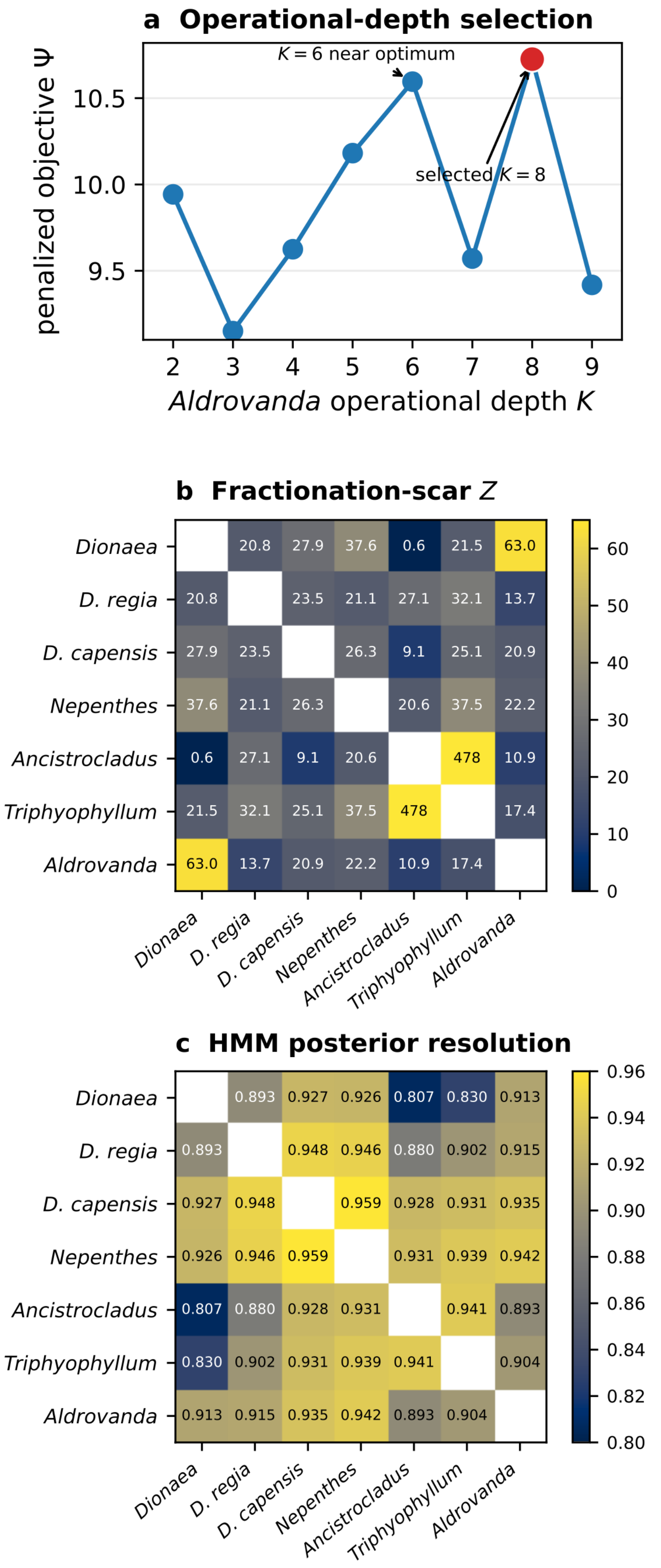
a Operational-depth selection
penalized objective Ψ
10.5
10.0
9.5
K = 6 near optimum
selected K = 8
2 3 4 5 6 7 8 9
Aldrovanda operational depth K
b Fractionation-scar Z
Dionaea
D. regia
D. capensis
Nepenthes
Ancistrocladus
Triphyophyllum
Aldrovanda
20.8 27.9 37.6 0.6 21.5 63.0
20.8 23.5 21.1 27.1 32.1 13.7
27.9 23.5 26.3 9.1 25.1 20.9
37.6 21.1 26.3 20.6 37.5 22.2
0.6 27.1 9.1 20.6 478 10.9
21.5 32.1 25.1 37.5 478 17.4
63.0 13.7 20.9 22.2 10.9 17.4
60 50 40 30 20 10 0
c HMM posterior resolution
0.893 0.927 0.926 0.807 0.830 0.913
0.893 0.948 0.946 0.880 0.902 0.915
0.927 0.948 0.959 0.928 0.931 0.935
0.926 0.946 0.959 0.931 0.939 0.942
0.807 0.880 0.928 0.931 0.941 0.893
0.830 0.902 0.931 0.939 0.941 0.904
0.913 0.915 0.935 0.942 0.893 0.904
0.96 0.94 0.92 0.90 0.88 0.86 0.84 0.82 0.80

**Supplementary Note 3 Fig. 3. Operational-depth selection and pairwise evidence.** (a) Penalized network objective across *Aldrovanda* $\boldsymbol{K} = \mathbf{2}$-9. (b) Genome-wide fractionation-scar $\boldsymbol{Z}$ statistics; the color scale is clipped for display because the *Ancistrocladus-Triphyophyllum* value is 477.64. (c) Mean posterior probability assigned to resolved HMM states.

Complete pairwise numerical output, including $G^2$ values, null means and standard deviations, HMM log scores, and Viterbi scores, is retained in the machine-readable tables global_generalized_scar_summary.tsv and hmm_pair_summary.tsv (see Zenodo archive at XXX); **Supplementary Note 3 Fig. 3** displays all 21 scar $Z$ values and all 21 posterior-resolution values.

***The optimized seven-taxon fractionation network.*** At $Aldrovanda\ K = 8$, the optimum contains ten parental-lineage classes. Four are broadly shared ($P1$-$P4$), $P5$ links two taxa, and $P6$-$P10$ are singleton/residual classes under the current sampling. The only within-species parental-class coassignment involves two $Aldrovanda$ threads, L3 and L5, both assigned to $P3$. The solution has $U = 11.0261$, $D = 1$, and $\Psi = 10.7261$.

Parental-class membership does not simply follow the species tree. *Ancistrocladus* and *Triphyophyllum* have identical $P1 + P2$ compositions and are connected by the strongest scar signal in the analysis. *Dionaea* is $P1 + P3$. *D. regia* is $P4 + P1 + P2$: only one of its three threads, R2, joins the *Dionaea* thread D1 in $P1$, whereas its other two threads belong to $P4$ and $P2$. *D. capensis* contains all three *D. regia* classes ($P1$, $P2$, $P4$) but its other three threads are $P3$, $P5$, and $P6$, rather than second copies of the three *D. regia* classes. *Nepenthes* resolves into five distinct classes ($P1$, $P2$, $P4$, $P9$, $P10$). *Aldrovanda* has eight threads assigned to seven classes and contains both *Dionaea muscipula* classes, $P1$ and $P3$, with $P3$ represented twice.

Several cross-taxon occupancy patterns are informative. $P1$ occurs in all seven scored taxa, while $P2$ occurs in six and is absent only from *Dionaea muscipula*. Their broad distributions are compatible with deep genomic components, although scars do not date them. Because parental-class assignments are transitive, the occurrence of P1 across all seven taxa does not imply significant direct fractionation-scar support between every pair. For example, *Dionaea* and *Ancistrocladus* both contribute a $P1$ thread despite their nonsignificant genome-wide pairwise scar comparison (Z = 0.571, empirical P = 0.262).

The sequence-based phylogenetic backbone projection sharpens the interpretation of the missing $P2$: with either of the exactly tied $P1$-root or $P2$-root solutions, $P2$ is reconstructed in the ancestor of *Dionaea* plus *Drosera* and then decreases to zero on the *Dionaea* branch. Thus, lineage-specific non-retention or loss is the most parsimonious reconciliation of the present terminal compositions; a replacement mechanism is not required by these data. The independent evidence that *Dionaea* is an allotetraploid and that snap-trap relationships are heterogeneous among gene trees remains relevant biological context, but it does not by itself identify what caused the $P2$ absence: $P1$ and $P2$ remain exactly tied as the possible single component at the carnivorous root, so their relative ancestral status is unresolved.

$P3$ is shared by *Dionaea*, *D. capensis*, and *Aldrovanda* and is represented twice in *Aldrovanda*; $P5$ is shared only by *D. capensis* and *Aldrovanda*. This combination makes a mosaic *Aldrovanda* history involving components related to both snap-trap and *Drosera*-side genomic pools plausible, but neither class identifies an extant parent, and ghost progenitors remain allowed. $P4$, shared by *D. regia*, *D. capensis*, *Nepenthes*, and *Aldrovanda*, is especially sensitive to the backbone placement of

*Nepenthes*. Because *Nepenthes* is sister to *Drosophyllum* plus (*Ancistrocladus*-*Triphyophyllum*), rather than lying on the branch leading to the snap-trap/*Drosera* clade, the parsimonious projection does not carry one uninterrupted $P4$ lineage vertically through these taxa; instead it requires separate $P4$ acquisitions for *Nepenthes* and for the *Aldrovanda*-*Dionaea*-*Drosera* side of the backbone. $P4$ is therefore better interpreted as a shared component affinity compatible with reticulation or unsampled/ghost ancestry than as straightforward evidence for continuous vertical inheritance from one sampled backbone ancestor. Main-text **Fig. 6c-e** nevertheless makes it intriguing, though not proven, that the $P4$-associated *Nepenthes* thread N3 could correspond to the dominant *Nepenthes* subgenomic signal; confirming that possibility would require an explicit thread-to-subgenome assignment.

**Supplementary Note 3 Fig. 4** is intentionally drawn as lanes rather than as a tree. Horizontal lines are equivalence classes created by the optimization, and are not directions of species descent. This keeps the primary result, i.e., which genomic threads belong to the same inferred parental class, separate from the later reconciliation of those classes with a bifurcating sequence-based tree.

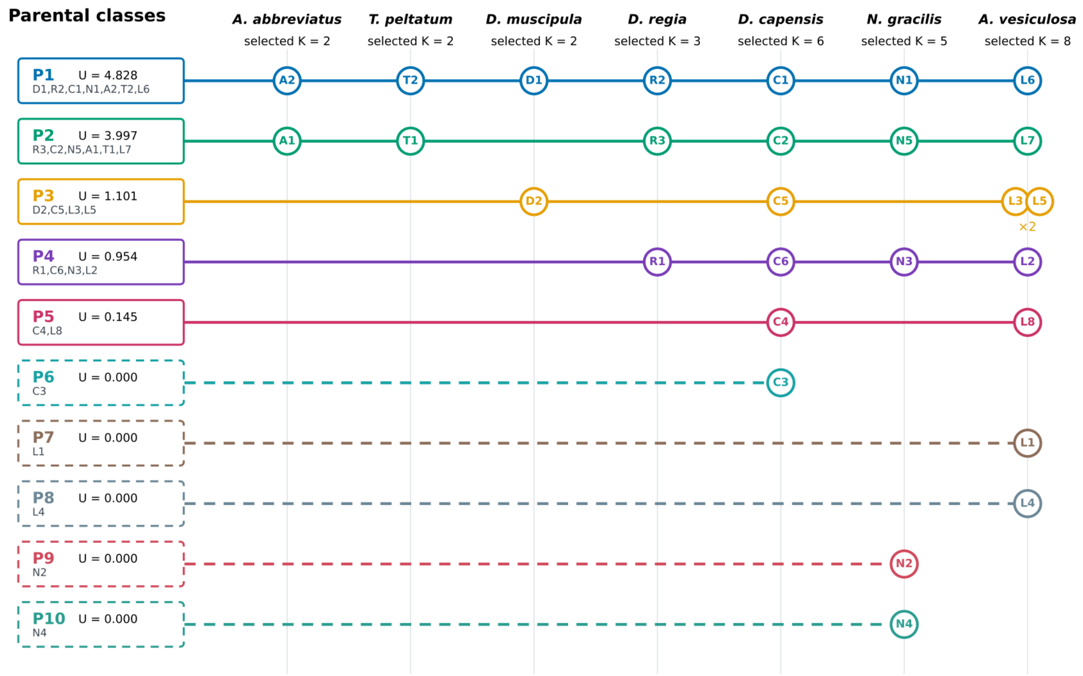


**Supplementary Note 3 Fig. 4**. **Tree-independent seven-taxon fractionation/HMM network at selected *Aldrovanda vesiculosa* $\boldsymbol{K = 8}$**. Horizontal colored lanes are the optimized transitive parental-lineage classes $\boldsymbol{P1}$-$\boldsymbol{P10}$, not branches of a species tree. Labeled circles show terminal thread assignments for each scored taxon. Dashed classes are singleton/ghost-compatible components under the present sampling. *Ancistrocladus* and *Triphyophyllum* have identical $\boldsymbol{P1 + P2}$ compositions; *Dionaea* is $\boldsymbol{P1 + P3}$; only one of the three *D. regia* fractionation threads joins a *Dionaea* class: R2 and Dionaea thread D1 are both assigned to $P1$. The repeated $P3$ assignment of Aldrovanda threads L3 and L5 is the sole within-species parental-class coassignment in the optimum. *Vitis* provides the common ancestral coordinate system but is not a terminal taxon in this tree-independent partition. The design of this diagram was assisted by ChatGPT (GPT-5.6 Sol); its scientific content and final presentation were reviewed by the authors.

***Synthesis across taxa and comparison with Supplementary Note 2 and Fig. 7.*** **Supplementary Note 3 Fig. 4** and **Supplementary Note 3 Table 2** summarize the terminal compositions. The comparison with **Supplementary Note 2** is strongest for *D. regia*. **Supplementary Note 2** identifies R2 as the clearest channel-specific structural affinity to *Dionaea*. Independently, the scar/HMM network places *D. regia* fractionation thread R2 in parental class $P1$ together with *Dionaea* thread

D1. However, $P1$ occurs in all seven sampled taxa and therefore does not represent a *Dionaea*-specific relationship. Moreover, the structural channels and fractionation threads were inferred from different features and have not been directly equated. The two analyses are compatible with shared ancestral genomic components, but the fractionation result does not independently establish the preferential *Dionaea* affinity detected in the structural analysis. The sharing of an ancestral genomic component between *D. regia* and *Dionaea* is compatible with **Fig. 7**, although the broad distribution of $P1$ across the sampled taxa does not independently support a preferential relationship between these two lineages.

This concordance does not distinguish **Fig. 7** models I and II. A shared fractionation history can arise because a genomic component entered *D. regia* through direct reticulation from a *Dionaea muscipula*-like lineage, as in model I, or because both lineages retained an older component from deeper shared ancestry, as in model II. Fractionation scars identify shared component history but do not supply direction or calendar time. The preferred model I in the main-text synthesis therefore remains a conclusion from the combined structural and phylogenomic evidence, while the present network independently supports shared ancestral component history without distinguishing a direct *Dionaea*-like contribution from deeper shared ancestry.

**Supplementary Note 3 Table 2. Synthetic interpretation of the seven scored taxa.** The fractionation-network column reports only what follows from the scar/HMM optimization. The final column places that result beside the structural interpretation developed in **Supplementary Note 2** and main-text **Fig. 7**.

| **Taxon** | **Terminal composition** | **Fractionation-network inference** | **Relationship to Supplementary Note 2 / Fig. 7** |
|---|---|---|---|
| *Ancistrocladus abbreviatus* | $P1 + P2$ | Same two parental classes as *Triphyophyllum*; pairwise scar $Z = 477.64$. | Strong independent support for a shared two-component genomic history. |
| *Triphyophyllum peltatum* | $P1 + P2$ | Same two parental classes as *Ancistrocladus*. | Concordant with their shared polyploid history in the **Fig. 7** synthesis. |
| *Dionaea muscipula* | $P1 + P3$ | Two distinct classes. $P1$ is universal; $P3$ is shared with *D. capensis* and *Aldrovanda*. $P2$, present in every other scored taxon, is absent. | The formal projection reconstructs $P2$ in the *Dionaea*+*Drosera* ancestor and a terminal decrease on the *Dionaea* branch, making lineage-specific non-retention/loss the simplest reconciliation. The independent allopolyploid evidence provides context but does not require a replacement mechanism. $P1$ supplies the class involved in the *D. regia* channel-specific affinity. |
| *Drosera regia* | $P4 + P1 + P2$ | Only one of the three *D. regia* fractionation threads joins a *Dionaea* class: R2 and *Dionaea* thread D1 are both assigned to $P1$. Because $P1$ is shared across all seven taxa, this correspondence does not independently establish preferential affinity to Dionaea. | This concordance is compatible with the one-component *Dionaea*-like pattern identified in **Supplementary Note 2**, where structural channel R2 showed the clearest preferential affinity to *Dionaea*. The two analyses use different features and do not assume that similarly named channels and threads are biologically identical. The result is compatible with both **Fig. 7** models I and II; fractionation scars alone do not determine donor direction or timing. |
| *Drosera capensis* | $P1 + P2 + P6 + P5 + P3 + P4$ | Contains all three *D. regia* classes, but the other three threads are three different classes rather than duplicate copies of the *D. regia* set. | Supports shared ancestral substrate while arguing against a simple retained doubling of the extant *D. regia*-like three-component architecture. |
| *Nepenthes gracilis* | $P1 + P9 + P4 + P10 + P2$ | Five distinct parental classes; $P4$ is also present in *D. regia*, *D. capensis*, and *Aldrovanda*. | On the sequence-based phylogenetic backbone used here, $P4$ in *Nepenthes* is reconciled independently from $P4$ on the snap-trap/*Drosera* side, favoring a reticulate or ghost-compatible interpretation over simple continuous vertical inheritance. It may still correspond to the dominant *Nepenthes* subgenomic signal highlighted in |

| Taxon | Terminal composition | Fractionation-network inference | Relationship to Supplementary Note 2 / Fig. 7 |
|---|---|---|---|
| | | | main-text **Fig. 6**, but that requires explicit thread-to-subgenome assignment. |
| *Aldrovanda vesiculosa* | $P7 + P4 + P3 + P8 + P3 + P1 + P2 + P5$ | Eight threads, seven classes; both *Dionaea* classes occur, *P*3 is represented by two distinct *Aldrovanda* fractionation threads, and *P*5 is shared only with *D. capensis*. | Supports a complex mosaic history with affinities to several genomic pools. A snap-trap/*Drosera*-side reticulate history is plausible, but the *P* classes do not identify extant parental species. Selected $K = 8$ is not a ploidy estimate. |

***Projecting the fixed network onto a bifurcating backbone.*** The network is the primary fractionation result. Projection is a separate reconciliation step that places inheritance, acquisition, duplication, and loss of the fixed terminal compositions parsimoniously on the independently inferred sequence-based phylogenetic backbone. This is necessary because a bifurcating tree cannot represent all parental histories of an allopolyploid without adding reticulation information. In the backbone used here, *Nepenthes* is sister to the clade comprising *Drosophyllum* and (*Ancistrocladus*, *Triphyophyllum*), whereas the *Drosera–Dionaea–Aldrovanda* clade forms the other sampled carnivorous branch. This placement of *Nepenthes* is also recovered in the protein-based ASTRAL reconstruction (**Supplementary Materials Fig. S6**). However, the selected projection backbone differs from that reconstruction in the internal relationships of *D. regia*, *Dionaea*, *Aldrovanda*, and the remaining *Drosera* species, consistent with the phylogenetic discordance documented across the sequence-based analyses. The projection is therefore conditional on the specified backbone, and inferred component acquisitions and losses may differ under alternative resolutions.

The projection is a phylogenetic reconciliation and ancestral-state optimization problem. Minimum-change assignment of internal states on a fixed tree is a classical problem[87,88], and copy-number methods such as CAFE similarly infer ancestral family sizes on a supplied phylogeny from extant copy counts[89]. More directly for polyploidy, GRAMPA reconciles gene trees with multiply labelled species trees to place polyploid events and parental lineages while minimizing duplication/loss discordance[90]. Reconciliation methods such as ecceTERA and RANGER-DTLx explicitly accommodate transfers involving unsampled or extinct lineages[91,92]. Synesth, and its later extension FullSynesth, move still closer in data type by reconciling syntenic units with a species tree under segmental duplication, loss, fission, transfer and unsampled-lineage events[93,94]. The projection here is narrower in input but different in direction: the terminal $P$-class multiplicities have already been fixed by the tree-independent fractionation analysis, the species-tree topology is not searched, and dynamic programming asks only how those fixed components can be reconciled with that backbone under the stated lexicographic event criterion.

For each parental lineage $P$ and backbone node $v$, the projection assigns copy number $n_{vP} \in \{0,1,\dots,n_P^{\max}\}$. Sampled terminal states are fixed exactly to the optimized network; backbone-only taxa such as *Drosophyllum* and *D. spatulata* are unobserved; they are not zero-copy observations. On a parent-child edge with states $a \to b$, a transition $0 \to b > 0$ means that the lineage enters that branch from outside the vertically inherited state; if more than one copy enters, the excess copies are counted separately as specified in Eq. (12). A transition $b > a > 0$ contributes copy increases according to the child-minus-parent difference, and $a > b$ contributes copy-number decreases according to the parent-minus-child difference.

Dynamic programming minimizes the lexicographic score

$$
\begin{gathered}
(R, D, G, L, O), \\
r(a,b) = \mathbf{1}[a = 0 < b], \qquad d(a,b) = \mathbf{1}[a = 0 < b](b-1) + \mathbf{1}[a > 0]\max(b-a, 0), \qquad \ell(a,b) = \max(a-b, 0), \\
R = \sum_{e,P} r(a_{eP}, b_{eP}), \qquad D = \sum_{e,P} d(a_{eP}, b_{eP}), \qquad L = \sum_{e,P} \ell(a_{eP}, b_{eP}), \\
G = \sum_{P} \mathbf{1}[\exists e : r(a_{eP}, b_{eP}) = 1], \qquad O = \sum_{v,P} n_{vP}.
\end{gathered} \tag{12}
$$

in that order.

Because the objective is lexicographic, an extra event in an earlier category cannot be offset by fewer events in a later category. $R$ is the total number of branches requiring acquisition of a previously absent parental lineage; biologically, such an acquisition can represent reticulation or ancestry entering from an unsampled/extinct source. $D$ is the total number of copy increases, including excess copies that accompany an acquisition into a branch when the child state exceeds one. $G$ counts how many *distinct* parental classes require at least one such external source anywhere in the reconstruction; this prevents repeated acquisitions of many unrelated ghost components from being treated as equivalent to repeated use of the same component. $L$ is the total number of copy-number decreases. Finally, $O = \sum_{v,P} n_{vP}$ is the summed reconstructed-state occupancy over all reconstructed nodes, internal and terminal, and parental classes; it is used only after $R$, $D$, $G$, and $L$ tie, favoring the solution that carries fewer unnecessary parental copies through reconstructed states. The total R, D, and L values are obtained by summing the corresponding edgewise costs in Eq. (12) over backbone branches and parental classes.

Ghost ancestry is explicit in the projection. A $P$ class may enter from an unsampled or extinct lineage without being assigned to any sampled donor taxon. With the sequence-based phylogenetic backbone used in **Supplementary Note 3 Fig. 5a**, the raw optimum is $R = 11$, $D = 1$, $G = 9$, $L = 5$, and $O = 62$. The carnivorous root is constrained to one parental-lineage copy. $P1$ and $P2$ are exactly tied under all five criteria, so $P1$ is drawn at the root only because of deterministic label ordering, not because the analysis establishes it as ancestral to $P2$.

After dynamic programming, the renderer applies two biological display rules. First, terminal non-contribution is shown by omission rather than by a terminal "loss" symbol; loss glyphs are reserved for explicit internal positive-to-zero changes. Second, if all scored descendants of a monophyletic clade have an identical terminal composition, one shared event is placed on their subtending stem. This collapses the identical $P1 + P2$ compositions of *Ancistrocladus* and *Triphyophyllum* to one shared stem event. The repeated $P3$ in *Aldrovanda* is shown as a copy increase.

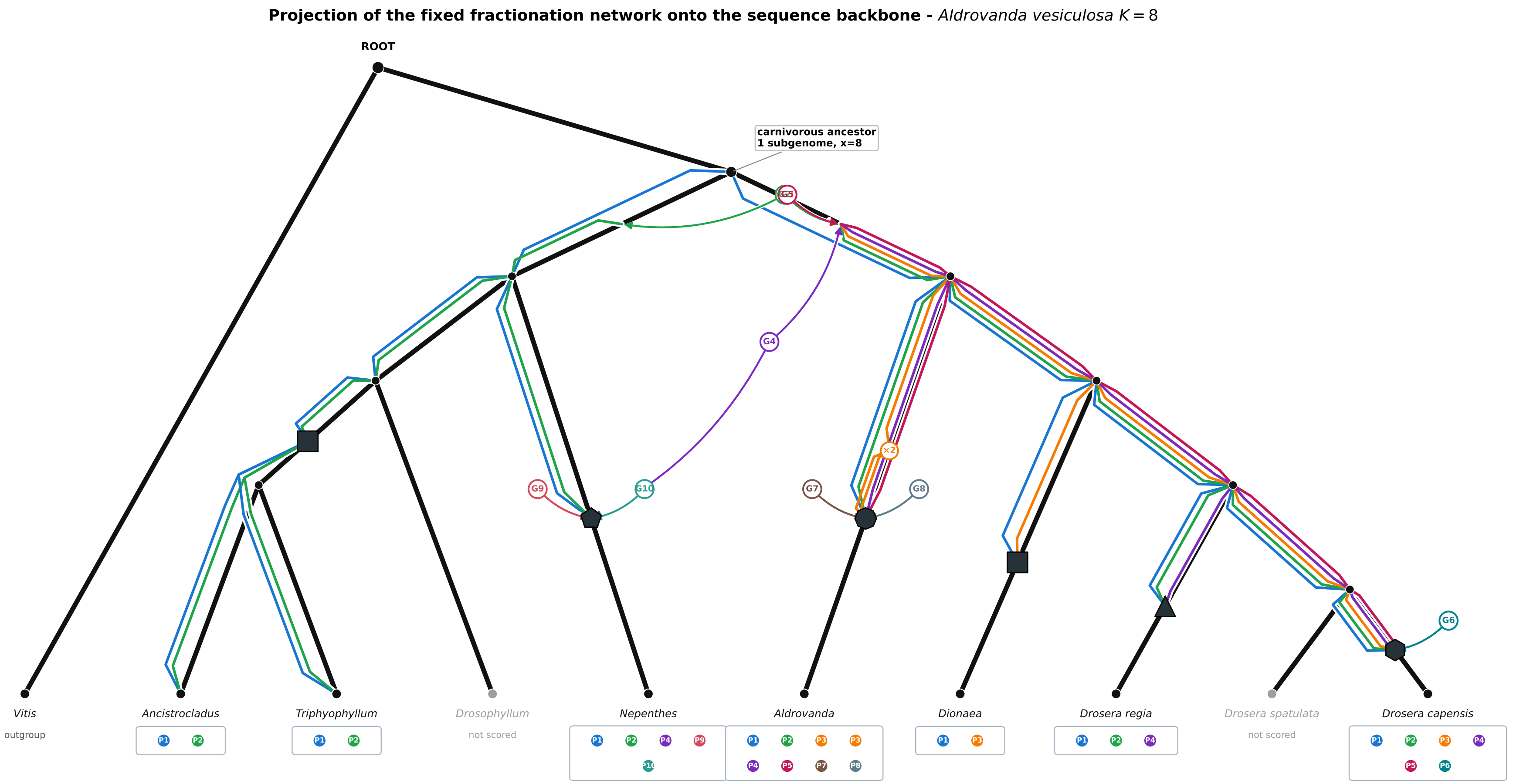


**Supplementary Note 3 Fig. 5. (a) Algorithmic projection of the fixed fractionation network onto the sequence-based phylogenetic backbone.** The backbone places *Nepenthes* sister to *Drosophyllum* plus (*Ancistrocladus*-*Triphyophyllum*). Black lines are the phylogenetic backbone; colored lanes are parental classes already fixed by the tree-independent optimization; dashed black terminal branches are backbone taxa not scored by the fractionation analysis; dashed colored arrows are ghost/reticulate inputs required by reconciliation; event symbols summarize terminal component number, and $\times\mathbf{2}$ marks the repeated $\boldsymbol{P3}$ contribution to *Aldrovanda*. The shared *Ancistrocladus*-*Triphyophyllum* event reflects their identical $\boldsymbol{P1} + \boldsymbol{P2}$ compositions. Acquisitions are reconciliation requirements, not direct observations of a particular donor event.

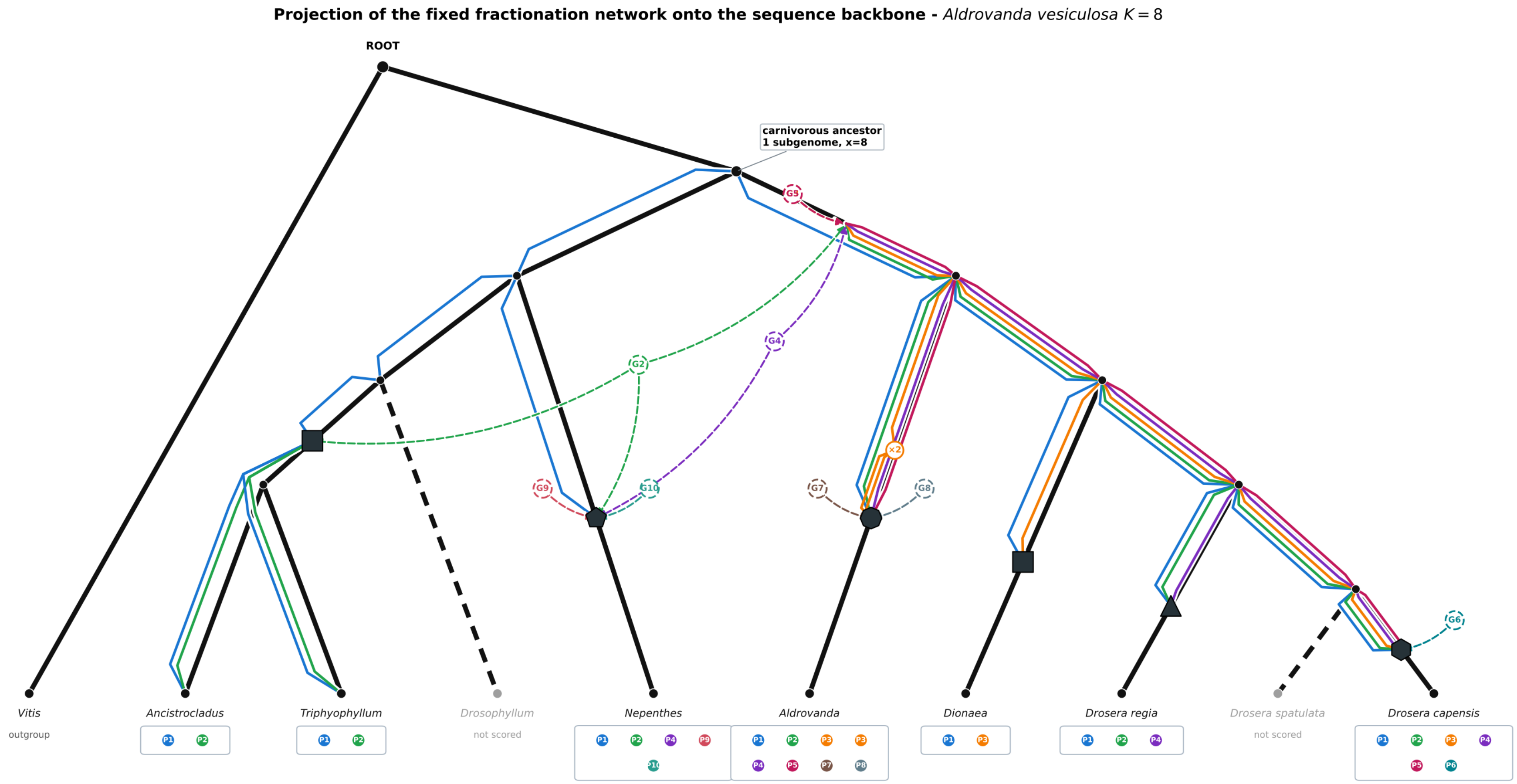


**Supplementary Note 3 Fig. 5, continued: (b) External-information-guided heuristic modification of the projection.** This panel is a biologically informed companion to the formal algorithmic projection in panel a. It preserves the same terminal $\boldsymbol{P}$ compositions but adjusts internal event placement to accommodate two independent considerations not encoded in the dynamic-programming criterion itself: (i) the *Ancistrocladus-Triphyophyllum* tetraploidy is treated as exclusive of *Drosophyllum*, consistent with an $\boldsymbol{n = 6}$ karyotype plausibly derived from an $\boldsymbol{x = 8}$ ancestor by two chromosome fusions, so $\boldsymbol{P2}$ is introduced only on the stem leading to *Ancistrocladus* plus *Triphyophyllum* after the *Drosophyllum* split; and (ii) only one *Nepenthes* component is carried vertically into the *Nepenthes* branch, whereas $P2$, $P4$, $P9$, and $P10$ are represented as separate inputs along that branch rather than as uninterrupted vertical inheritance from an ancestor shared with the *Ancistrocladus–Triphyophyllum* lineage. Panel **b** is therefore interpretive and hypothesis-guided, whereas panel **a** remains the fully algorithmic projection. The design of this FractLineage output was assisted by ChatGPT (GPT-5.6 Sol); its scientific content and final presentation were reviewed by the authors.

***Scope and limitations.*** The method uses historical information complementary to sequence substitution and chromosome-number evidence. It addresses three questions: whether genomes share non-random exact retention patterns; whether particular copies correspond coherently along ancestral chromosomes; and whether those pairwise signals can be assembled into one transitive multi-taxon parental partition without excessive unsupported merging or within-species duplication. The result is a network of genomic components rather than another species tree.

The signal has clear limitations. A fractionation scar does not date the event that created it, identify an extinct donor, or by itself distinguish direct hybridization from retention of deeper ancestral structure. Operational $K$ is not ploidy. Singleton classes are explicitly compatible with unsampled or extinct ghost ancestry and are not literal one-species parents. The projection is not a second inference of terminal ancestry: its terminal states are fixed before the tree is introduced. Its role is to show where a reticulate component network is compatible or incompatible with a bifurcating backbone and which extra acquisitions or copy-number changes are required to reconcile them.

Within these limits, the analysis provides an independent counterpart to the schematic logic of **Fig. 7**. It recovers a single *Dionaea*-associated component within *D. regia*, supports substantial but non-identical ancestral substrate between *D. regia* and *D. capensis*, strongly unites *Ancistrocladus* and *Triphyophyllum*, recovers five *Nepenthes* components, and identifies a deeper *Aldrovanda* composition containing both *Dionaea* classes. These results refine the synthesis without assigning temporal or directional information that the fractionation signal cannot provide.

***Data requirements and transferable implementation.*** The workflow is defined by explicit data objects rather than taxon-specific drawing rules. The minimal upstream requirement is a common ancestral coordinate system with syntenic retained-copy calls for each sampled taxon. In the present analysis, the principal gene-level files are FractBias fractbias-genes.csv and fractbias-results.csv outputs, from which the selected local copies are converted to binary retention matrices. Global scar tests use deterministic random seeds and 5,000 fixed-margin randomizations. HMMs use 100-gene blocks, all injective mappings, the unresolved state and transition scheme stated above, and $\kappa = 1$. Threads are built from scar/HMM relationship features without local rank. The parental partition uses Eq. 11 with $\tau = 0.30$ and $\lambda_D = 0.30$ in every taxon.

A different clade requires (i) syntenic retained-copy matrices indexed by a shared ancestral coordinate, (ii) a candidate operational depth $K$ for each taxon, and, only for the final reconciliation, (iii) an independently selected bifurcating backbone in Newick format. The common coordinate need not be *Vitis*, as was used here: it can be any sufficiently conserved reference or reconstructed ancestral-block system that permits homologous retained copies to be compared at the same ancestral loci. Similarly, the method does not require exactly seven sampled taxa. Once the retention matrices are supplied, the scar null, HMM state definition, chromosome-wise threading, transitive optimization, and within-species coassignment penalty are unchanged. For projection, the supplied tree can be the best-supported sequence topology for the focal group, but the projection does not search or alter that topology; it overlays the already inferred component network onto the tree and reports the extra reticulation or copy-number events required for compatibility.

Again, the closest precedent pipelines cover individual parts of the workflow rather than the full sequence. POInT and SCORPiOs are the strongest antecedents for extracting historical information from post-WGD retention/loss and conserved synteny; POInT and genomic ancestry HMMs provide direct precedents for spatial state phasing; clique partitioning and correlation clustering supply the mathematical basis for the transitive component partition; and GRAMPA, Synesth and FullSynesth are the closest downstream analogues for reconciling polyploid or syntenic histories to a species tree. We are not aware of a prior implementation combining the same stages and data objects in this order: an exact-pattern fractionation test, injective-copy HMM, cross-chromosome thread construction, tree-independent global parental-class optimization, and only then fixed-state projection to an external bifurcating backbone.

### Supplementary References


1. Murashige, T. & Skoog, F. A revised medium for rapid growth and bio assays with tobacco tissue cultures. *Physiologia plantarum* **15**(1962).
2. Steinmüller, K. & Apel, K. A simple and efficient procedure for isolating plant chromatin which is suitable for studies of DNase I-sensitive domains and hypersensitive sites. *Plant molecular biology* **7**, 87-94 (1986).
3. Hatano, S., Yamaguchi, J. & Hirai, A. The preparation of high-molecular-weight DNA from rice and its analysis by pulsed-field gel electrophoresis. *Plant Science* **83**, 55-64 (1992).
4. Cairns, J. *et al.* CHiCAGO: robust detection of DNA looping interactions in Capture Hi-C data. *Genome biology* **17**, 127 (2016).
5. Putnam, N.H. *et al.* Chromosome-scale shotgun assembly using an in vitro method for long-range linkage. *Genome research* **26**, 342 (2016).
6. Koren, S. *et al.* Canu: scalable and accurate long-read assembly via adaptive k-mer weighting and repeat separation. *Genome Res* **27**, 722-736 (2017).
7. Zimin, A.V. *et al.* The MaSuRCA genome assembler. *Bioinformatics* **29**, 2669-2677 (2013).
8. Zimin, A.V. *et al.* Hybrid assembly of the large and highly repetitive genome of Aegilops tauschii, a progenitor of bread wheat, with the MaSuRCA mega-reads algorithm. *Genome research* **27**, 787-792 (2017).
9. Chikhi, R. & Medvedev, P. Informed and automated k-mer size selection for genome assembly. *Bioinformatics* **30**, 31-37 (2014).
10. Bolger, A.M., Lohse, M. & Usadel, B. Trimmomatic: a flexible trimmer for Illumina sequence data. *Bioinformatics* **30**, 2114-2120 (2014).
11. Manni, M., Berkeley, M.R., Seppey, M. & Zdobnov, E.M. BUSCO: assessing genomic data quality and beyond. *Current Protocols* **1**, e323 (2021).
12. Gurevich, A., Saveliev, V., Vyahhi, N. & Tesler, G. QUAST: quality assessment tool for genome assemblies. *Bioinformatics* **29**, 1072-1075 (2013).
13. Robertson, G. *et al.* De novo assembly and analysis of RNA-seq data. *Nature methods* **7**, 909-912 (2010).
14. Haas, B.J. *et al.* De novo transcript sequence reconstruction from RNA-seq using the Trinity platform for reference generation and analysis. *Nature protocols* **8**, 1494-1512 (2013).
15. Kim, D., Paggi, J.M., Park, C., Bennett, C. & Salzberg, S.L. Graph-based genome alignment and genotyping with HISAT2 and HISAT-genotype. *Nature biotechnology* **37**, 907-915 (2019).
16. Pertea, M. *et al.* StringTie enables improved reconstruction of a transcriptome from RNA-seq reads. *Nature biotechnology* **33**, 290-295 (2015).
17. Gilbert, D. Gene-omes built from mRNA-seq not genome DNA. (2013).
18. Flynn, J.M. *et al.* RepeatModeler2 for automated genomic discovery of transposable element families. *Proceedings of the National Academy of Sciences* **117**, 9451-9457 (2020).
19. Chen, N. Using Repeat Masker to identify repetitive elements in genomic sequences. *Current protocols in bioinformatics* **5**, 4.10. 1-4.10. 14 (2004).
20. Haas, B.J. *et al.* Automated eukaryotic gene structure annotation using EVidenceModeler and the Program to Assemble Spliced Alignments. *Genome biology* **9**, R7 (2008).

21. Hoff, K.J., Lange, S., Lomsadze, A., Borodovsky, M. & Stanke, M. BRAKER1: unsupervised RNA-Seq-based genome annotation with GeneMark-ET and AUGUSTUS. *Bioinformatics* **32**, 767-769 (2016).
22. Stanke, M. *et al.* AUGUSTUS: ab initio prediction of alternative transcripts. *Nucleic acids research* **34**, W435-W439 (2006).
23. Lomsadze, A., Ter-Hovhannisyan, V., Chernoff, Y.O. & Borodovsky, M. Gene identification in novel eukaryotic genomes by self-training algorithm. *Nucleic Acids Research* **33**, 6494-6506 (2005).
24. Slater, G.S.C. & Birney, E. Automated generation of heuristics for biological sequence comparison. *BMC bioinformatics* **6**, 31 (2005).
25. Lovell, J.T. *et al.* GENESPACE tracks regions of interest and gene copy number variation across multiple genomes. *elife* **11**, e78526 (2022).
26. Albert, V.A. & Krabbenhoft, T.J. Navigating the CoGe Online Software Suite for Polyploidy Research. in *Polyploidy: Methods and Protocols* (ed. Van de Peer, Y.) 19-45 (Springer US, New York, NY, 2023).
27. Joyce, B.L. *et al.* FractBias: a graphical tool for assessing fractionation bias following polyploidy. *Bioinformatics* **33**, 552-554 (2016).
28. Hofstatter, P.G. *et al.* Repeat-based holocentromeres influence genome architecture and karyotype evolution. *Cell* (2022).
29. Jia, K.H. *et al.* SubPhaser: a robust allopolyploid subgenome phasing method based on subgenome-specific k-mers. *New Phytologist* **235**, 801-809 (2022).
30. Wickham, H. *et al.* Welcome to the Tidyverse. *Journal of open source software* **4**, 1686 (2019).
31. Hadley, W. *Ggplot2: Elegrant graphics for data analysis*, (Springer, 2016).
32. Neuwirth, E. & Neuwirth, M.E. Package 'RColorBrewer'. *ColorBrewer Palettes* (2014).
33. Wilke, C.O. Ridgeline Plots in 'ggplot2'[R Package Ggridges Version 0.5. 3]. *January. https://cran. r-project. org/web/packages/ggridges/index. html* (2021).
34. Aphalo, P. ggpmisc: Miscellaneous Extensions to "ggplot2"(R package version 0.3. 6). (2020).
35. Sensalari, C., Maere, S. & Lohaus, R. ksrates: positioning whole-genome duplications relative to speciation events in KS distributions. *Bioinformatics* **38**, 530-532 (2022).
36. Dainat, J. AGAT: Another Gff Analysis Toolkit to handle annotations in any GTF/GFF format (version 1.4.0). (Zenodo, 2022).
37. Saul, F. *et al.* Subgenome dominance shapes novel gene evolution in the decaploid pitcher plant Nepenthes gracilis. *Nature plants* **9**, 2000-2015 (2023).
38. Denoeud, F. *et al.* The coffee genome provides insight into the convergent evolution of caffeine biosynthesis. *science* **345**, 1181-1184 (2014).
39. Emms, D.M. & Kelly, S. OrthoFinder: phylogenetic orthology inference for comparative genomics. *Genome biology* **20**, 1-14 (2019).
40. Emms, D.M. & Kelly, S. STAG: Species tree inference from all genes. Preprint at https://doi.org/10.1101/267914. (2018).
41. Emms, D.M. & Kelly, S. STRIDE: species tree root inference from gene duplication events. *Molecular biology and evolution* **34**, 3267-3278 (2017).
42. Novák, P., Hoštáková, N., Neumann, P. & Macas, J. DANTE and DANTE_LTR: lineage-centric annotation pipelines for long terminal repeat retrotransposons in plant genomes. *NAR Genomics and Bioinformatics* **6**(2024).

43. Neumann, P., Novák, P., Hoštáková, N. & Macas, J. Systematic survey of plant LTR-retrotransposons elucidates phylogenetic relationships of their polyprotein domains and provides a reference for element classification. *Mobile DNA* **10**, 1 (2019).
44. Novák, P., Neumann, P. & Macas, J. Global analysis of repetitive DNA from unassembled sequence reads using RepeatExplorer2. *Nature Protocols* **15**, 3745-3776 (2020).
45. Altschul, S.F., Gish, W., Miller, W., Myers, E.W. & Lipman, D.J. Basic local alignment search tool. *Journal of molecular biology* **215**, 403-410 (1990).
46. Yu, Y., Ouyang, Y. & Yao, W. shinyCircos: an R/Shiny application for interactive creation of Circos plot. *Bioinformatics* **34**, 1229-1231 (2017).
47. Fan, H., Ives, A.R., Surget-Groba, Y. & Cannon, C.H. An assembly and alignment-free method of phylogeny reconstruction from next-generation sequencing data. *BMC genomics* **16**, 522 (2015).
48. Oliveira, L. *et al.* KNL1 and NDC80 represent new universal markers for the detection of functional centromeres in plants. *Chromosome Research* **32**, 3 (2024).
49. Macas, J., Neumann, P. & Navrátilová, A. Repetitive DNA in the pea (Pisum sativum L.) genome: comprehensive characterization using 454 sequencing and comparison to soybean and Medicago truncatula. *BMC Genomics* **8**, 427 (2007).
50. Katoh, K. & Standley, D.M. MAFFT Multiple Sequence Alignment Software Version 7: Improvements in Performance and Usability. *Molecular Biology and Evolution* **30**, 772-780 (2013).
51. Price, M.N., Dehal, P.S. & Arkin, A.P. FastTree 2–approximately maximum-likelihood trees for large alignments. *PloS one* **5**, e9490 (2010).
52. Letunic, I. & Bork, P. Interactive Tree of Life (iTOL) v6: recent updates to the phylogenetic tree display and annotation tool. *Nucleic acids research* **52**, W78-W82 (2024).
53. Paradis, E. & Schliep, K. ape 5.0: an environment for modern phylogenetics and evolutionary analyses in R. *Bioinformatics* **35**, 526-528 (2018).
54. Wickham, H. dplyr: A grammar of data manipulation. *R package version 04.* **3**, p156 (2015).
55. Montero, H., Freund, M. & Fukushima, K. Convergent losses of arbuscular mycorrhizal symbiosis in carnivorous plants. *bioRxiv*, 2025.04. 03.646726 (2025).
56. Mirarab, S. *et al.* ASTRAL: genome-scale coalescent-based species tree estimation. *Bioinformatics* **30**, i541-i548 (2014).
57. Wong, T.K. *et al.* IQ-TREE 3: Phylogenomic Inference Software using Complex Evolutionary Models. (2025).
58. Veleba, A. *et al.* Evolution of genome size and genomic GC content in carnivorous holokinetics (Droseraceae). *Ann Bot* **119**, 409-416 (2017).
59. Robinson, J.T. *et al.* Integrative genomics viewer. *Nat Biotechnol* **29**, 24-6 (2011).
60. Jaccard, P. Étude comparative de la distribution florale dans une portion des Alpes et des Jura. *Bull Soc Vaudoise Sci Nat* **37**, 547-579 (1901).
61. Ward Jr, J.H. Hierarchical grouping to optimize an objective function. *Journal of the American statistical association* **58**, 236-244 (1963).
62. Salton, G., Wong, A. & Yang, C.-S. A vector space model for automatic indexing. *Communications of the ACM* **18**, 613-620 (1975).
63. Shannon, C.E. A mathematical theory of communication. *The Bell system technical journal* **27**, 379-423 (1948).

64. Pielou, E.C. The measurement of diversity in different types of biological collections. *Journal of theoretical biology* **13**, 131-144 (1966).
65. Cracau, D. & Lima, J.E.D. On the normalized herfindahl-hirschman index: a technical note. *International Journal on Food System Dynamics* **7**, 382-386 (2016).
66. Kuhn, H.W. The Hungarian method for the assignment problem. *Naval research logistics quarterly* **2**, 83-97 (1955).
67. Stull, G.W., Pham, K.K., Soltis, P.S. & Soltis, D.E. Deep reticulation: the long legacy of hybridization in vascular plant evolution. *The Plant Journal* **114**, 743-766 (2023).
68. Scannell, D.R., Byrne, K.P., Gordon, J.L., Wong, S. & Wolfe, K.H. Multiple rounds of speciation associated with reciprocal gene loss in polyploid yeasts. *Nature* **440**, 341-345 (2006).
69. Scannell, D.R. *et al.* Independent sorting-out of thousands of duplicated gene pairs in two yeast species descended from a whole-genome duplication. *Proceedings of the National Academy of Sciences* **104**, 8397-8402 (2007).
70. Conant, G.C. & Wolfe, K.H. Probabilistic cross-species inference of orthologous genomic regions created by whole-genome duplication in yeast. *Genetics* **179**, 1681-1692 (2008).
71. Hao, Y. *et al.* Convergent evolution of polyploid genomes from across the eukaryotic tree of life. *G3* **12**, jkac094 (2022).
72. Parey, E. *et al.* Synteny-guided resolution of gene trees clarifies the functional impact of whole-genome duplications. *Molecular biology and evolution* **37**, 3324-3337 (2020).
73. Rokas, A. & Holland, P.W. Rare genomic changes as a tool for phylogenetics. *Trends in ecology & evolution* **15**, 454-459 (2000).
74. Patefield, W.M. Algorithm AS 159: an efficient method of generating random R× C tables with given row and column totals. *Journal of the Royal Statistical Society. Series C (Applied Statistics)* **30**, 91-97 (1981).
75. Rabiner, L.R. A tutorial on hidden Markov models and selected applications in speech recognition. *Proceedings of the IEEE* **77**, 257-286 (1989).
76. Liu, K.J. *et al.* An HMM-based comparative genomic framework for detecting introgression in eukaryotes. *PLoS computational biology* **10**, e1003649 (2014).
77. Mitros, T. *et al.* Genome biology of the paleotetraploid perennial biomass crop Miscanthus. *Nature communications* **11**, 5442 (2020).
78. Session, A. & Rokhsar, D. Transposon signatures of allopolyploid genome evolution. Nat. Commun. 14, 3180. (2023).
79. Viterbi, A. Error bounds for convolutional codes and an asymptotically optimum decoding algorithm. *IEEE transactions on Information Theory* **13**, 260-269 (1967).
80. Mahmood, K., Webb, G.I., Song, J., Whisstock, J.C. & Konagurthu, A.S. Efficient large-scale protein sequence comparison and gene matching to identify orthologs and co-orthologs. *Nucleic acids research* **40**, e44-e44 (2012).
81. Grötschel, M. & Wakabayashi, Y. A cutting plane algorithm for a clustering problem. *Mathematical Programming* **45**, 59-96 (1989).
82. Bansal, N., Blum, A. & Chawla, S. Correlation clustering. *Machine learning* **56**, 89-113 (2004).
83. Wittkop, T. *et al.* Partitioning biological data with transitivity clustering. *Nature methods* **7**, 419-420 (2010).
84. Zhao, T. & Schranz, M.E. Network approaches for plant phylogenomic synteny analysis. *Current opinion in plant biology* **36**, 129-134 (2017).

85. Yan, Z., Cao, Z., Liu, Y., Ogilvie, H.A. & Nakhleh, L. Maximum parsimony inference of phylogenetic networks in the presence of polyploid complexes. *Systematic Biology* **71**, 706-720 (2022).
86. Jones, G., Sagitov, S. & Oxelman, B. Statistical inference of allopolyploid species networks in the presence of incomplete lineage sorting. *Systematic biology* **62**, 467-478 (2013).
87. Fitch, W.M. Toward defining the course of evolution: minimum change for a specific tree topology. *Systematic Biology* **20**, 406-416 (1971).
88. Sankoff, D. Minimal mutation trees of sequences. *SIAM Journal on Applied Mathematics* **28**, 35-42 (1975).
89. De Bie, T., Cristianini, N., Demuth, J.P. & Hahn, M.W. CAFE: a computational tool for the study of gene family evolution. *Bioinformatics* **22**, 1269-1271 (2006).
90. Thomas, G.W., Ather, S.H. & Hahn, M.W. Gene-tree reconciliation with MUL-trees to resolve polyploidy events. *Systematic biology* **66**, 1007-1018 (2017).
91. Jacox, E., Chauve, C., Szöllősi, G.J., Ponty, Y. & Scornavacca, C. ecceTERA: comprehensive gene tree-species tree reconciliation using parsimony. *Bioinformatics* **32**, 2056-2058 (2016).
92. Weiner, S. & Bansal, M.S. Improved duplication-transfer-loss reconciliation with extinct and unsampled lineages. *Algorithms* **14**, 231 (2021).
93. Delabre, M. & El-Mabrouk, N. Synesth: comprehensive syntenic reconciliation with unsampled lineages. *Algorithms* **17**, 186 (2024).
94. Gascon, M., Delabre, M. & El-Mabrouk, N. FullSynesth: syntenic reconciliation of a set of consistent gene trees. *Theory of Computing Systems* **70**, 8 (2026).
